# Early mechanisms of aortic failure in a zebrafish model for thoracic aortic dissection and rupture

**DOI:** 10.1101/2024.02.12.580022

**Authors:** Michiel Vanhooydonck, Maxim Verlee, Marta Santana Silva, Lore Pottie, Annekatrien Boel, Matthias Van Impe, Hanna De Saffel, Lisa Caboor, Piyanoot Tapaneeyaphan, Anne Bonnin, Patrick Segers, Adelbert De Clercq, Andy Willaert, Delfien Syx, Patrick Sips, Bert Callewaert

**Affiliations:** Center for Medical Genetics Ghent (CMGG), Department of Biomolecular Medicine, Ghent University, 9000 Ghent, Belgium; Ghent-Fertility and Stem Cell Team (G-FaST), Department for Reproductive Medicine, Ghent University, 9000 Ghent, Belgium; Biophysical Models for Medical Applications (bioMMeda), Institute of Biomedical Engineering and Technology (IBiTech), Ghent University, 9000 Ghent, Belgium; Paul Scherrer Institut, Swiss Light Source, 5232 Villigen PSI, Switzerland; Evolutionary Developmental Biology, Biology Department, Ghent University, 9000 Ghent, Belgium

## Abstract

Thoracic aortic aneurysm and dissection (TAAD) associates with a high mortality rate. Despite the existence of different mouse models for TAAD, the underlying disease mechanisms remain elusive. Treatment options are limited and mainly consist of surgical repair at critical aortic diameters as current pharmacological interventions are unable to stop disease progression.

In humans, loss of function (LOF) of *SMAD3* and *SMAD6* impairs vascular homeostasis, increasing the risk for TAAD. We developed a zebrafish model for thoracic aortic dissection/rupture by targeting both ohnologs of *smad3* and *smad6*. At 10 days post fertilization, we found an increased diameter of the ventral aorta in *smad3a*^−/−^;*smad3b*^−/−^ double knockout zebrafish, while *smad6a*^−/−^;*smad6b*^−/−^ double knockout zebrafish have a reduced aortic diameter associated with early mortality. We discovered that a *smad3a*^−/−^;*smad3b*^−/−^;*smad6a*^−/−^;*smad6b*^−/−^ quadruple knockout (qKO) zebrafish model is viable and survives to adulthood, although exposure to stress leads to sudden death. Histological analysis of the adult ventral aorta shows medial elastolysis, aortic dissections and ruptures at sites exposed to high biomechanical stress. RNA-sequencing of 5 days post fertilization qKO zebrafish indicates a profile of reduced negative regulation of proteolysis and upregulation of melanogenesis, a previously unaddressed pathway in this pathology. We confirm that pharmacological modulation of tyrosinase, the enzyme responsible for the production of melanin, influences aortic morphology.

Overall, the qKO mutant, thus far the only known zebrafish model of thoracic aortic dissection and rupture, reveals novel SMAD3/6-dependent pathways that impact thoracic aortic homeostasis, in this way opening avenues for the development of novel treatments in TAAD.

## INTRODUCTION

Thoracic aortic dissection (TAD) affects 3-4 individuals per 100,000 people per year^1^. This life-threatening event is often, but not exclusively, preceded by thoracic aortic aneurysm (TAA). The disease mechanisms are incompletely understood, but evidence from genetic disorders implicates altered cell-matrix interactions (by impaired hemodynamic sensing or impaired matrix assembly), as well as aberrant transforming growth factor β (TGF-β) signaling^2–5^. Despite several mouse models for TAD^6^, the initial molecular mechanisms remain obscure.

Members of the TGF-β superfamily include TGF-β proteins, bone morphogenetic proteins (BMPs), activins, and growth and differentiation factors (GDFs). Binding of TGF-β to the tetrameric TGFβ-receptor complex results in phosphorylation of the receptor-regulated (R-) SMAD2 and −3 proteins. Binding of BMP and GDF to the tetrameric BMP-receptor complex results in phosphorylation of R-SMAD1, −5, and −8. Phosphorylated R-SMAD proteins recruit the common mediator (Co-) SMAD4 to form a heterotrimeric complex and translocate to the nucleus to activate the transcription of the TGF-β or BMP target genes^7^. The TGF-β and BMP signaling is negatively regulated by the inhibitory (I-) SMAD6 and - 7 proteins. I-SMAD6 primarily suppresses the pathway activated by the BMP-receptor, whereas I-SMAD7 inhibits both the TGF-β and BMP induced pathways^8^.

Pathogenic variants in *SMAD3* cause Loeys-Dietz syndrome type 3 (LDS3). LDS3 typically presents with early-onset osteo-arthritis, and arterial aneurysm and dissection, mainly affecting the ascending aorta. Other features include arterial tortuosity, mitral valve prolapse and craniofacial characteristics^2,9^. Pathogenic variants in *SMAD6* are associated with craniosynostosis^10^, radio-ulnar synostosis^11^ or congenital heart disease including bicuspid aortic valve^12^ with or without TAAD^13^ and Shone complex with a hypoplastic ascending aorta and arch^14^. Biallelic variants in *SMAD6* underly more complex cardiovascular phenotypes^15^. Interestingly, identical pathogenic variants in *SMAD6* may result in either cardiovascular anomalies, craniosynostosis or radioulnar synostosis^13^. Because *SMAD3* and *SMAD6* have been in linkage disequilibrium throughout evolution from teleost fish to human, it has been postulated that *SMAD3* could have a modifier role on *SMAD6* deficiency^16^. In line with this observation, we identified a 1.5 Mb 15q22.31-15q23 microdeletion encompassing *SMAD3* and *SMAD6* in an 11-year-old-boy and his 33-year-old mother, both presenting with bicuspid aortic valve and arterial tortuosity, but no aneurysms, along with skeletal and cutaneous anomalies of LDS3, hinting that deletion of *SMAD6* may influence the phenotype caused by *SMAD3* LOF (**Supplementary Case Presentation**^17^). Due to the teleost specific genome duplication, zebrafish have two ohnolog copies for both *SMAD3* and *SMAD6*^18^. We established zebrafish models deficient for each of these ohnologs identified as *smad3a* and *smad3b,* and *smad6a* and *smad6b*, respectively. The human SMAD3 (ENSP00000332973) protein has a sequence similarity of 97.17% with zebrafish Smad3a (ENSDARP00000045373) and 93.4% with zebrafish Smad3b (ENSDARP00000043454). SMAD6 (ENSP00000288840) is less conserved with a similarity of 53.3% and 62.38% with zebrafish Smad6a (ENSDARP00000112408) and Smad6b (ENSDARP00000091342) respectively. Smad6a and Smad6b show 58.33% protein sequence similarity, which is considered large enough to assume similar function^19^. The sequence similarity and the same spatial expression patterns of *Smad3a* and *Smad3b* as well as *Smad6a* and *smad6b* suggest^20–22^.

In this study, we investigated modifying effects of *smad3a/smad3b* on *smad6a/smad6b* and vice versa which enabled us to model TAAD in zebrafish. Additionally, we identified early molecular mechanisms in aortic failure to identify druggable pathways.

## METHODS

### Ethics statement and housing

This study was approved by the Animal Ethics Committee of the Ghent University Faculty of Medicine and Health Sciences (Permit number: ECD 14/70) and housed adhering to the general guidelines, in agreement with EU Directive 2010/63/EU for laboratory animals^23,24^.

### Zebrafish lines: design and generation

*Smad3a*^sa2363/+^ was acquired from the Zebrafish International Research Center (ZIRC). All other models were generated using CRISPR/Cas9 gene editing technology utilizing CRISPRdirect software^25–27^. Specific gBlocks for all target sites are listed in **Supplementary Table 1**. Genotyping primers are given in **Supplementary Table 2**. All variants are described using Genome Reference Consortium Zebrafish Build 11 (GRCz11) and described in **Supplementary Table 3**.

### Vasculogenesis in embryos

Zebrafish were outcrossed to the transgenic *Tg(fli1:EGFP)* line in order to visualize endothelial cells as previously described^28^. Head length was used as normalization factor. An overview of the acquired measurements and specific breeding schemes are given in **Supplementary Fig. 1a-c**.

### 3D reconstruction of the ventral aorta in adult zebrafish

Serial 5 µm paraffin cross sections of the entire ventral aorta of zebrafish between 6 and 13 months post fertilization (mpf) were stained with Weigert’s iron hematoxylin, rinsed in water and stained with Resorcin-Fuchsin for one hour followed by two wash steps in 95% ethanol and one wash with reverse osmosis (RO) water. Slides were dehydrated and mounted with Entellan mounting medium. Pictures of serial sections were aligned via the TrakEM2 1.0^29–31^ and 3D reconstructed with Mimics 24.0 software.

### Ultrasound analysis

Ultrasound analysis was performed on zebrafish of 6-7 mpf as previously described^32^. All measurements were performed in Vevo Lab 5.5.0.

### Acute net handling stress induction

Induction of handling stress was adapted from previously described procedures^33^. Six mpf zebrafish were netted and air suspended for 90 seconds, returned to the housing tank for 90 seconds, netted and air suspended for 90 seconds and returned to the housing tank.

### Alizarin Red staining for mineralized bone

Alizarin red staining for mineralized bone was performed as previously described^34^, at 13 mpf. The qKO were stained after sudden death between 6 and 13 mpf.

Ectopic bone on the sinistral skeletal elements of the vertebral bodies was scored on abdominal and caudal vertebrae of adult zebrafish in lateral position^35^.

Skull morphometrics were performed on images of the zebrafish heads in lateral position. Acquired lengths were measured in ImageJ and are given in **Supplementary Fig. 1b**. Diameter of the eye was used for normalization since eye diameter correlates with standard length^36^. Cranial surface area was measured with standard length as normalization factor.

### Transcriptome analysis

RNA sequencing was performed on five pools, each containing five qKO or WT control cousins. Following RNA extraction (RNeasy® Mini kit), libraries were prepared (TruSeq Stranded Total RNA kit from Illumina) and sequenced on a NovaSeq6000 instrument (Illumina).

Following quality control ^37,38^, raw reads were mapped (STAR^39^) and through R packages, additional normalization and filtering was performed^40–42^. The top list of DEG was obtain via differential expression analysis (edgeR), with the necessary adjustments to account for multiple testing^43,44^. An adjusted p-value of 0.05 and a fold change of 2 were used as thresholds to determine genes of relevance. Finally gene enrichment analysis was carried out^45^. Further details can be found in the **Supplementary Methods**.

### Synchrotron phase contrast micro Computed tomography imaging

Propagation-based phase-contrast synchrotron X-ray imaging was performed at the TOMCAT (X02DA) beamline of the Swiss Light Source (Paul Scherrer Institute in Villigen, Switzerland) as previously described^46^. Tomographic reconstruction was performed using the Gridrec algorithm after applying the Paganin phase retrieval method^47,48^.

### 3D Biomechanical modeling to assess principal stress at systolic peak in the aortic wall

In a previous study^46^, zebrafish-specific 3D biomechanical fluid-structure interaction models of five different 13-month-old zebrafish were developed. All parameter settings as described in^46^ were maintained except for flow through right aortic arch 2, 3 and 4, which was set to nearly zero in order to mimic the vascular organization found in qKO mutants.

### Transmission electron microscopy

In short, WT and qKO zebrafish of 16 mpf were euthanized, fixed in Karnovsky fixative, followed by decalcification for three weeks. Post-fixation in 1 % osmium tetroxide, 0.1M cacodylate buffer with 8% saccharose and 0.004% CaCl_2_ was followed by *en bloc* staining in 1 % aqueous uranyl acetate. After dehydration, the samples were embedded in Spurr’s resin. 80 nm sections were stained with uranyl acetate and lead citrate and examined by transmission electron microscopy (TEM) (JEM 1010, JEOL) equipped with a CCD side-mounted Veleta camera (EMSIS).

### Swimming behavior studies

Swimming behavior of 9 mpf WT, *smad3a^+/−^;smad3b^−/−^;smad6a^+/−^;smad6b^−/−^* and qKO zebrafish was analyzed in a custom-made dark observation chamber (Noldus) equipped with the Basler GenICam. Data was analyzed using the EthoVision XT 17 software (Noldus). Zebrafish were placed individually in a tank and could acclimatize for 10 minutes before a 10-minute test period, was performed in the dark. Movement is depicted as total distance travelled during the test period, normalized for standard length. Movement frequency was also recorded.

### 1-phenyl-2-thiourea treatment

At 1 day post fertilization (dpf), 0.003% 1-phenyl-2-thiourea treatment (PTU) in E3 medium was administered to WT zebrafish. The medium was changed daily until 10 dpf. Starting from 5 dpf, zebrafish were given dry food for 2 hours daily, after which the medium was refreshed again.

### Statistical analyses

All statistical analyses were performed with GraphPad Prism version 10.1.1 for Windows (GraphPad Software, Boston, Massachusetts USA, www.graphpad.com). Kruskal-Wallis test with Dunn’s multiple comparisons test against WT controls was used to determine if there was an increase in ectopic bone on the skeletal elements of the vertebral bodies. For PTU treatment, a two-tailed t-test was used, when standard deviation differed significantly between the groups, Welch’s correction was applied. For all other comparisons, One-way ANOVA with Dunnett’s multiple comparison test against WT controls was used. Brown-Forsythe and Welch ANOVA with Dunnett’s T3 multiple comparison test against WT controls was used when the genotypes showed a significantly different standard deviation, determined with the Brown-Forsythe test.

## RESULTS

### Disruption of one of the smad3 or smad6 ohnologs does not show a cardiovascular or skeletal phenotype in zebrafish

Using CRISPR/Cas9 gene editing technology, we generated frameshift indels in the *smad3b*, *smad6a*, and *smad6b* genes, resulting in the generation of a premature termination codon in each mutant: *smad3b*^c.455_459delinsATG/c.455_459delinsATG^, *smad6a*^c.905_906delTG/c.905_906delTG^ and *smad6b*^c.283_287delACGGT/c.283_287delACGGT^ (**Supplementary Table 3**). To study *smad3a* function we made use of the available *Smad3a*^sa2363/+^ line, in which a premature termination codon is introduced (c.682G>T). For the reader’s benefit, we will use the term single knockout (SKO) for *smad3a, smad3b, smad6a* and *smad6b* single homozygous knockout zebrafish. At 5 dpf, measurement of the ventral aorta in the transgenic *Tg(fli1:EGFP)* reporter background is similar for heterozygous and SKO zebrafish for each gene as compared to sibling controls (**Supplementary Fig. 2**).

However, incross of *smad3a* SKO results in100% embryonic lethality at 2 dpf with severe body axis deformities and pericardial edema formation, while incross of *smad3b* SKO and *smad3a^+/−^;smad3b^−/−^* shows offspring with normal survival, suggesting that maternal *smad3a* expression is necessary for early embryonic survival, which is supported by previously published results^20^. Incross of *smad6a* or *smad6b* SKO showed normal survival.

In 6-7 mpf adult zebrafish, ultrasound measurements show that all measured parameters of cardiac function are similar in all SKO compared heterozygous mutants and sibling controls, with an exception of *smad6a^−/−^* which shows a small increase in normalized projected surface area of the ventricle in diastole (**Supplementary Fig. 3-4**).

In addition, skeletal evaluation and alizarin red staining for mineralized bone at 13 mpf do not show any vertebral column deformities or an increase in ectopic bone formation (**Fig. 4, Supplementary Fig. 5-6**). Skull morphometrics are normal, except for the *smad3a* SKO, in which a decrease of the snout to frontal bone distance is observed (**Supplementary Fig. 7-8**).

### Smad3a/b DKO and smad6a/b DKO show altered vasculogenesis

To bypass embryonic lethality due to loss of maternal *smad3a*, we performed a cross between a *smad3a^+/−^;smad3b^−/−^* female and *smad3a/b* DKO male zebrafish to obtain 50% *smad3a^+/−^;smad3b^−/−^* and 50% *smad3a/b* DKO zebrafish, which survive normally. At 10 dpf, the *smad3a/b* DKO show a significant increase in normalized surface area of the aortic segment between aortic arch 3 and 4 (p = 0.0449), a reduced normalized length of the segment (p = 0.0366) and an increased normalized mean aortic diameter (p = 0.0335). The diameter adjacent to the bulbus arteriosus remains unchanged (**Supplementary Fig. 9**).

The *smad6a/b* DKO show normal early survival rates, as indicated by normal Mendelian distribution of all genotypes in 10 dpf-old offspring obtained from *smad6a^+/−^;smad6b^+/−^* incrosses. Nevertheless, *smad6a/b* DKO zebrafish are unable to survive to adulthood.

*Smad6a/b* DKO zebrafish show a significant reduction of the ventral aortic diameter at 10 dpf (p<0.0001). The diameter adjacent to the bulbus arteriosus as well as the normalized surface area of the aorta segment is decreased (p<0.0001, p<0.0001). To a lesser extent, the diameter of the aorta segment of *smad6a^+/−^;smad6b^−/−^* is also reduced (p = 0.0016), while the length of the aorta segment remains unchanged in all *smad6* knockouts (**Supplementary Fig. 9**).

Ultrasound analysis at 6-7 mpf of all viable *smad3* (*smad3a^+/−^;smad3b^−/−^*, *smad3a^−/−^;smad3b^+/−^* and *smad3a/b* DKO) or *smad6* (*smad6a^+/−^;smad6b^−/−^* and *smad6a^−/−^;smad6b^+/−^*) KO combinations do not show any significant differences in cardiac parameters (**Supplementary Fig. 3**-4).

### Loss of smad3 ohnologs in zebrafish induces ectopic bone formation and vertebral column deformations

Alizarin red staining for mineralized bone on 13 mpf *smad3a/b* DKO zebrafish shows ectopic bone formation on the ribs, neural arches, neural spines, haemal arches, and haemal spines (p = 0.0012) (**Fig. 4**) as well as hyperlordosis, notochord mineralization and notochord sheet mineralization (**Supplementary Fig. 5**). Overall, the thickness of the haemal arches is increased (**Fig. 4**) and the normalized surface area of the cranial roof is reduced compared to WT controls (p = 0.0101) (**Supplementary Fig. 8**). The presence of a single WT copy of either *smad3a* or *smad3b* prevents any of these skeletal malformations to develop (**Fig. 4, Supplementary Fig. 8**). Skull morphometrics show a mild decrease of the distance between the tip of the snout and the most posterior part of the supraoccipital bone in s*mad3a^−/−^* zebrafish (p = 0.0486), but not in the *smad3a/b* DKO (**Supplementary Fig. 7**).

Likewise, a significant increase in ectopic bone can be seen in 13 mpf *smad6a^−/−^ ;smad6b^+/−^* (p = 0.0022) as well as in *smad6a^+/−^;smad6^−/−^* (p = 0.0033) zebrafish, although to a lesser extent than observed in the *smad3a/b* DKO (**Fig. 4**). Since the *smad6a/b* DKO does not reach adulthood, these features cannot be analyzed in this genotype. No premature fusion of the cranial roof bones is observed, and the normalized surface areas of the cranial roof are similar to controls (**Supplementary Fig. 8**). Skull morphometrics do show a decrease in length between the tip of the snout and tip of the frontal bone in *smad6a^+/−^;smad6b^−/−^* mutants (p = 0.0056) (**Supplementary Fig. 7**).

### Loss of smad3 and smad6 ohnologs in zebrafish induces ventral aortic dissection and rupture

To investigate potential modifying effects of *smad3* on *smad6* and vice versa, we generated a *smad3a^−/−^;smad3b^−/−^;smad6a^−/−^;smad6b^−/−^* qKO zebrafish line (**Supplementary Table 3**).

Unlike the *smad6a/b* DKO, qKO zebrafish reach adulthood and generate viable offspring. Microscopic analysis of vasculogenesis at 10 dpf shows a severe decrease of the normalized surface area of the aortic projection (p<0.0001), the normalized mean diameter of the aortic segment between aortic arch 3 and 4 (p = 0.0241), and the normalized diameter adjacent to the bulbus arteriosus (p = 0.0005) compared to WT controls, similar to *smad3a^+/−^;smad3b^−/−^;smad6a^+/−^;smad6b^−/−^* zebrafish (p<0.0001, p<0.0001, p = 0.0032, respectively). The normalized length of the aortic segment is only decreased in the qKO (p = 0.006) (**Fig. 1, Supplementary Fig. 9**). In addition, the qKO shows tortuosity of the ventral aorta, aortic arches and hypobranchial artery (**Fig. 1, Supplementary Fig. 10**).

**Figure 1:**
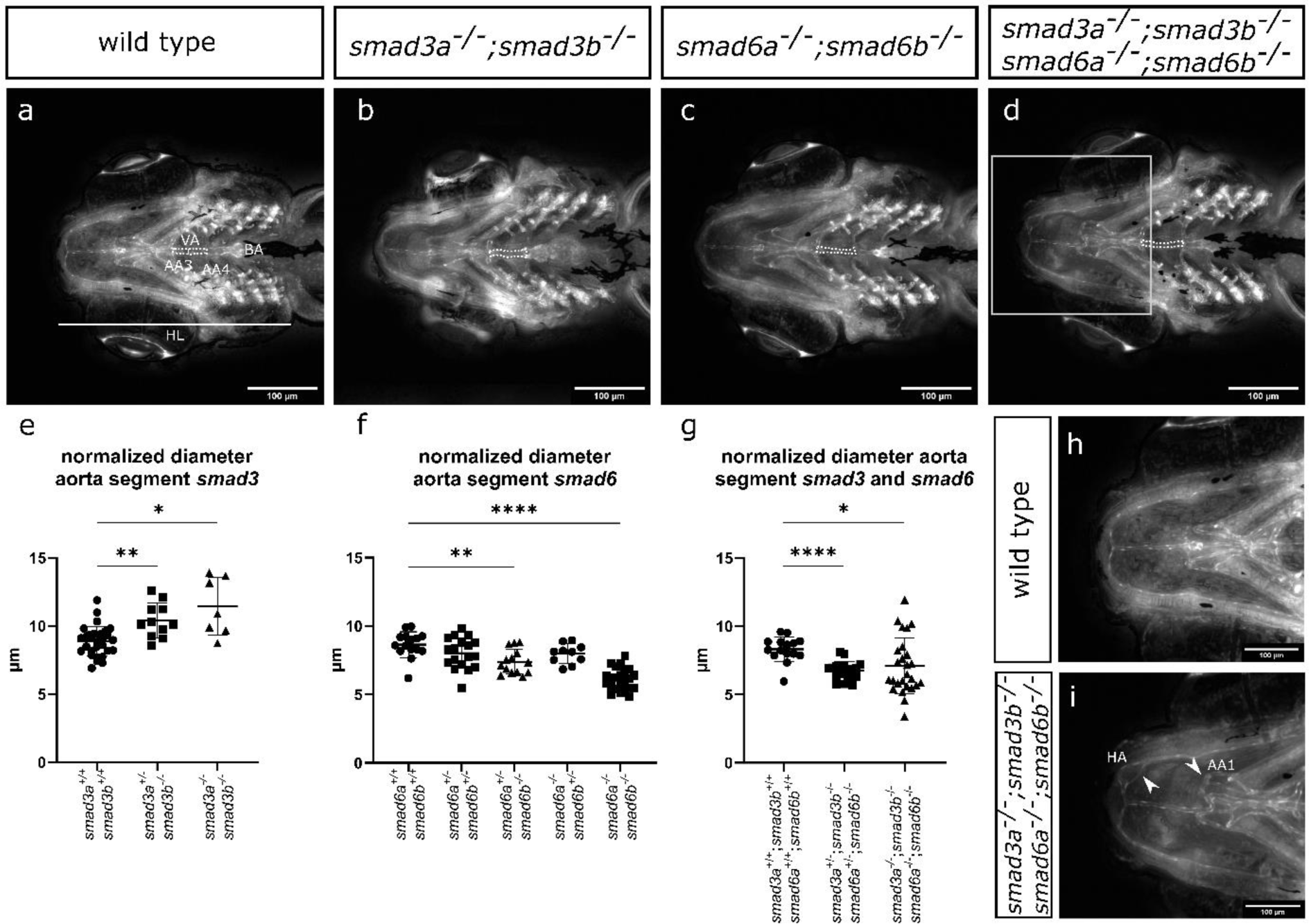
Altered vasculogenesis at 10 dpf in double and qKO mutants. **(a-d)** Ventral view of the cardiovascular structures at 10 dpf of *smad* knockout lines crossed with a *Tg(fli1:EGFP)* reporter line. Legend: “VA” ventral aorta, “BA” bulbus arteriosus, “AA3” aortic arch 3, “AA4” aortic arch 4, “HL” head length. The projected surface area of the ventral aorta between aortic arch 3 and aortic arch 4 has been outlined with a white dashed line. **(e)** *smad3a^+/−^;smad3b^−/−^ and smad3a^−/−^;smad3b^−/−^* zebrafish of 10 dpf show an increase in normalized aorta diameter (n=29-11-7). **(f)** *smad6a^+/−^;smad6b^−/−^* as well as *smad6a^−/−^;smad6b^−/−^* show a decrease in diameter of the ventral aorta (n=15-18-14-10-23). **(g)** *smad3a^+/−^;smad3b^−/−^;smad6a^+/−^;smad6b^−/−^* and *smad3a^−/−^;smad3b^−/−^;smad6a^−/−^;smad6b^−/−^* both show a decrease in diameter of the ventral aorta (n=15-18-26). **(h,i)** Overview of the hypobranchial artery in a WT control and qKO zebrafish line. Tortuosity and irregular branching of the hypobranchial artery (HA) and aortic arch 1 (AA1) can be observed **(i)** magnification of boxed area in d. **(a-d,h,i)** stack focused Z-stack pictures obtained with ZEISS Axio Observer.Z1 microscope **(f)** One-way ANOVA with Dunnett’s multiple comparison test against WT controls. **(e,g)** Brown-Forsythe and Welch ANOVA with Dunnett’s T3 multiple comparison test against WT controls. Asterisks indicate significant differences. *p<0.05, **p<0.01, ***p<0.001, ****p<0.0001. Data represented as average ± standard deviation.

In adult qKO zebrafish, we observed sudden deaths, mostly after breeding, netting, or changing the tank positions. To investigate stress as a trigger, we performed a netting test that is harmless in WT controls, but results in a mortality rate of 60% in qKO. Sudden death associated with a red discoloration anterior in the abdominal region. Sectioning of the entire ventral aorta and staining with Resorcin-Fuchsin to visualize the elastic fibers shows the presence of aortic intramural hematomas and false lumens in multiple qKO. 3D-reconstructions of the aorta based on histological images and from synchrotron micro-CT images of intact zebrafish, confirm the presence of regions of ventral aortic damage (**Fig. 2** and **Supplementary movie 1-8**).

**Figure 2:**
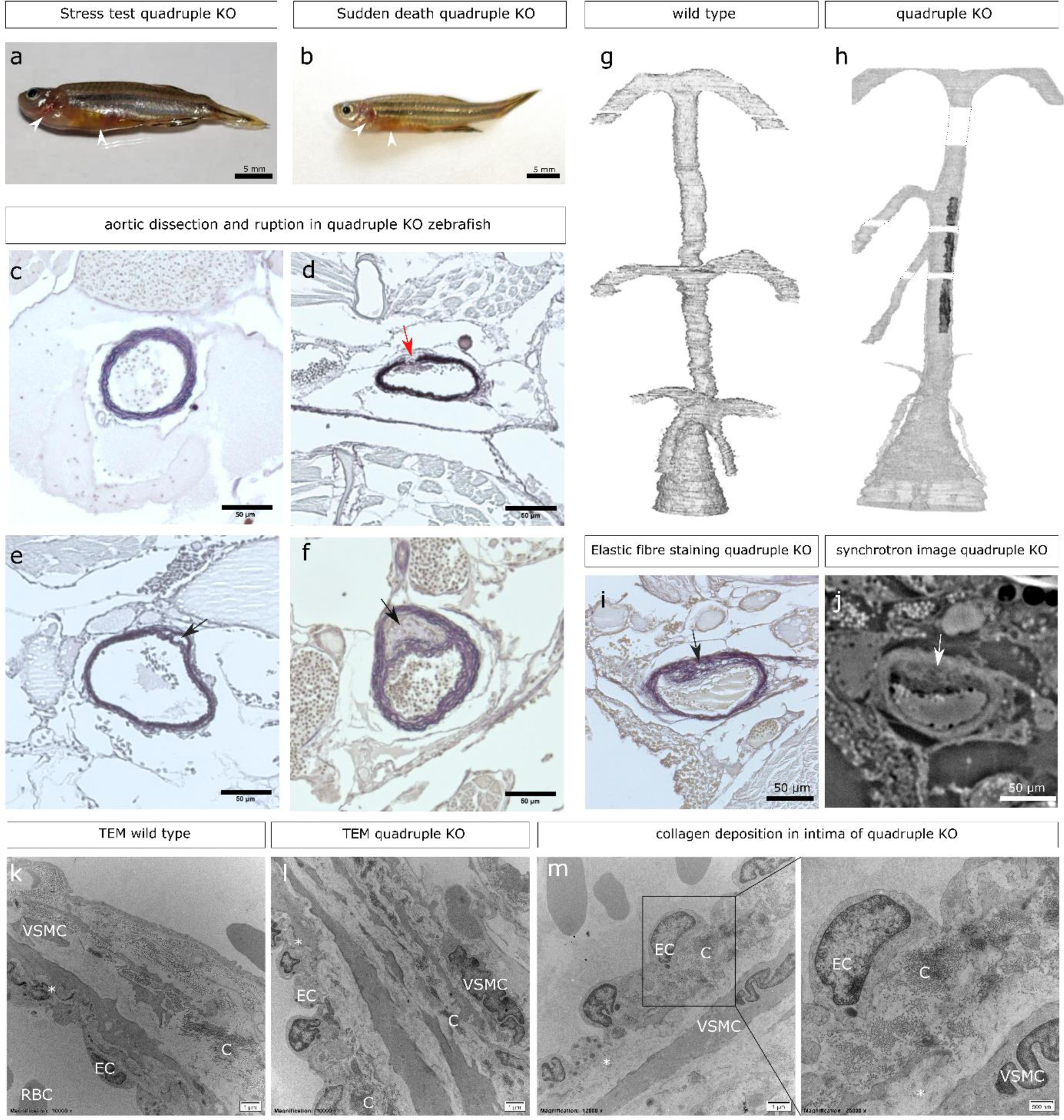
qKO zebrafish exhibit thoracic aortic dissection and rupture. **(a, b)** qKO zebrafish that died suddenly show similar external features compared to qKO zebrafish that die due to stress induction. Red discoloration around the heart and gill area (anterior abdominal area) is indicated with a white arrowhead. **(c)** Resorcin-Fuchsin staining of WT control. **(d)** Resorcin-Fuchsin staining of the elastic fiber on a 5 µm cross-section of the ventral aorta shows rupture of the aortic wall (red arrow). **(e,f)** Resorcin-Fuchsin staining of the elastic fiber on 5 µm thick cross-sections of the ventral aorta. False lumen is indicated with a black arrow. **(g,h)** 3D reconstruction using 3D-modelling software Mimics 24.0 of the ventral aorta shows a decrease in diameter of the aortic arches. Symmetrical branching of the aortic arches is lost in the qKO model. Region with aortic damage has been colored black **(i,j).** The area of damage identified on a histological resorcin-fuchsin staining of the ventral aorta can similarly be observed on a synchrotron micro-CT scan of the same region. Damage is indicated with arrows. **(k,l)** TEM of aortic wall in WT and qKO, respectively. Severe decrease of elastic fiber in the inner elastic lamina is apparent. **(m)** TEM of qKO shows collagen deposition in intima. **(k,l,m)** Legend: “VSMC” vascular smooth muscle cell, “EC” endothelial cell, “RBC” red blood cell, “C” collagen, * indicates inner elastic lamina.

Regions of ventral aortic damage start close to a branching point (6/6 qKO, 0/4 WT) and dissection or rupture is present at the most affected sites, (5/6 qKO, 0/4 WT). The diameter of the aortic branches is asymmetrically distributed in all qKO zebrafish (6/6 qKO, 0/4 WT) with one side consistently showing narrower branches than the other. In more severe cases, one or more aortic arches are missing (2/5 qKO, 0/4 WT) or attached at the wrong side of the ventral aorta (1/6 qKO, 0/4 WT) (**Supplementary Movie 9-13**). No morphological differences can be detected in WT controls (**Supplementary Movie 14-16**) or s*mad3a/b* DKO zebrafish (**Supplementary Movie 17-19**).

TEM shows a severely decreased elastin deposition in the internal elastic lamina of 15 mpf qKO zebrafish. Additionally, abnormal collagen deposition is detected near the intima of the ventral aorta (**Fig. 2**).

Cardiac ultrasound analysis in 6-7 mpf qKO zebrafish shows regurgitation on ventricular in-(p = 0.0009) and outflow (p<0.0001), suggestive of valve dysfunction (**Supplementary Fig. 11**). Elastin staining of paraffin sections of the valves shows hypertrophy of the valve interstitial cells, resulting in a more rounded appearance of the qKO bulboventricular valve in contrast to the normal heart-shaped structure in WT control zebrafish. Similarly, the atrioventricular valve shows hypertrophy and altered morphology of the valve interstitial cells (**Fig. 3, Supplementary Fig. 12**).

**Figure 3:**
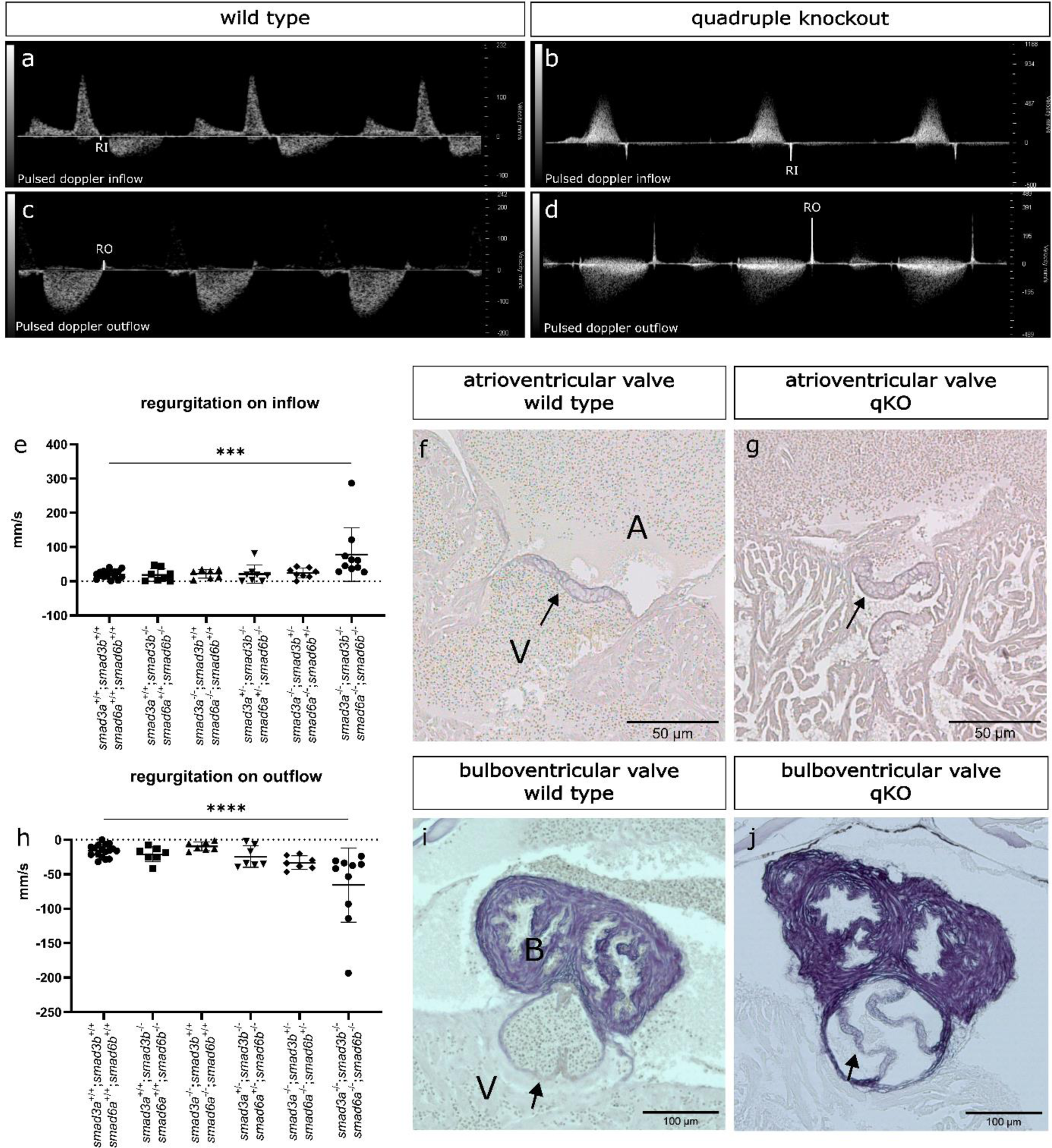
Increased regurgitation and abnormal valve morphology in qKO zebrafish. **(a, c)** Pulsed wave doppler of inflow **(a)** and outflow **(c)** of blood in and out of the ventricle of WT control zebrafish. **(b, d)** Pulsed wave doppler of inflow **(b)** and outflow **(d)** of blood in and out of the ventricle of qKO zebrafish. **(e)** Regurgitation on inflow is significantly increased in the qKO zebrafish model (n=17-8-7-8-8-10). **(f, g)** Resorcin Fuchsin staining shows hypertrophy of the valve interstitial cells and abnormal leaflet morphology in the atrioventricular valve of qKO zebrafish. **(h)** Increased regurgitation on outflow in qKO (n=17-7-7-7-7-10). **(i,j)** Resorcin Fuchsin staining of the bulboventricular valve in WT and qKO zebrafish shows hypertrophy of the valve and altered orientation in the qKO. **(a-j)** Legend: “RI” regurgitation inflow, “RO” regurgitation outflow, “A” atrium, “V” ventricle, “B” bulbus arteriosus. Black arrows indicate valves. **(e,h)** One-way ANOVA with Dunnett’s multiple comparison test against WT controls. Asterisks indicate significant differences. ***p<0.001, ****p<0.0001. Data represented as average ± standard deviation.

### Aortic damage in the qKO zebrafish correlates with increased stress at branching regions

In qKO zebrafish, asymmetric branching and reduction of the diameter of the aortic arches is apparent. To determine if the morphological changes of the aorta in adult qKO zebrafish increase the principal stress within the vessel wall and to identify which locations endure the largest differences, aortic morphology was compared using *in silico* simulation models representing 13 mpf WT and qKO zebrafish aortae. To model the severe reduction of the diameter of the qKO aortic arches, the blood flow through the right aortic arches (aortic arch 2, 3 and 4) was reduced to nearly zero in the model. The principal stress within the aortic wall was increased in all 5 qKO simulations, with the most affected region close to branching points. This corresponds to the *ex vivo* observations of the location of regions of damage in the ventral aorta of qKO zebrafish (**Supplementary Fig. 13**).

### qKO zebrafish show skeletal aberrations, but largely normal swimming behavior

The qKO show an increased presence of ectopic bone formation (p = 0.0014). Scoliosis, hyperlordosis and hyperkyphosis, bending of the arches and ribs, as well as intervertebral ligament mineralization and notochord mineralization are apparent in the qKO zebrafish only (**Fig. 4, Supplementary Fig. 14**).

**Figure 4:**
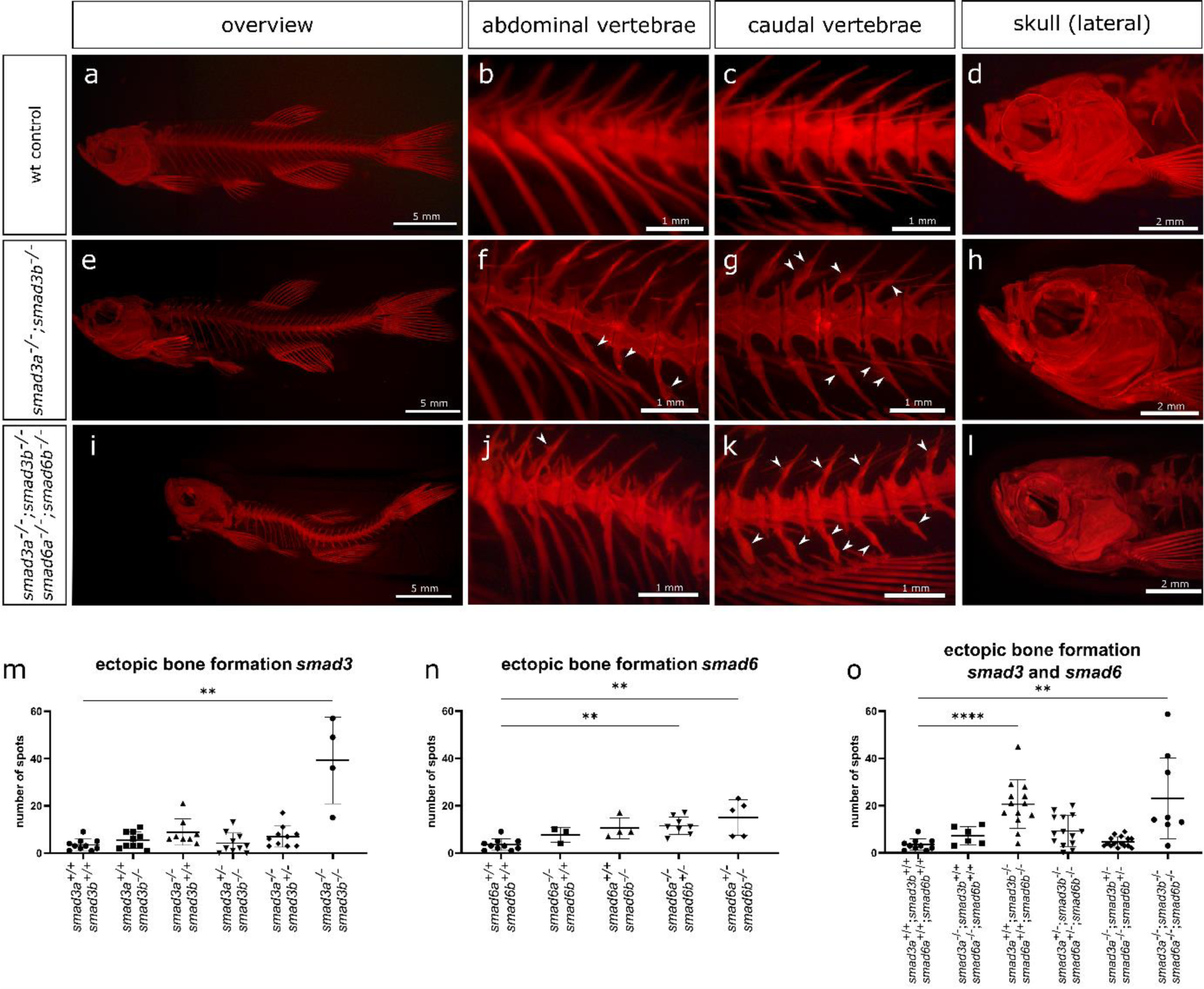
qKO zebrafish show scoliosis, lordosis and ectopic bone formation in the vertebral column. **(a-l)** Overview pictures of alizarin red staining for mineralized bone of abdominal and caudal vertebrae, and the skull (lateral view) of WT (**a-d**), *smad3a^−/−^;smad3b^−/−^* DKO zebrafish (**e-h**), and *smad3a^−/−^;smad3b^−/−^;smad6a^−/−^;smad6b^−/−^* qKO zebrafish (**i-l**). Ectopic bone has been indicated with white arrowheads. Scoliosis and lordosis can be observed in (**i**). **(k)** Notochord sheet mineralization can also be observed between the vertebrae. Small craniofacial aberrations are detected. **(l)** The nasal bone structures appear to be missing, resulting in a clockwise rotation of the snout. **(m-o)** Quantification of spots of ectopic bone in *smad3* (n=10-10-8-10-10-4), *smad6* (n=10-4-3-8-5), *smad3/smad6* (n=10-6-13-14-15-8) genotypes. **(a-l)** Whole-mount alizarin red staining for mineralized bone pictures taken with Leica M165 FC Fluorescent Stereo Microscope. **(m-o)** Kruskal-Wallis test with Dunn’s multiple comparisons test against WT controls. *p<0.05, **p<0.01, ***p<0.001, ****p<0.0001. Data represented as average ± standard deviation.

In qKO zebrafish, the nasal region of the skull is underdeveloped with a significantly reduced distance between the tip of the snout and tip of the frontal bone (p<0.0001). (**Fig. 4, Supplementary Fig. 7**). Interestingly, qKO show normal values for the normalized surface area of the cranial roof in contrast to the *smad3a/b* DKO, which was significantly decreased (p = 0.0101) (**Supplementary Fig. 8**). Since qKO embryos show a tortuous hypobranchial artery, which provides blood flow to the teeth, tooth morphology was evaluated, but appears normal (**Supplementary Fig. 15**).

At 9 mpf, qKO zebrafish show a decrease in movement frequency (p = 0.0197), but not in normalized total distance traveled compared to WT zebrafish (**Supplementary Fig. 16**).

### *Bulk RNA sequencing of 5 dpf qKO* shows a reduction in the negative regulation of proteolysis and upregulation of melanogenesis

Bulk RNA sequencing in 5 dpf qKO and 2^nd^ degree related WT control zebrafish identified 1131 differentially expressed genes (DEG) (adjusted p-value <0.05) (**Supplementary Table 4**). The top 10 enriched GO-terms in the upregulated gene set mostly involve pigmentation (pigment metabolic process, pigment biosynthetic process, melanosome organization, pigment granule organization, melanocyte differentiation, pigmentation, developmental pigmentation). One of the central DEG driving this enrichment is the *mitfa* transcription factor, which is known to signal downstream of BMP and governs melanocyte differentiation and development via tyrosinase (log FC *tyr* 2.01), tyrosinase related protein 1 (log FC *tyrp1a* 1.82 and log FC *tyrp1b* 2.08), the premelanosome protein (log FC *pmela* 1.44 and log FC *pmelb* 3.66), dopachrome tautomerase (log FC *dct* 2.6), and the melanosomal transporter (log FC *slc24a5* 2.35)^49,50^. The top 10 enriched GO-terms in the downregulated gene set are linked to negative regulation of proteolysis (negative regulation peptidase activity, negative regulation proteolysis, regulation of peptidase activity, negative regulation of endopeptidase activity, regulation of proteolysis) and immune response (regulation of immune response, positive regulation of immune system process) (**Supplementary Table 5-6**). A heatmap with stringent thresholds (p.adjust ≥ 0.0001 and |log2FC| ≥ 1) and a volcano plot are shown in **Fig. 5**. Annotated dysregulated genes with a known role in aortic wall homeostasis, include *loxl3a*, *emilin3a* and *fbn2b* (elastic fiber assembly), *vcana* and *lamc2* (vascular cell adhesion glycoproteins), *mhc1uma* (a zebrafish specific MHC class I gene), and *uts2a* (a vasoconstrictor linked to hypertension and heart failure in humans). Dysregulation of *omd* (a transcription factor, activated via BMP signaling, that links to cell adhesion and bone mineralization) might relate to the observed bone phenotypes.

**Figure 5:**
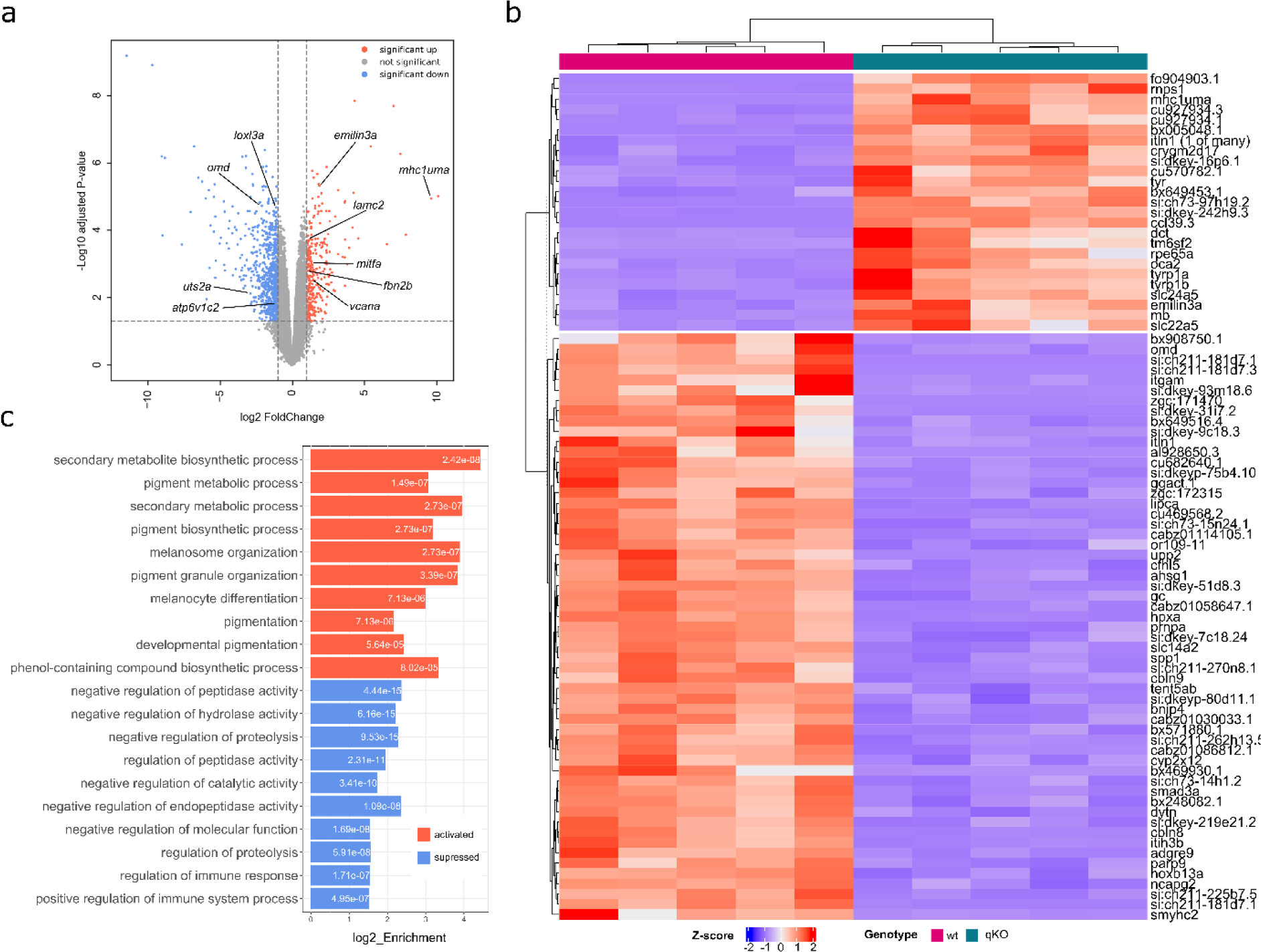
RNA sequencing data shows downregulation of negative regulators of proteolysis and upregulation of melanogenesis. **(a)** Volcano Plot showing upregulated genes (red) and downregulated genes (blue) with genes of importance annotated. Thresholds: p.adjust ≥ 0.05 and |log2FC| ≥ 1; **(b)** Heatmap with top DEG results between WT and qKO. Thresholds: p.adjust ≥ 0.0001 and |log2FC| ≥ 1 ; **(c)** Top 10 hits of GO enrichment analysis for both sets of upregulated (red) and downregulated (blue) genes. Numbers inside the bars represent the corresponding adjusted p-value value.

### Tyrosinase inhibition increases diameter of the ventral aorta in 10 dpf zebrafish

To investigate the involvement of melanogenesis-related pathways in aortic development, we administered 0.003% PTU, a known inhibitor of tyrosinase in the medium of WT zebrafish embryos starting at 1 dpf. At 10 dpf, treated zebrafish show a significant increase of normalized ventral aortic surface area (p = 0.0066) and mean normalized aorta diameter between aortic arch 3 and 4 (p = 0.0065) compared to vehicle control. Length of the aortic segment and diameter adjacent to the bulbus arteriosus remained unchanged compared to non-treated sibling controls (**Fig. 6**).

**Figure 6:**
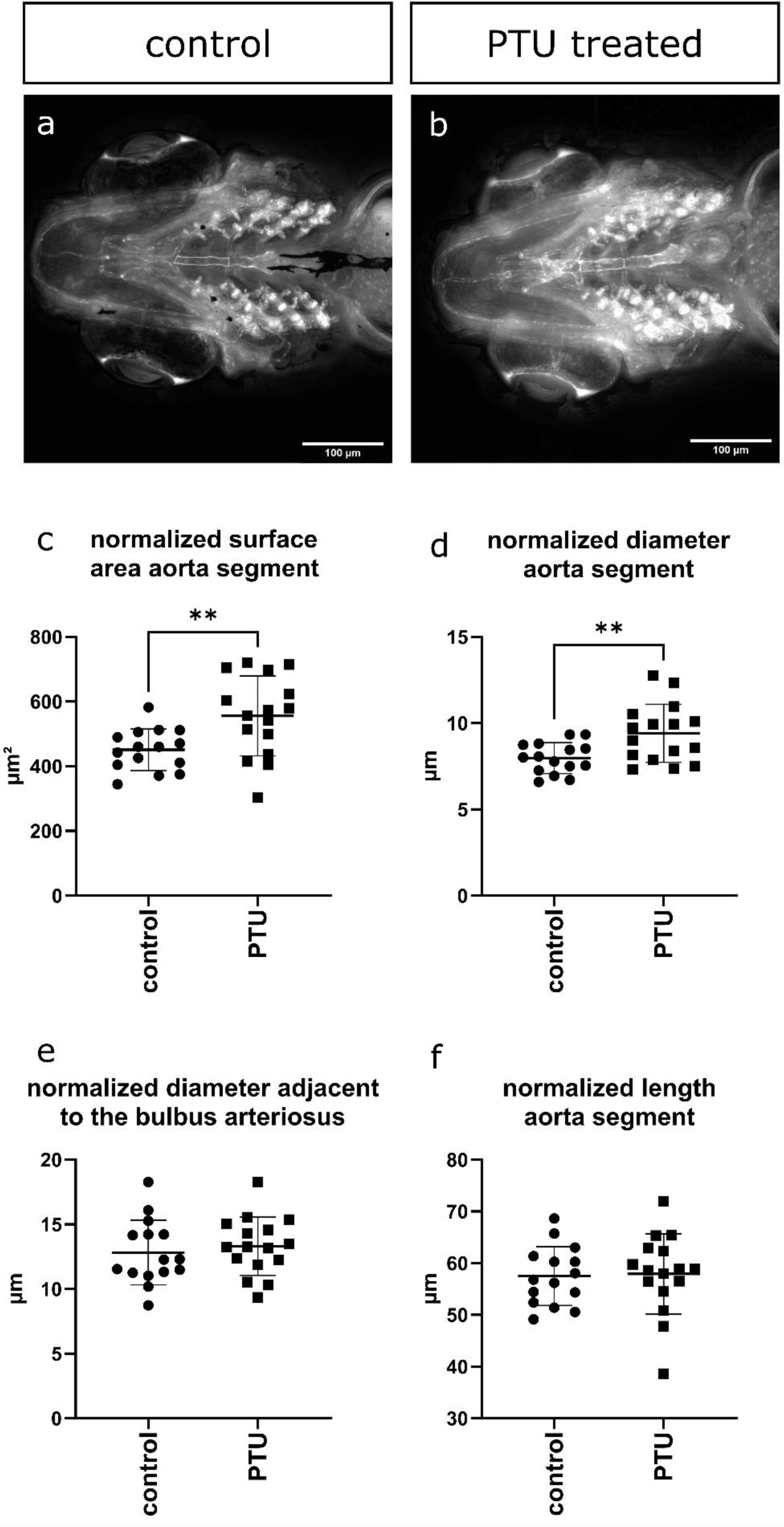
Tyrosinase inhibition increases the aorta diameter of 10 dpf zebrafish. **(a)** Ventral view of the vasculature of 10 dpf WT control zebrafish. **(b)** Ventral view of the vasculature of 10 dpf WT zebrafish treated with 0.003% PTU treatment starting at 24 hpf. **(c, d)** PTU-treated zebrafish show a significant increase in normalized surface area of the aortic segment and aortic diameter (n=15-16). **(e,f)** Normalized diameter adjacent to the bulbus arteriosus and length of the aorta segment is similar between treated and non-treated zebrafish (n=15-16). **(a,b)** Dashed white line indicates aorta segment between aortic arch 3 and 4. Stack focused Z-stack pictures obtained with ZEISS Axio Observer.Z1 microscope**. (c,d)** Unpaired two-tailed t-test with Welch’s correction. **(e,f)** Unpaired two-tailed t-test. Asterisks indicate significant differences. **p<0.01. Data represented as average ± standard deviation.

## DISCUSSION

TGFβ and BMP signaling show significant crosstalk during vascular and skeletal development and homeostasis. Our results support the existence of modifier effects particularly on aortic phenotypes associated with *SMAD6*-deficiency, which range from bicuspid aortic valve with or without ascending aortic dilatation to Shone complex with a hypoplastic ascending aorta and aortic arch.

In the genome of mammals and teleost fish, *SMAD3* and *SMAD6* locate closely together, but in different topologically associated domains^16^. Since zebrafish express *smad3a*, *smad3b, smad6a* and *smad6b*^51–54^, this model provides flexibility to investigate the interactions between these genes upon gene dosage. Although maternal *smad3a* is required during early embryogenesis, which is supported by previous findings^48^, SKO show no obvious vascular or skeletal phenotype, except for mild craniofacial alterations in *smad3a^−/−^* zebrafish. However, *smad3a/b* or *smad6a/b* DKO do manifest cardiovascular and mild skeletal phenotypes, supporting compensation between the ohnologs of either smad3 or smad6, in line with their strongly conserved protein structure ^19^.

*Smad6a/b* and *smad3a/b* DKO show opposite vascular phenotypes. *Smad3a/b* DKO zebrafish show an increase in aortic diameter in early larval stages. This aligns with previous findings in which loss of *alk5^−/−^*, the type 1 TGF-β receptor which activates SMAD3, leads to an increased diameter of the cardiac outflow tract in zebrafish embryos, followed by a failed development of the ventral aorta causing death at 7 dpf^55^. It is plausible that in *smad3a/b* DKO limited amounts of Smad2/Smad4 complexes partially rescue the phenotype preventing early death and an adult cardiovascular phenotype, in contrast to Alk5 loss of function, in which both Smad2 and Smad3 will not be activated^56^.

S*mad6a/b* DKO show a reduced aortic diameter, and early death. The qKO also shows a reduction of the aortic diameter and arterial tortuosity, but survives to adulthood. The observation that the *smad3a^+/−^;smad3b^−/−^;smad6a^+/−^;smad6b^−/−^* shows a more severe reduction in aortic diameter than the qKO might be explained by the exclusion of the most affected qKO embryos, since clear edge detection of the ventral aorta in these embryos is hindered due to excessive tortuosity. Hence, it seems reasonable to generalize that smad3 loss of function increases, while lack of smad6 decreases the diameter of the aorta. The overall net result in the qKO shows a decreased diameter of the ventral aorta with asymmetrical branching and a high susceptibility for dissection and rupture in adult zebrafish. It has been previously reported that zebrafish *smad6b* is involved together with BMP in modulation of lateral sprouting of the dorsal aorta^54^.

Abnormal aortic branching results in altered hemodynamics and elevated systolic wall stress, as confirmed by our fluid-structure interaction models presented here. Areas exposed to perturbed local flow patterns prove prone to aortic wall damage and dissection, as demonstrated by histological and synchrotron imaging of the aorta. TEM analysis reveals a severe reduction of elastic fiber deposition in the inter-elastic laminae, which can be linked to a lessened resilience to hemodynamic stress. Additionally, abnormal collagen deposition near the intima of qKO zebrafish recapitulate increased collagen deposition in human TAAD^57^, indicating that similar pathophysiological processes are activated in our zebrafish model. The presence of an atrioventricular and bulboventricular valve defect in the qKO is in line with the role of SMAD6 in human bicuspid aortic valve disease^58^. It is conceivable that unopposed loss of both *smad6* ohnologs in zebrafish leads to a failure to maintain aortic homeostasis or to severe valve dysfunction, resulting in the premature death observed in the *smad6a/b* DKO late juvenile zebrafish.

The spectrum of bone phenotypes observed in the different genotypes confirms the modulatory interactions between the different *smad* genes. Indeed, *smad3a/b* DKO, but not *smad3a^+/−^;smad3b^−/−^* or *smad3a^−/−^;smad3b^+/−^* zebrafish, show increased ectopic bone formation, a decreased normalized cranial roof surface area, and vertebral column axis deviations. In contrast, the *smad6a^+/−^;smad6b^−/−^* and the *smad6a^−/−^;smad6b^+/−^* zebrafish do show an increase in ectopic bone formation, although it needs to be highlighted that adult *smad6a/b* DKO cannot be studied since they do not survive until adulthood. The qKO zebrafish show more ectopic bone formation, and more severe column axis deviations. These observations suggest a larger impact of BMP signaling on skeletal development, that is further impacted by loss of TGFβ signaling.

Enrichment analysis of bulk RNA sequencing data of qKO embryos at 5 dpf revealed upregulation of multiple pathways related to pigmentation, including the central transcription factor *mitfa* as well as downstream effectors in melanin production such as *tyrp1a*, *tyrp1b*, *tyr* and *dct*. MITF is a known downstream target of BMP signaling that is inhibited by SMAD6^59–61^. MITF also acts upstream of lysosome biogenesis and it was previously shown that lysosomal dysfunction in zebrafish due to loss of function of *atp6v1e1b* associates with dilatation of the ventral aorta and narrow aortic branches^28^. Finally, MITF is known to be important for cell metabolism and cell cycle regulation, promoting mitochondrial biogenesis and having complex effects on proliferation depending on the level of MITF activity^62^. It is conceivable that one or more of these effects of MITF are linked to the development of the phenotype observed in our qKO model.

The transcriptional changes of pigmentation-related pathways in the qKO model prompted us to test the consequences of pharmacological inhibition of tyrosinase, a key enzyme responsible for eumelanin production, at a dose commonly used in zebrafish research to block pigment formation. Inhibition of tyrosinase via PTU treatment resulted in an increase of the aorta diameter while a decreased aortic diameter in qKO at 10 dpf is associated with upregulated tyrosinase expression. This strongly supports a role for the pigment biosynthesis pathway in aortic homeostasis. Therefore, the use of PTU for imaging purposes should be approached with caution due to the potential for unintended effects on other organ systems, especially when evaluating cardiovascular phenotypes. Undesired effects of PTU have been reported previously including autophagy activation^63^, activation of cytochrome P4501A1 (*cyp1a1*)^64^, suppression of retinol-binding protein 4 (*rbp4*)^65^ and reduction of the diameter of the eye^66^.

Despite upregulation of melanin-producing pathways, no increase of skin pigmentation could be observed in the embryo or adult qKO zebrafish. Previous research showed that tyrosinase follows a specific spatiotemporal expression pattern during embryogenesis, which is not confined to skin cells^67^. At 7 dpf, small amounts of melanin can already be detected around the dorsal aorta, while at 1 month, melanin deposition is apparent^68^. It is therefore tempting to speculate that pigmentation-related pathways affect vascular development, particularly since it was recently shown that tyrosinase reduces expression of vascular endothelial growth factors^69^.

The relevance of the qKO model to study the disease mechanisms of dissection is further supported by the transcriptomic changes concordant with established models and known pathogenetic mechanisms of TAD. Altered elastic fiber homeostasis, which was confirmed in the qKO aortic wall using TEM, is reflected in upregulation of *Fbn2b*^70^ (logFC 1.04), a fibrillin known to be involved in endocardial morphogenesis, which might be upregulated as a response to the coarctation in the qKO, and downregulation of *mfap4* (logFC −3.26), an important component of the extracellular matrix (ECM) involved in elastic fiber assembly. In zebrafish, MFAP4 also regulates the balance between myeloid and lymphoid development^71^. Another immune cell-related gene expressed differently in qKO zebrafish is *itgam* (LogFC -5.33), coding for a component of the heterodimeric α_M_β_2_ integrin expressed on leukocytes which is also known as macrophage-1 antigen (Mac-1) or CD11b. This protein plays a crucial role in binding to ECM components and intracellular adhesion molecules, involved in adhesion and transmigration of leukocytes across blood vessels. The qKO transcriptomic profile also indicates that matrix proteolysis is increased, due to downregulation of the metalloproteinase inhibitor *timp4* (LogFC −3.28), in combination with upregulation of *mmp11b* (logFC 1.34) and *mmp13a* (logFC 1.35). These targets are all widely established in the pathogenesis of vascular and bone homeostasis ^72–74^. *Loxl3a*, an orthologue of the human *LOXL3*^75^ important for crosslinking of elastin and collagen, is significantly downregulated in qKO zebrafish. In mice, deletion of *Loxl3* results in abnormal skeletal development and is expressed in the precursors of the occipital and interparietal bones and nasal area, affected in the qKO model^76^.

In conclusion, Smad3-deficiency modifies Smad6-deficient phenotypes whereby *smad3a^−/−^;smad3b^−/−^;smad6a^−/−^;smad6b^−/−^* qKO zebrafish model aortic dissection and rupture, resulting from defective vasculogenesis and resultant local wall stress. In addition to known signatures of TAAD in human and mouse models, this model implicates the pigmentation pathway in the development of the aortic phenotypes. Follow-up studies on the precise contribution of these pathways are warranted and are likely to lead to novel targets for therapeutic intervention in TAAD.

## NON-STANDARD ABBREVIATIONS AND ACRONYMS

AA3: aortic arch 3
AA4: aortic arch 4
BMPs: bone morphogenetic proteins
Co-SMAD: common mediator SMAD
CPM: counts per million
DEG: differentially expressed genes
DKO: double knockout
dpf: days post fertilization
ECM: extracellular matrix
FDR: false discovery rate
GDFs: differentiation factors
GO: gene ontology
GRCz11: Genome Reference Consortium Zebrafish Build 11
GSEA: gene set enrichment analysis
ID: intellectual disability
I-SMAD: inhibitory SMAD
KEGG: Kyoto Encyclopedia of Genes and Genomes
LDS3: Loeys-Diets syndrome type 3
LOF: loss of function
mpf: months post fertilization
MR: magnetic resonance
PSI: Paul Scherrer Institut
PTU: 1-phenyl-2-thiourea treatment
qKO: smad3a^−/−^;smad3b^−/−^;smad6a^−/−^;smad6b^−/−^ quadruple knockout
RO: reverse osmosis
R-SMAD: receptor-regulated SMAD
SKO: single knockout
*smad3a/b* DKO: *smad3a^−/−^;smad3b^−/−^* double knockout
*smad6a/b* DKO: *smad6a^−/−^;smad6b^−/−^* double knockout
TAA: thoracic aortic aneurysm
TAAD: thoracic aortic aneurysm and dissection
TAD: thoracic aortic dissection
TEM: transmission electron microscopy
TGF-β: transforming growth factor β
ZIRC: Zebrafish International Research Center

## ACKNOWLEDGMENTS

The authors thank the family for their participation in this research. We also would like to thank the Zebrafish Facility Ghent Core at Ghent University, and particularly Karen Vermeulen for the diligent care for the zebrafish. We acknowledge the Paul Scherrer Institut (PSI), Villigen, Switzerland for provision of synchrotron radiation beamtime at the TOMCAT beamline X02DA of the Swiss Light Source, and would like to thank Karo De Rycke, Violette Deleeuw, Marina Horvat, Yousof Mohammad Asaad Abdel-Raouf (Ghent University) and Isaac Rodriguez Rovira (University of Barcelona) for their assistance in acquiring the synchrotron imaging data. Finally, we would like to thank the Nematology Research Unit, member of the UGent TEM Core Facility and the sequencing core of UZ Ghent.

## DATA AVAILABILITY

Raw RNAseq data is available at Gene Expression Omnibus with accession number GSE249792. To review GEO accession GSE249792. For reviewers, following link is available to access the data. https://eur03.safelinks.protection.outlook.com/?url=https%3A%2F%2Fwww.ncbi.nlm.nih.gov%2Fgeo%2Fquery%2Facc.cgi%3Facc%3DGSE249792&data=05%7C02%7Cmichiel.vanhooydonck%40ugent.be%7Cbeba6c62b40641ced96408dbfa282a45%7Cd7811cdeecef496c8f91a1786241b99c%7C1%7C0%7C638378822642521272%7CUnknown%7CTWFpbGZsb3d8eyJWIjoiMC4wLjAwMDAiLCJQIjoiV2luMzIiLCJBTiI6Ik1haWwiLCJXVCI6Mn0%3D%7C3000%7C%7C%7C&sdata=7g1JeMJOvj4lZVmQ6FVdkN%2FA2zrRUcuVlw6Hcear5vs%3D&reserved=0

Enter token obmhwiqkbbsprkt into the box.

All other data will be made available upon request.

## AUTHOR CONTRIBUTIONS

Conceptualization: MVH., PSI., BC. Methodology: MVH., MVE., MSS., LP., AKB., DS., PSI., BC. Software: MSS., MVI. Formal analysis: MVH., MSV., MVI., DS. Investigation: MVH., MVE., MVI., MSS., LP., AKB., HDS., LC., PT., DS. Resources: PSI., BC. Data Curation: MVH., MSS. Writing (original draft): MVH., MVE. Writing (review & editing): MVH., MVE., MSS., LP., AKB, MVI., HDS., LC., PT., AB., P.SE., ADC., AW., DS., PSI., BC. Visualization: MVH. MVE. Supervision: PSI., BC. Funding acquisition: PSI., BC.

## SUPPLEMENTARY MATERIAL

Supplementary Case Presentation

Supplementary Methods

Supplementary References

Supplementary Figures 1 – 17

Supplementary Tables 1 – 7

Supplementary Movies 1 – 19

## FIGURES WITH FIGURE LEGENDS

Links with access to high quality figures and the supplementary movies. Lower quality figures can be found below:

Supplementary Figure 1 https://figshare.com/s/1d96715da2007c11ae96

Supplementary Figure 2 https://figshare.com/s/7117068f173e59e3b7e7

Supplementary Figure 3 https://figshare.com/s/8a7ce09585ca754dedd1

Supplementary Figure 4 https://figshare.com/s/e754b6c96ea2c0588a15

Supplementary Figure 5 https://figshare.com/s/c902712aae5ffea8e862

Supplementary Figure 6 https://figshare.com/s/9bcafde922edd89090ad

Supplementary Figure 7 https://figshare.com/s/00039d2f1472d980fb15

Supplementary Figure 8 https://figshare.com/s/19aa5c0c33dfbf52f76b

Supplementary Figure 9 https://figshare.com/s/3b70b2174c355aafa4c8

Supplementary Figure 10 https://figshare.com/s/2ed90596b1477259fc3f

Supplementary Figure 11 https://figshare.com/s/5c8b0c3c23600bd54b15

Supplementary Figure 12 https://figshare.com/s/2c1733efb6b877eed39a

Supplementary Figure 13 https://figshare.com/s/2a70a063f0dbda9c8bac

Supplementary Figure 14 https://figshare.com/s/58a4c79c1b31b4474c1c

Supplementary Figure 15 https://figshare.com/s/5ccc12c8bf07a477ab19

Supplementary Figure 16 https://figshare.com/s/64b5c5461c87858830e5

Supplementary Figure 17 https://figshare.com/s/a950dec1eb4b32f8559a

Supplementary Movie 1 https://figshare.com/s/5f3e5147898632f68274

Supplementary Movie 2 https://figshare.com/s/cdf6d880d1a241c750f4

Supplementary Movie 3 https://figshare.com/s/5b6d3fb558aeb40771d2

Supplementary Movie 4 https://figshare.com/s/f2c26de3a7403d10148c

Supplementary Movie 5 https://figshare.com/s/03f83afad3ad6383aa20

Supplementary Movie 6 https://figshare.com/s/51aa1e42b4cb45ecdc60

Supplementary Movie 7 https://figshare.com/s/415c7431de05ecfc5449

Supplementary Movie 8 https://figshare.com/s/8a2169e10ff223bc971d

Supplementary Movie 9 https://figshare.com/s/a3aaebcb00ccb50f9b4d

Supplementary Movie 10 https://figshare.com/s/84ac6e35c61e47142339

Supplementary Movie 11 https://figshare.com/s/b87dfe1d640a284edf86

Supplementary Movie 12 https://figshare.com/s/f74f27f17b2cb7343559

Supplementary Movie 13 https://figshare.com/s/b9f1f922f31ada373b87

Supplementary Movie 14 https://figshare.com/s/9a1291cd30650944a05f

Supplementary Movie 15 https://figshare.com/s/3a08c958c980bb16c948

Supplementary Movie 16 https://figshare.com/s/5cd90d5c165d52dfa319

Supplementary Movie 17 https://figshare.com/s/3bac7c34e759d5849442

Supplementary Movie 18 https://figshare.com/s/db1b70cf18c9fe058a57

Supplementary Movie 19 https://figshare.com/s/769438c9b5a3cfade65d

