## Supplementary Information for "Early mechanisms of aortic failure in a zebrafish model for thoracic aortic dissection and rupture"

### SUPPLEMENTARY CASE PRESENTATION

The proband is an 11-year-old-boy, carried to term after an uncomplicated pregnancy to nonconsanguineous parents of Turkish origin. Evaluation at birth showed normal anthropometry. At the age of 4 months, he was referred to the clinical genetics department because of delayed neuromotor development and craniofacial dysmorphisms, including downslanted palpebral fissures, hypertelorism, a prominent nose with a wide nasal bridge, and large protruding ears. Inspection of the oral cavity revealed a highly arched palate. His skin was thin, hyperextensible, and he showed mild hyperkeratosis on his soles. Musculoskeletal features, including arachnodactyly, pectus excavatum, pes planum, and hyperlaxity of both small and large joints were present. Anthropometry at follow-up was normal. Cardiac examination with ultrasound revealed the presence of a bicuspid aortic valve. Magnetic resonance (MR) angiography at the age of 9 showed mild tortuosity of the vertebral arteries. There was also mild intellectual disability, and the neuromotor development was mildly delayed. MR imaging of the brain at age 11 was normal.

Genetic analysis with array comparative genomic hybridization (180k Agilent array) revealed a 15q22.31q23 deletion, inherited from his mother, encompassing 8 protein-coding genes, including *SMAD3*, *SMAD6*, *AAGAB.* The mother showed bicuspid aortic valve, mild but stable aortic root dilatation (Z-score 2.38), low normal intellectual capabilities, and craniofacial features similar to her son. Heterozygous pathogenic variants in *AAGAB* cause palmoplantar keratoderma[17] (**Supplementary Fig. 17**). Heterozygous loss-of-function PVs in *SMAD3* and *SMAD6* have been associated with LDS3[2] and BAV[12,13], respectively. His brother presented with isolated intellectual disability

(ID), but did not carry the 15q22.31q23 deletion. Exome sequencing was performed on both the proband and his brother but did not reveal an additional genetic cause for the ID in either of them. For the other affected genes, no gene-phenotype relations are currently described in OMIM (October 2023).

To our surprise, despite the deletion of both *SMAD3* and *SMAD6*, the cardiovascular phenotype seemed less pronounced in the 15q22.31q23 microdeletion syndrome than in patients with a pathogenic variant in *SMAD3* alone[9]. (**Supplementary Table 8**). We therefore hypothesized an attenuating effect of *SMAD6* on the *SMAD3* LOF phenotype.

### SUPPLEMENTARY METHODS

#### *Ethics statement*

This study was approved by the Animal Ethics Committee of the Ghent University Faculty of Medicine and Health Sciences (Permit number: ECD 14/70). Protocols adhered to the general guidelines, in agreement with EU Directive 2010/63/EU for laboratory animals, for zebrafish handling, mating, embryo collection and maintenance[23,24]. Informed consent to report clinical data was obtained from the parents of the patient.

#### *Zebrafish husbandry*

Zebrafish lines were housed at the zebrafish facility Ghent in semi-closed recirculating housing systems (ZebTEC and WTU systems, Tecniplast) at a constant temperature (27-28°C), pH (~7,5), conductivity (~550µS) and light dark cycle (14/10). Zebrafish were fed every day with dry food (Gemma Micro) and with artemia (Ocean Nutrition).

#### *Zebrafish lines: design and generation*

*Smad3a*$^{sa2363/+}$ was acquired from the Zebrafish International Research Center (ZIRC). Three different sgRNA molecules were designed for *smad3b, smad6a* and *smad6b,* respectively, using the CRISPRdirect software[25]. gBlocks (IDT) with the following structure were designed for the three target sites: 5′-CCGCTAGCTAATACGACTCACTATA–**GG**-**$N_{18}$**-GTTTTAGAGCTAGAAATAGCAAGTTAAAATAAGGCTAGTCCGTTATCAACTTGAAAAAGTGGCACCGAGTCGGTGCTTTT-3′, with $N_{18}$ representing the last 18 nucleotides of the protospacer, preceded by GG nucleotides (in bold), which assure optimal T7 polymerase-dependent *in vitro* transcription. Specific gBlocks for all target sites are listed in **Supplementary Table 1**. Lyophilized gBlocks were dissolved in nuclease-free water (10 ng/µL) and 4 µL was used as template for *in vitro* transcription with the MEGAshortscript™ T7 Transcription Kit (Invitrogen). To obtain maximum yield, incubation at 37°C was carried out overnight. No further alterations to the manufacturer's guidelines were made. The transcripts were purified using the MEGAclear™ Kit (Life Technologies) following manufacturer's procedures[26]. Zygotes with an AB background were injected in the nucleus during the single-cell stage with 250 pg of Cas9 wild type (WT) nuclease protein with nuclear localization signal (ToolGen). Identification of the founder zebrafish was performed according to the workflow previously described[27]. Genotyping was performed with primers and PCR protocols listed in **Supplementary Table 2**. All variants are described using Genome Reference Consortium Zebrafish Build 11 (GRCz11). All utilized zebrafish lines are described in **Supplementary Table 3**.

##### *Vasculogenesis in embryos*

Zebrafish were outcrossed to the transgenic *Tg(fli1:EGFP)* line in order to visualize endothelial cells. 5 or 10 days post fertilization (dpf) embryos were immobilized using 0.8% seaPlaque low melting agarose (Lonza) with 160 mg/mL ethyl 3-aminobenzoate methanesulfonate to sustain anesthesia. A Z-stack recording of the entire ventral aorta from the heart until the terminal branching point was taken using a ZEISS Axio Observer.Z1 microscope (Zeiss). Via ImageJ, the plugin "stack focuser" was used to create a focused image of the entire Z-stack. For each zebrafish, the projected normalized surface area, normalized projected length and normalized average projected diameter between aortic arch 3 (AA3) and aortic arch 4 (AA4) (calculated as the projected aortic area between aortic arch 3 and 4 divided by the aortic length of the segment) was measured as well as the normalized projected aorta diameter adjacent to the bulbus arteriosus. Normalization factors were obtained by dividing the head length (distance between tip of snout and attachment of the pectoral fin) of each zebrafish by the average head length (**Supplementary Fig. 1a-b)**. In all experiments, a cohort comprising both male and female zebrafish were utilized. All measurements were performed by an observer blinded to the genotype and fish were randomly selected. For the single knockout (SKO) experiments, mutants and controls were offspring of the same heterozygous parent pair. For the qKO and *smad3a/b* DKO experiments, mutants were compared to second degree related wild WT controls. For *smad6a/b* DKO, mutants were compared to the same WT as the qKO (**Supplementary Fig. 1c)**. Only samples in which no clear edge-detection of the ventral aorta was visible were excluded.

##### *3D reconstruction of the ventral aorta in adult zebrafish*

Adult zebrafish between 6 and 13 months post fertilization (mpf) were embedded in paraffin using the Leica EG1160 paraffin embedding station (Leica Biosystems). Serial 5 µm paraffin cross sections of the entire ventral aorta were taken using a Microm HM355S Automatic Rotary microtome (Thermofisher Scientific). After deparaffinization, sections were placed in Weigert's iron hematoxylin (Sigma-Aldrich) for 8 minutes and rinsed under running tap water for 5 minutes. Next, Resorcin-Fuchsin (Sigma-Aldrich) was added for one hour followed by two wash steps in 95% ethanol and one wash with reverse osmosis (RO) water. Slides were dehydrated and mounted with Entellan mounting medium (Merck).

Pictures were taken of serial sections with a ZEISS Axio Observer.Z1 microscope (Zeiss) and aligned via the TrakEM2 1.0[29-31], a plugin of ImageJ. 3D modelling software Mimics 24.0 (Materialise) was used to generate the 3D reconstructions of the ventral aorta.

##### Ultrasound analysis

Ultrasound analysis was performed on zebrafish 6-7 mpf. Handling of the zebrafish was performed as previously described[32]. The long axis (LAX) provided 2D ventricular dimensions such as projected surface area of the ventricle in diastole and systole, which was normalized to body surface area. The abdominal-cranial axis provided color Doppler and pulse-wave Doppler for cardiovascular measurements. All measurements were performed in Vevo Lab 5.5.0. In all experiments, a cohort comprising both male and female zebrafish were utilized. All measurements were performed by an observer blinded to the genotype and fish were randomly selected.

##### Acute net handling stress induction

Induction of handling stress was adapted from previously described procedures[33]. Briefly zebrafish of 6 mpf were captured in a net and left suspended in the air for 90 seconds, returned to the housing tank for 90 seconds and netted again for 90 seconds. After the procedure, the zebrafish were returned to the housing tank and swimming behavior was monitored, in particular if zebrafish preferred to swim closer to the surface.

##### Alizarin Red staining for mineralized bone

Alizarin red staining for mineralized bone was performed as previously described[34]. All alizarin red stainings were performed at 13 mpf, except for the qKO, which were stained after sudden death between 6 and 13 mpf.

Ectopic bone on the skeletal elements of the vertebral bodies was scored on adult zebrafish in lateral position. Only abdominal (precaudal) and caudal vertebrae were considered[35]. Ectopic bone was scored for the left neural arches and neural spine, left haemal arches and haemal spine, and the left part of the vertebral body.

Skull morphometrics were performed on images of the zebrafish heads in lateral position obtained using a Leica M165 FC Fluorescent Stereo Microscope. In ImageJ, the following lengths were measured: distance of the tip of the snout to the most caudal part of the supraoccipital bone, distance between tip of the snout to the middle of the eye, with the middle of the eye determined as the intersection of the horizontal and vertical maximal diameter, distance of the tip of the snout to the most anterior part of the frontal bone. All lengths were normalized using the diameter of the eye because eye diameter correlates with standard length **(Supplementary Fig. 1b)**[36]. Measurements of the surface area of the cranium were taken. The surface area of the frontal and parietal bone and the frontal plus parietal plus supraoccipital bone was compared between the genotypes. For both measurements, the surface area was normalized using standard length (tip of the snout to posterior tip of the notochord) as normalization factor. All obtained genotypes are the result of heterozygous incrosses of the parent zebrafish, except for WT, which were re-used as controls in all experiments. In all experiments, a cohort comprising both male and female zebrafish were utilized. All measurements were performed by an observer blinded to the genotype and fish were randomly selected.

##### *Transcriptome analysis*

RNA sequencing was performed on five pools, each containing five qKO or WT control cousins. For RNA extraction, the RNeasy® Mini kit (Qiagen) was used following manufacturers guidelines. Only samples with RNA integrity number (RIN) values higher than 9.8 were considered. Next generation sequencing libraries were prepared according to the manufacturer's procedures (TruSeq Stranded Total RNA kit from Illumina) prior to being sequenced on a NovaSeq6000 instrument (Illumina). Read quality was verified using FastQC v0.11.9[37] followed by adapter trimming with Trim Galore v0.6.6[38]. Trimmed reads were mapped to *Danio rerio* genome version 11 using STAR v2.7.6a[39]. FeatureCounts package from SubRead v2.0.0 was used for count estimation per gene[42]. Duplication levels were evaluated using the DupRadar package[40] along with additional control statistics provided by MultiQC v1.9[41]. Count estimates were analyzed in R v4.1.1. An unsupervised multidimensional scaling plot was created via the EdgeR package[43]. Distances among samples in this plot approximate the log2 fold changes between each pair, based on the expression (counts per million) of the top 500 genes. Low expressed genes were removed using the filterByExpr function (edgeR), retaining genes with sufficiently large counts to be included in the statistical analysis[44]. Differential expression analysis was performed using negative binomial models followed by a quasi-likelihood (QL) F-test implemented in the edgeR package. P-values were adjusted using the Benjamini–Hochberg method and the false discovery rate (FDR) was controlled at 0.05 for the set of differentially expressed genes (DEG) A fold change threshold of 2 was used to select the relevant genes. Gene set enrichment analysis (GSEA) was applied for the DEG list using clusterProfiler and the enrichplot R packages[45]. The enriched terms in Gene Ontology (GO) and Kyoto Encyclopedia of Genes and Genomes (KEGG) databases were identified via the gseGO and gseKEGG functions, with a p-value significance threshold at 0.05.

##### Synchrotron phase contrast micro Computed tomography imaging

Propagation-based phase-contrast synchrotron X-ray imaging was performed at the TOMCAT (X02DA) beamline of the Swiss Light Source (Paul Scherrer Institute in Villigen, Switzerland) as previously described[46]. In short, 8 mpf zebrafish heads were positioned upright immobilized in agarose in a 1.5 mL Eppendorf tube. The tube was mounted to the robot sample holder using a specially designed 3D printed mold. Beamline settings were as follows: 21.8 keV monochromatic beam energy, 250 mm sample-detector distance, 1501 projections, 10 darks and 2 series of 50 flats where collected using a LUAg(Ce) 20 µm scintillator, 4x objective combined with a PCO.Edge 5.5 sCMOS camera leading to an effective pixel size of 1.625um. Tomographic reconstruction was performed using the Gridrec algorithm after applying the Paganin phase retrieval method[47,48].

##### 3D Biomechanical modeling to assess principal stress at systolic peak in the aortic wall

In a previous study[46], zebrafish-specific 3D biomechanical fluid-structure interaction models of five different 13-month-old zebrafish were developed. These WT geometries and inlet blood profiles were used as a starting point to resolve the hemodynamics and the mechanical stresses within the vessel wall of qKO mutants. Almost all parameter settings as described in[46] were maintained but to mimic the vascular organization found in qKO mutants, flow through right aortic arch 2, 3 and 4 was set to nearly zero.

##### Transmission electron microscopy

WT and qKO zebrafish of 16 mpf were euthanized and prefixed for 2 h in a vacuum chamber at room temperature followed by overnight incubation at 4°C in Karnovsky fixative (2.5% glutaraldehyde and 2% paraformaldehyde in 0.05 M sodium cacodylate buffer (pH 7.4)). Next, samples were decalcified for three weeks with decalcification buffer at 4°C (2.5% glutaraldehyde and 2% paraformaldehyde in 0.05 M sodium cacodylate buffer (pH 7.4), 0.1 M EDTA). Solution was refreshed every two days. Post-fixation for 2 h in 1% osmium tetroxide, 0.1M cacodylate buffer with 8% saccharose and 0.004% $CaCl_2$ was followed by *en bloc* staining for 1 h in 1% aqueous uranyl acetate. After dehydration in ethanol and isopropanol series, the samples were embedded in Spurr's resin. 80 nm sections were stained with uranyl acetate and lead citrate and examined by TEM (JEM 1010, JEOL) equipped with a CCD side-mounted Veleta camera (EMSIS).

##### ***Swimming behavior studies***

Swimming behavior of 9 mpf WT, *smad3a*$^{+/-}$;*smad3b*$^{-/-}$;*smad6a*$^{+/-}$;*smad6b*$^{-/-}$ and qKO zebrafish was analyzed in a custom-made dark observation chamber (Noldus) equipped with the Basler GenICam capturing 25 frames/second for top-view image acquisition. Data was analyzed using the EthoVision XT 17 software (Noldus). Zebrafish were placed individually in a tank (16 cm (L) x 10 cm (W) x 8 cm (H)) with system water and could acclimatize for 10 minutes before a 10-minute test period. The complete experiment was performed in the dark. For each zebrafish, nose-point, center-point, and tail-base detection was performed in the EthoVision XT 17 software. Movement is depicted as total distance travelled (in cm) during the 10-minute test period, normalized for standard length of each zebrafish. Movement frequency was also recorded during the 10-minute test period. In all experiments, a cohort comprising both male and female zebrafish were utilized. All measurements were performed by an observer blinded to the genotype.

##### *1-phenyl-2-thiourea treatment*

At 1 dpf, 0.003% 1-phenyl-2-thiourea treatment (PTU) in E3 medium was administered to WT zebrafish. The medium was changed daily until 10 dpf. Starting from 5 dpf, zebrafish were given dry food for 2 hours daily, after which the medium was refreshed again.

### SUPPLEMENTARY FIGURES

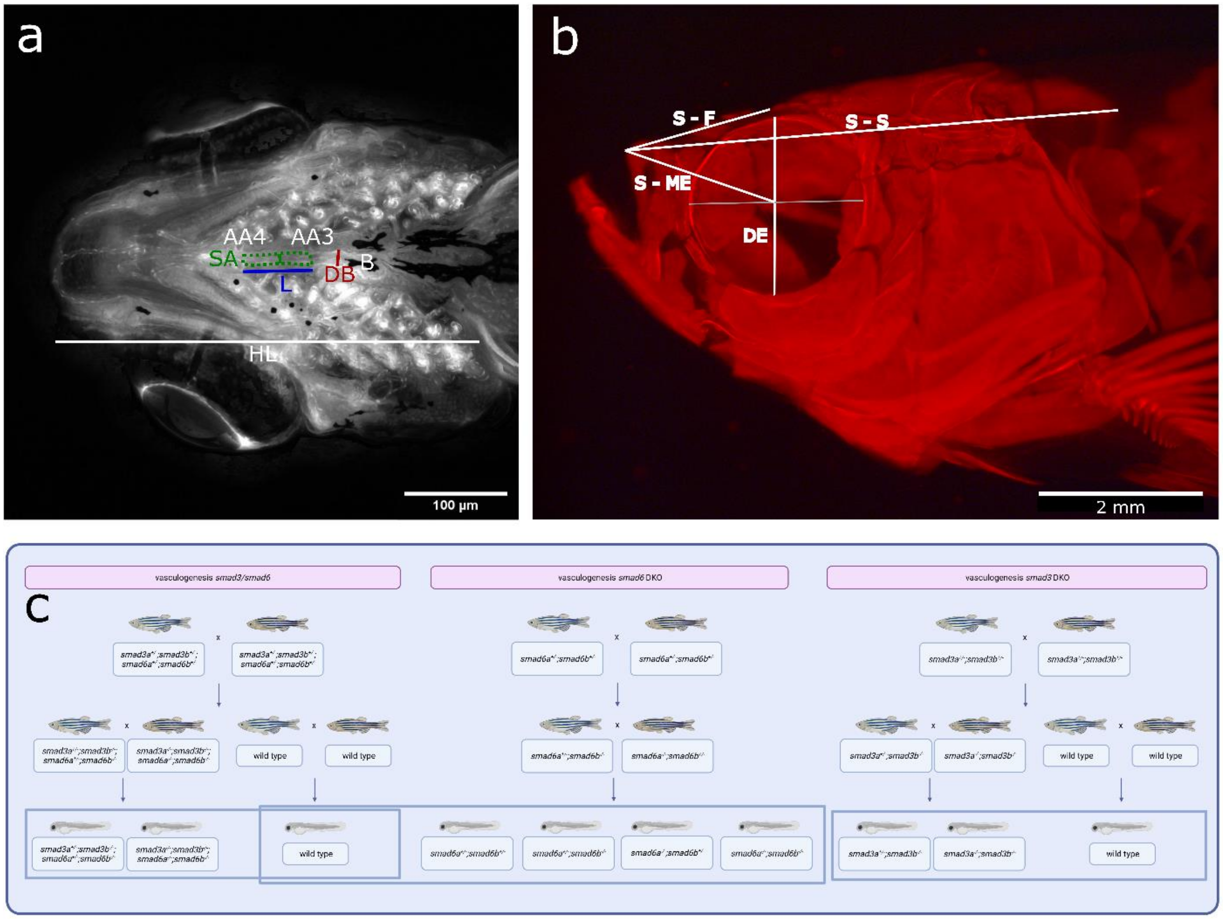


**Supplementary Figure 1: Overview of the measurements to evaluate vasculogenesis and skull morphometrics experiments and breeding scheme. (a)** Ventral view of the dorsal aorta at 10 dpf Legend: “B” bulbus arteriosus, “AA3” aortic arch 3, “AA4” aortic arch 4, “DB” diameter adjacent to the bulbus arteriosus (red), “SA” projected surface area of the ventral aorta between aortic arch 3 and 4 (green dashed line), aorta diameter (green line), “L” length of the aortic segment between aortic arch 3 and 4 (blue). “HL” head length (white). **(b)** Lateral view of an alizarin red staining for mineralized bone of the zebrafish skull at 13 mpf. Legend: “S-F” distance between tip of the snout and tip of the frontal bone, “S-S” distance between the tip of the snout and the most caudal point of the supraoccipital bone, “S-ME” distance between tip of the snout and middle of the eye. “DE” diameter of the eye. **(c)** Breeding scheme utilized for vasculogenesis experiments at 10 dpf. Created with Biorender.com.

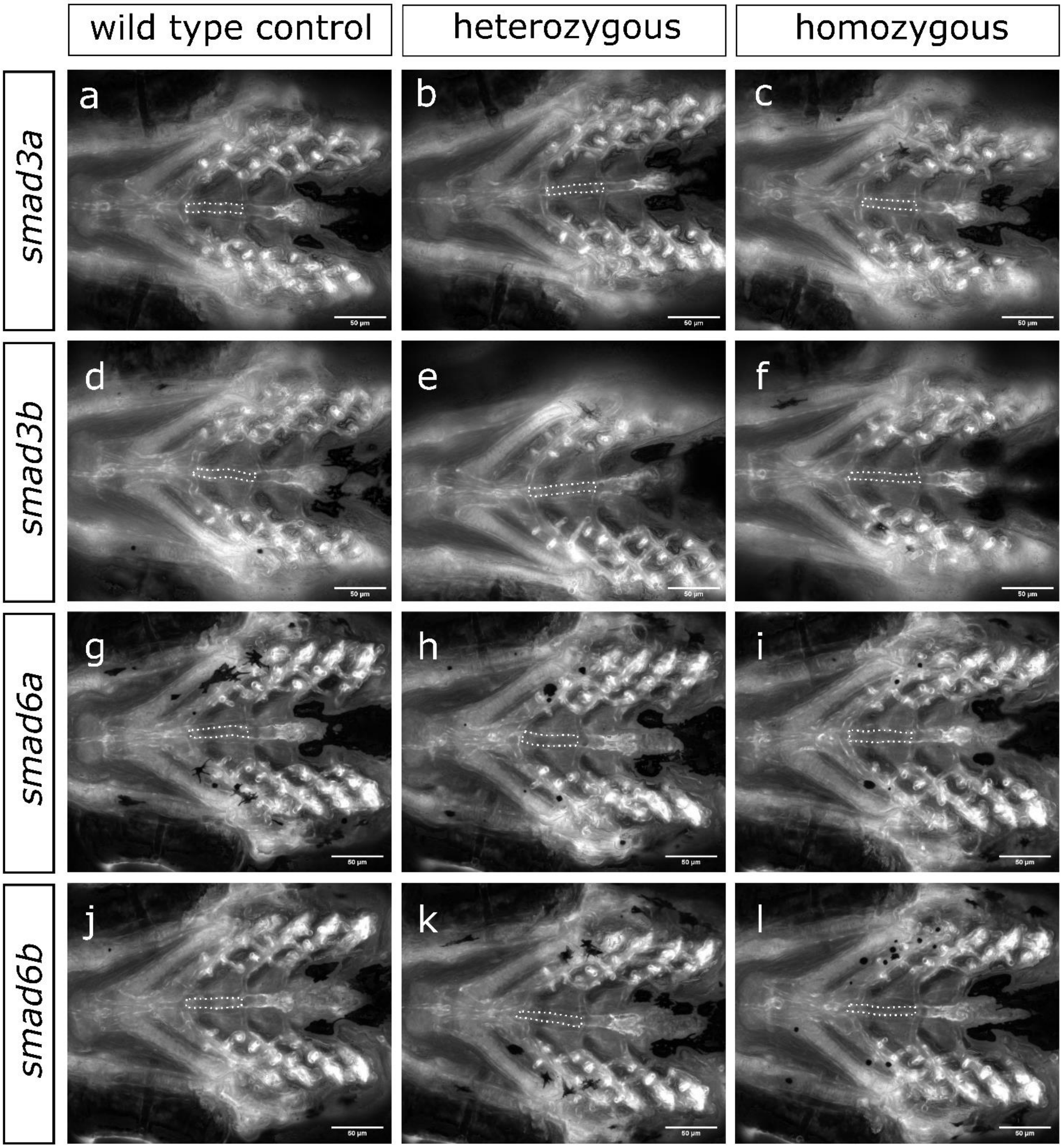
wild type control
heterozygous
homozygous
smad3a
smad3b
smad6a
smad6b
a
b
c
d
e
f
g
h
i
j
k
l

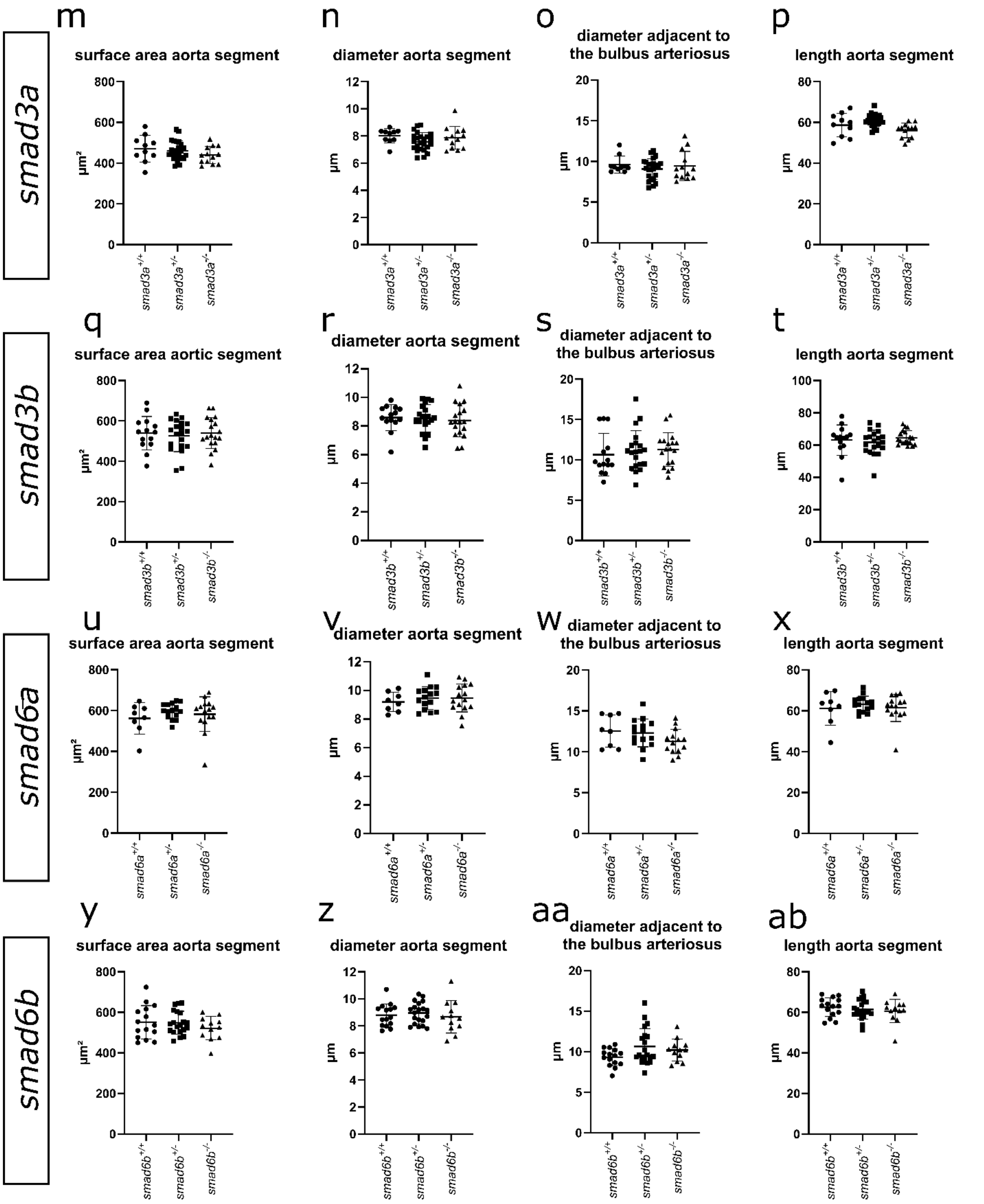


**Supplementary Figure 2: Vasculogenesis of 5 dpf single *smad3a*$^{-/-}$, *smad3b*$^{-/-}$, *smad6a*$^{-/-}$ or *smad6b*$^{-/-}$ knockouts. (a - l)** Ventral view of the cardiovascular system at 5

dpf of single *smad* knockout lines crossed with the *Tg(fli1:EGFP)* endothelial reporter line. Projected surface area of the ventral aorta between aortic arch 3 and 4 is indicated (white dashed line). **(m – ab)** No significant differences are detected in vasculogenesis of single *smad3a*$^{-/-}$ (n=10-24-13), *smad3b*$^{-/-}$ (n=14-20-18), *smad6a*$^{-/-}$ (n=8-15-15) or *smad6b*$^{-/-}$ (n=15-20-12) knockouts. **(a-l)** Stack focused Z-stack pictures obtained with ZEISS Axio Observer.Z1 microscope. **(m-o,q– z, ab)** One-way ANOVA with Dunnett's multiple comparison test against WT controls (m-o,q– z, ab) or Brown-Forsythe and Welch ANOVA with Dunnett's T3 multiple comparison test against WT controls **(p,aa)**. Data represented as average ± standard deviation.

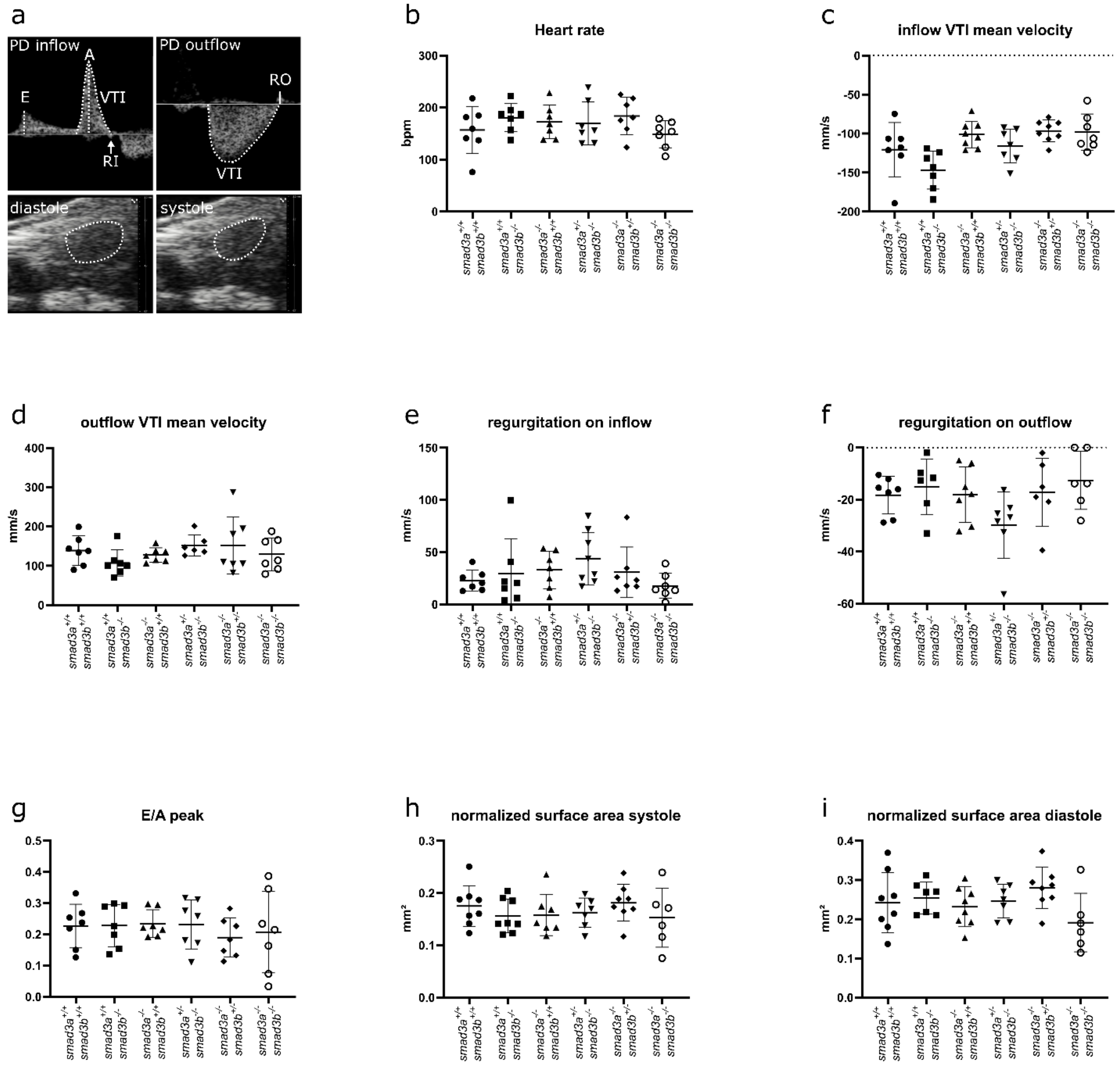


**Supplementary Figure 3: Ultrasound analysis of *smad3* mutant models at 6-7 mpf.** **(a)** Overview of pulsed wave doppler of the inflow (top left) and outflow (top right). Overview of B-mode imaging of ventricle in diastole (bottom left) and systole (bottom right). Ventricle is outlined with a dashed line . Legend: “E” early inflow peak, “A” Atrial peak, “VTI” velocity time integral, “RI” regurgitation peak inflow (white arrow), “RO” regurgitation peak outflow. “PD” pulsed wave doppler, “bpm” beats per minute. **(b-i)** No significant differences have been detected for heart rate (in beats per minute) (n=7-7-7-7-7-7), average inflow (n=7-7-8-7-8-7) and outflow (n=7-7-7-7-6-7) velocity, regurgitation inflow (n=7-7-7-8-7-7), regurgitation outflow (n=7-6-7-7-6-6), E/A peak (n=7-7-7-7-7-7) and normalized surface area of the ventricle in systole (n=8-8-7-7-8-6) or diastole (n=8-7-8-7-8-6). One-way ANOVA with Dunnett's multiple comparison test against WT controls (*smad3a*$^{+/+}$;*smad3b*$^{+/+}$). Data represented as average ± standard deviation.

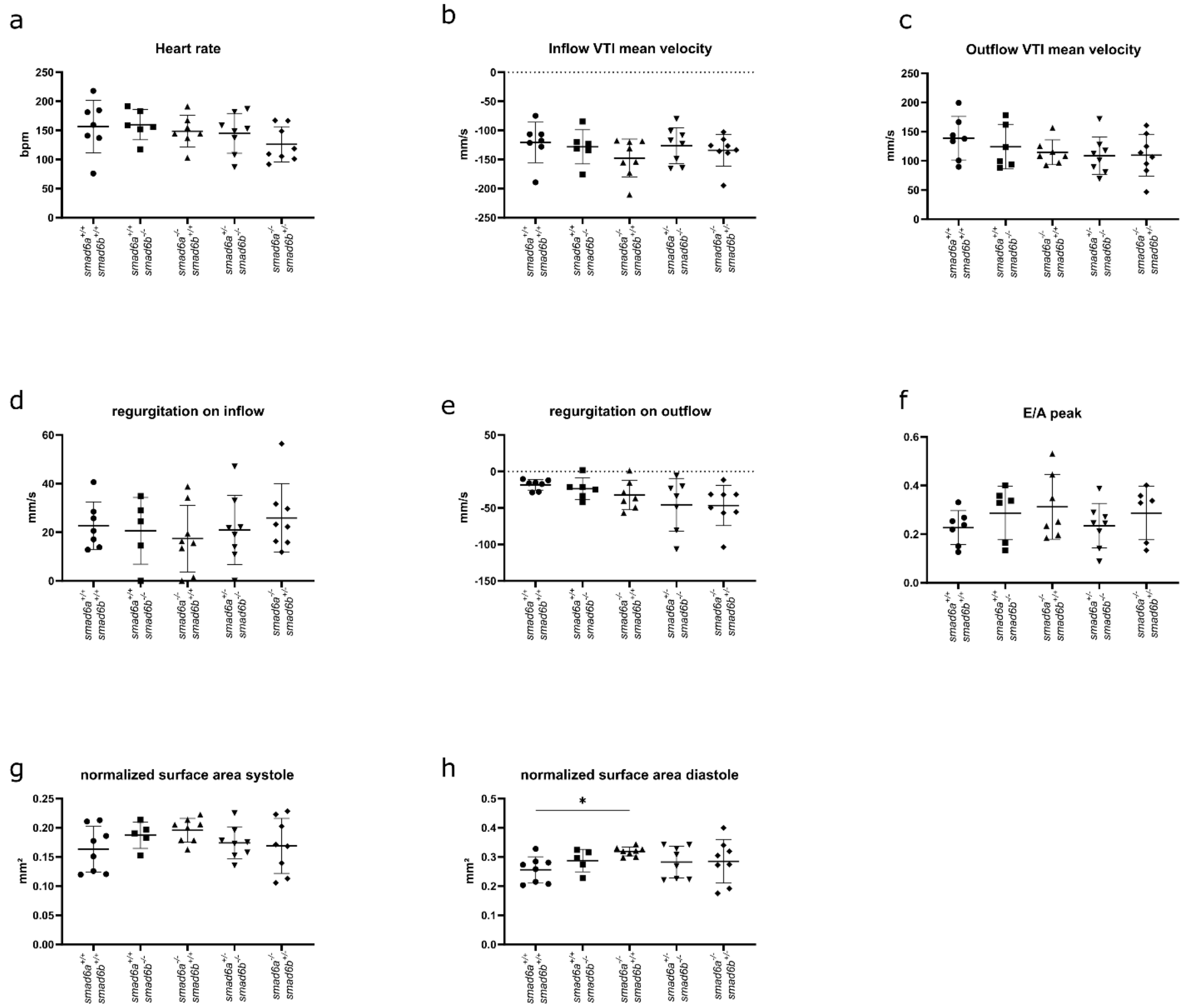


**Supplementary Figure 4: Ultrasound analysis of *smad6* mutant models at 6-7 mpf. (a-g)** No significant differences have been detected for heart rate (n=7-6-7-8-8), average inflow (n=7-6-8-8-8) and outflow (n=7-6-7-8-8) velocity, regurgitation inflow (n=7-6-8-8-8), regurgitation outflow (n=7-6-7-7-8) E/A peak (n=7-6-8-8-7) and normalized surface area of the ventricle in systole (n=8-5-8-8-8). **(h)** Normalized surface area of the ventricle in diastole (n=8-5-8-8-8) is increased for *smad6a*$^{-/-}$;*smad6b*$^{+/+}$.**(a-g)** One-way ANOVA with Dunnett's multiple comparison test against WT controls. **(h)** Brown-Forsythe and Welch ANOVA with Dunnett's T3 multiple comparison test against WT controls (*smad6a*$^{+/+}$;*smad6b*$^{+/+}$). Asterisks indicate significant differences. *p<0.05. Data represented as average ± standard deviation.

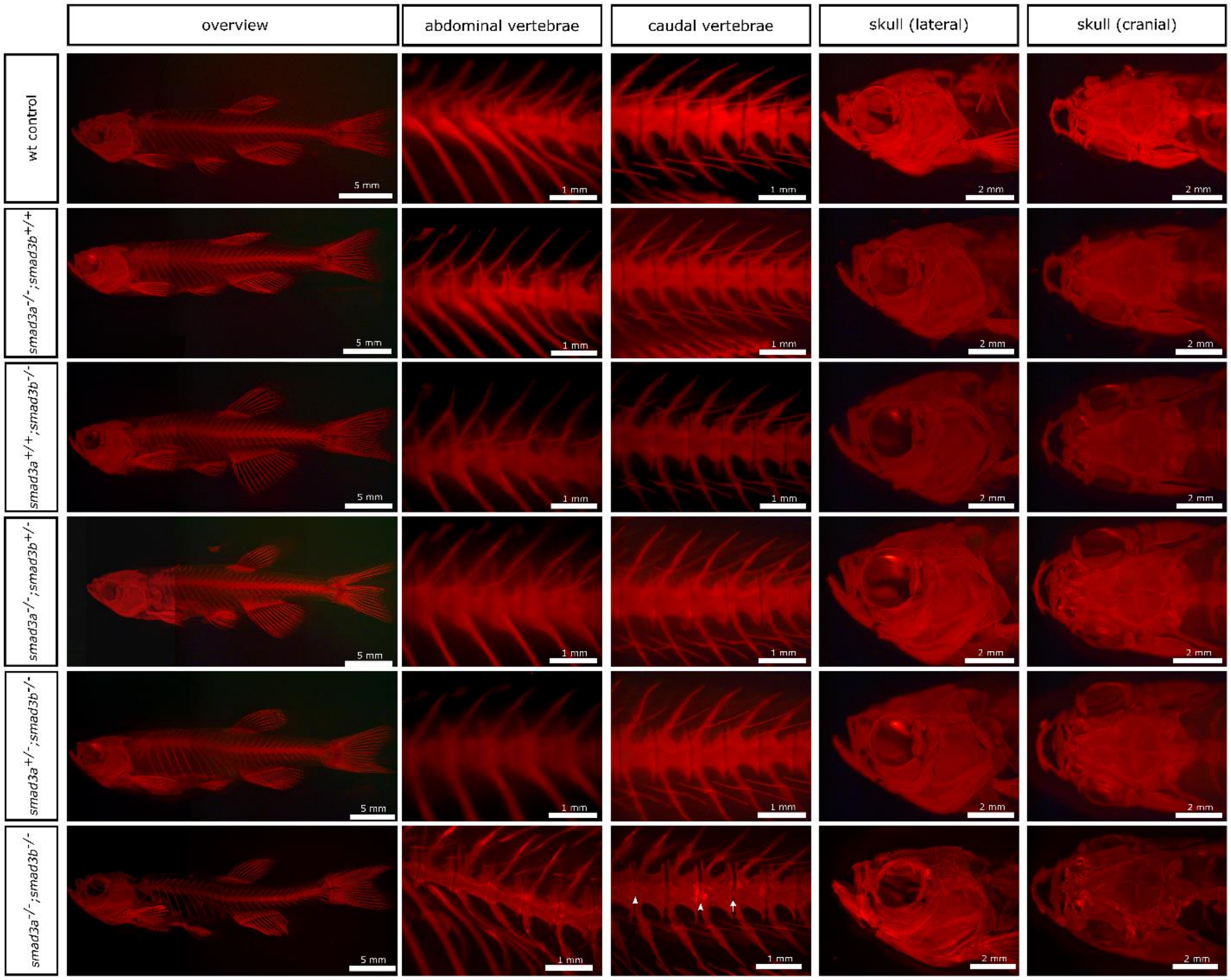


**Supplementary Figure 5: skeletal overview 13 mpf *smad3* knockout zebrafish.** Alizarin red staining for mineralized bone of a lateral overview picture, lateral view of abdominal and caudal vertebrae, and lateral and dorsal view of the skull, respectively of all *smad3* mutants. Notochord mineralization is indicated with a white arrowhead. Notochord sheet mineralization is indicated with a white arrow. Whole-mount alizarin red staining for mineralized bone pictures taken with Leica M165 FC Fluorescent Stereo Microscope.

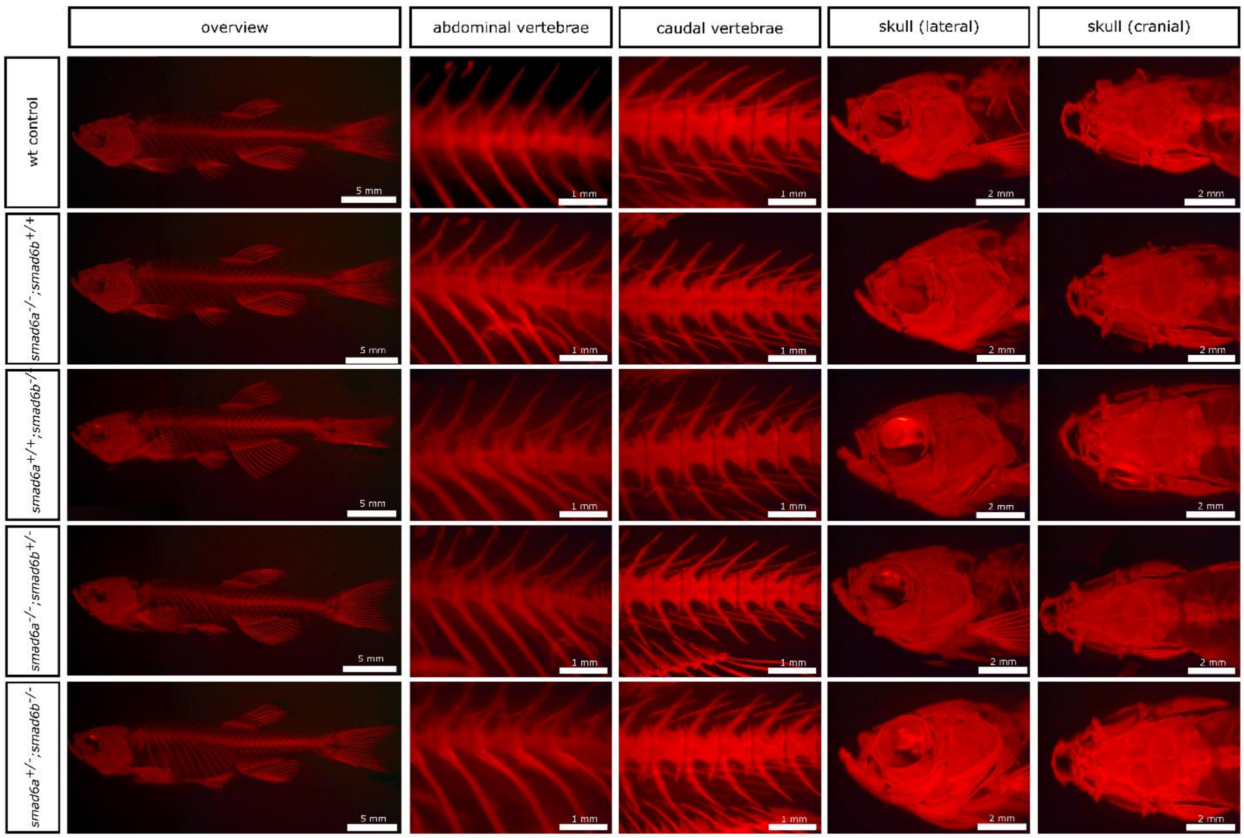


**Supplementary Figure 6: skeletal overview 13 mpf *smad6* knockout zebrafish.** Alizarin red staining for mineralized bone of a lateral overview picture, lateral view of abdominal and caudal vertebrae, and lateral and dorsal view of the skull, respectively of all viable *smad6* mutants. Whole-mount alizarin red staining for mineralized bone pictures taken with Leica M165 FC Fluorescent Stereo Microscope.

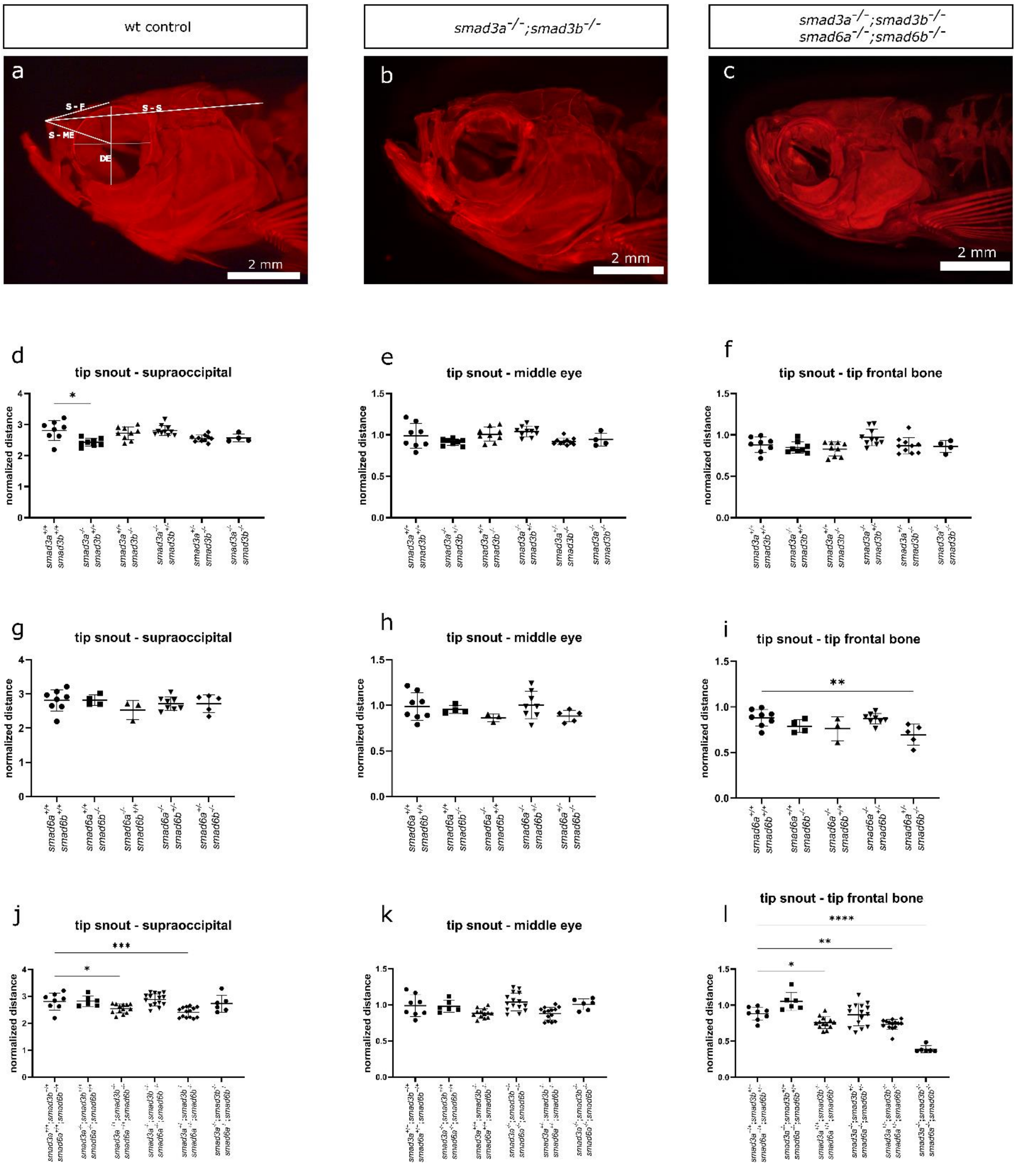


**Supplementary Figure 7: craniofacial aberrations in *smad3* and *smad6* knockout zebrafish. (a-c)** Overview of measured lengths shown on WT of 13 mpf and qKO between 8 and 13 mpf. Legend: "S-F" distance between tip of the snout and tip of the frontal bone.

“S-S” distance between tip of the snout and back of the supraoccipital bone. “S-ME” distance between tip of the snout and middle of the eye. “DE” diameter of the eye which was used as normalization factor. **(d-l)** Quantification of different lengths in the viable *smad3* (n=8-8-9-10-10-4), *smad6* (n=8-4-3-8-5) and *smad3/smad6* (n=8-6-13-15-14-6) genotypes. **(a-c)** Whole-mount alizarin red staining for mineralized bone pictures taken with Leica M165 FC Fluorescent Stereo Microscope. **(f-j)** One-way ANOVA with Dunnett’s multiple comparison test against WT controls. **(d,e,k,l)** Brown-Forsythe and Welch ANOVA with Dunnett’s T3 multiple comparison test against WT controls. Asterisks indicate significant differences. *p<0.05, **p<0.01, ***p<0.001, ****p<0.0001. Data represented as average ± standard deviation.

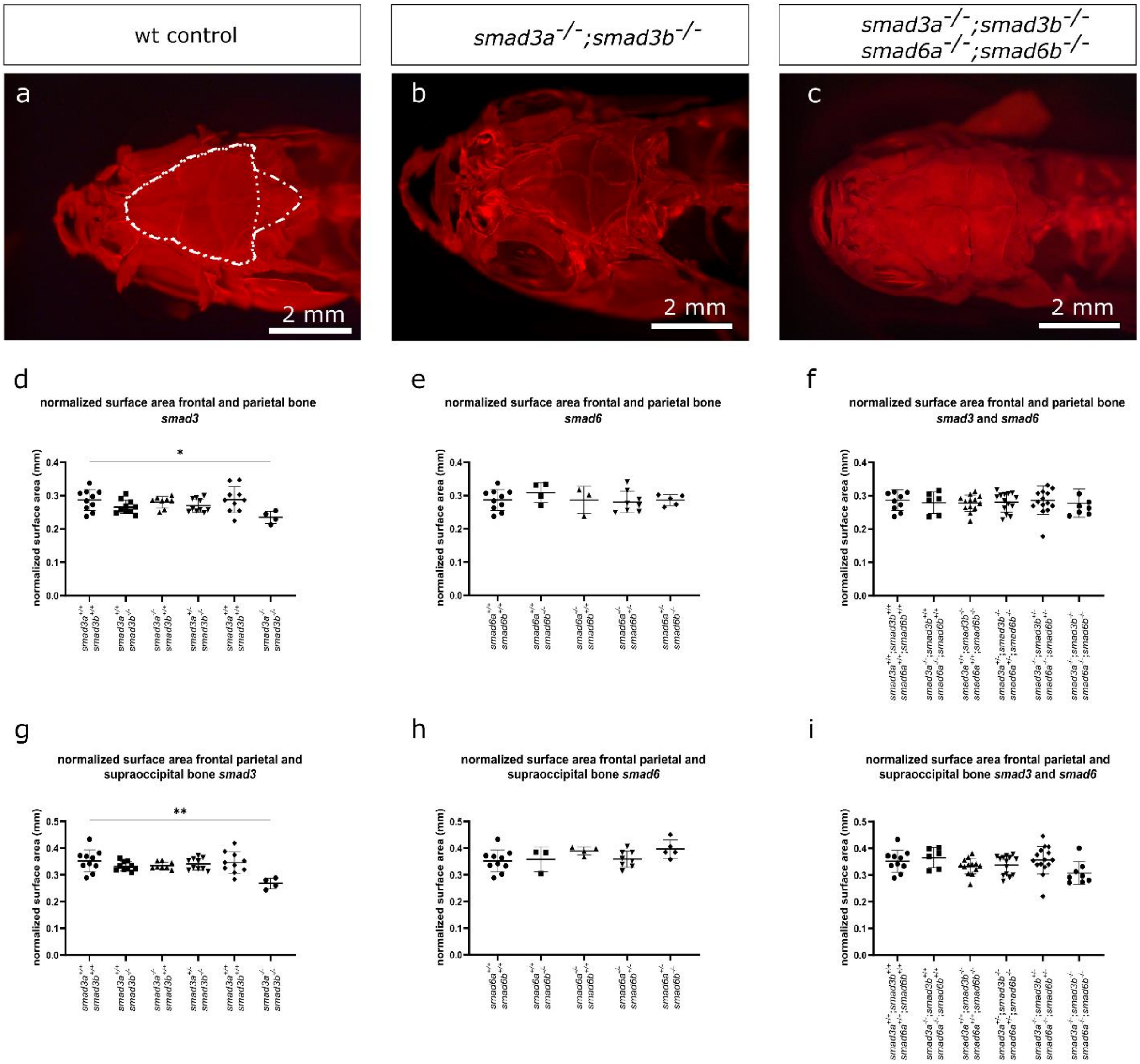


**Supplementary Figure 8: Reduction of cranial surface area in *smad3a*$^{-/-}$*;smad3b*$^{-/-}$ double knockout zebrafish at 13 mpf. (a-c)** Overview of the cranial roof of WT control, *smad3* (n=10-10-8-10-10-4), *smad6* (n=10-4-3-8-5) and *smad3/smad6* (n=10-6-13-14-15-8) genotypes. For the qKO zebrafish between 8 and 13 mpf are imaged (13 mpf is shown in c). Circumference of the frontal and parietal bone has been indicated with a dotted line. Circumference of the frontal, parietal and supraoccipital bone has been indicated with a dash-dotted line. **(d, g)** *smad3a*$^{-/-}$*;smad3b*$^{-/-}$ DKO show a significant reduction in normalized cranial roof surface area. This reduction is not observed in any other combination of genotypes. **(a-c)** Whole-mount alizarin red staining for mineralized bone pictures taken with Leica M165 FC Fluorescent Stereo Microscope. **(d, e, f, h, i)** One-way ANOVA with Dunnett's multiple comparison test against WT controls. **(g)** Brown-Forsythe and Welch ANOVA test with Dunnett's T3 multiple comparison test against WT controls. Asterisks indicate significant differences. *$p<0.05$, **$p<0.01$. Data represented as average ± standard deviation.

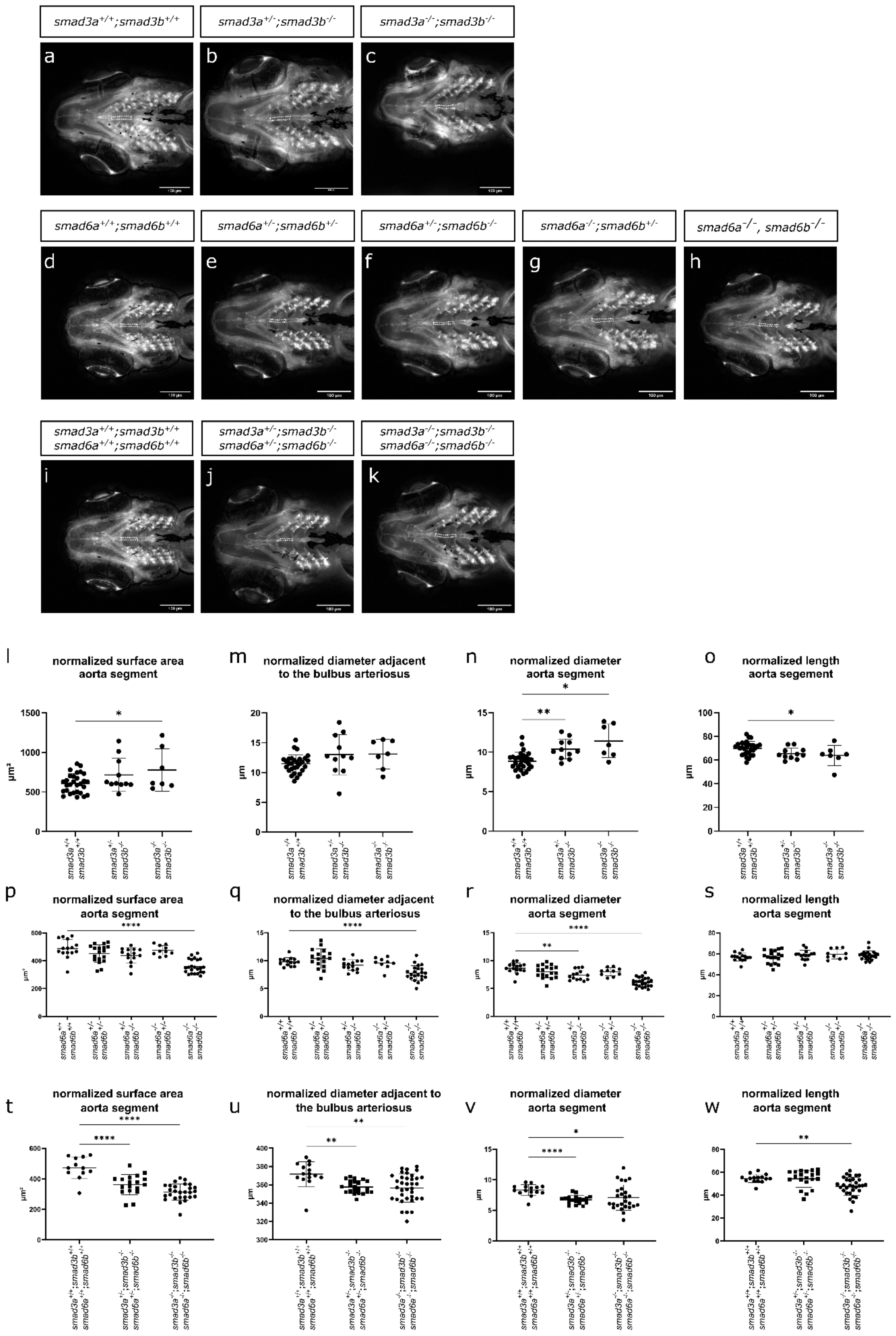
smad3a+/+;smad3b+/+
smad3a+/-;smad3b-/-
smad3a-/-;smad3b-/-
smad6a+/+;smad6b+/+
smad6a+/-;smad6b+/-
smad6a+/-;smad6b-/-
smad6a-/-;smad6b+/-
smad6a-/-, smad6b-/-
smad3a+/+;smad3b+/+ smad6a+/+;smad6b+/+
smad3a+/-;smad3b-/- smad6a+/-;smad6b-/-
smad3a-/-;smad3b-/- smad6a-/-;smad6b-/-
normalized surface area aorta segment
normalized diameter adjacent to the bulbus arteriosus
normalized diameter aorta segment
normalized length aorta segment
µm²
µm

**Supplementary Figure 9: Overview of vasculogenesis in double and qKO zebrafish.** **(a-k)** Ventral view of the cardiovascular structures at 10 dpf of *smad* knockout lines crossed with a *Tg(fli1:EGFP)* reporter line. Projected surface area of the ventral aorta between aortic arch 3 and 4 is indicated with a white dashed line. **(l-w)** quantification of measured areas and lengths in *smad3* (n=29-11-7)*, smad6* (n=15-18-1410-23) and *smad3/smad6* (n=15-22-36) knockouts. **(a-k)** Stack focused Z-stack pictures obtained with ZEISS Axio Observer.Z1 microscope. **(l,o,p,q,r,s,t,w)** One-way ANOVA with Dunnett's multiple comparison test against WT controls. **(m,n,u,v)** Brown-Forsythe and Welch ANOVA with Dunnett's T3 multiple comparison test against WT controls. Asterisks indicate significant differences. *p<0.05, **p<0.01, ***p<0.001, ****p<0.0001. Data represented as average ± standard deviation.

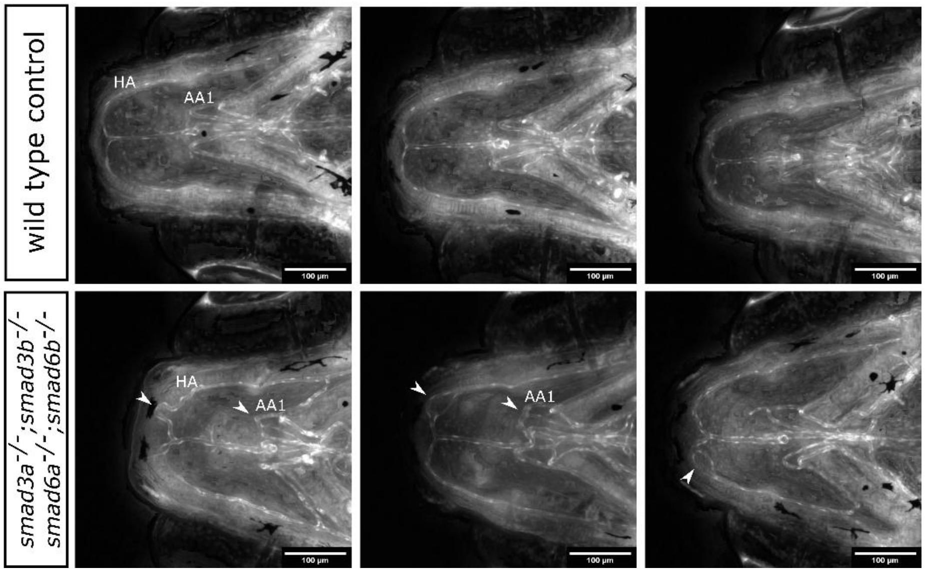


**Supplementary Figure 10: Arterial tortuosity and irregular branching in the hypobranchial artery and aortic arch 1 of qKO zebrafish at 10 days post fertilization.** Tortuosity and irregular branching are indicated with a white arrowhead. Legend: “HA” hypobranchial artery, “AA1” aortic arch 1. Stack focused Z-stack pictures obtained with ZEISS Axio Observer.Z1 microscope.

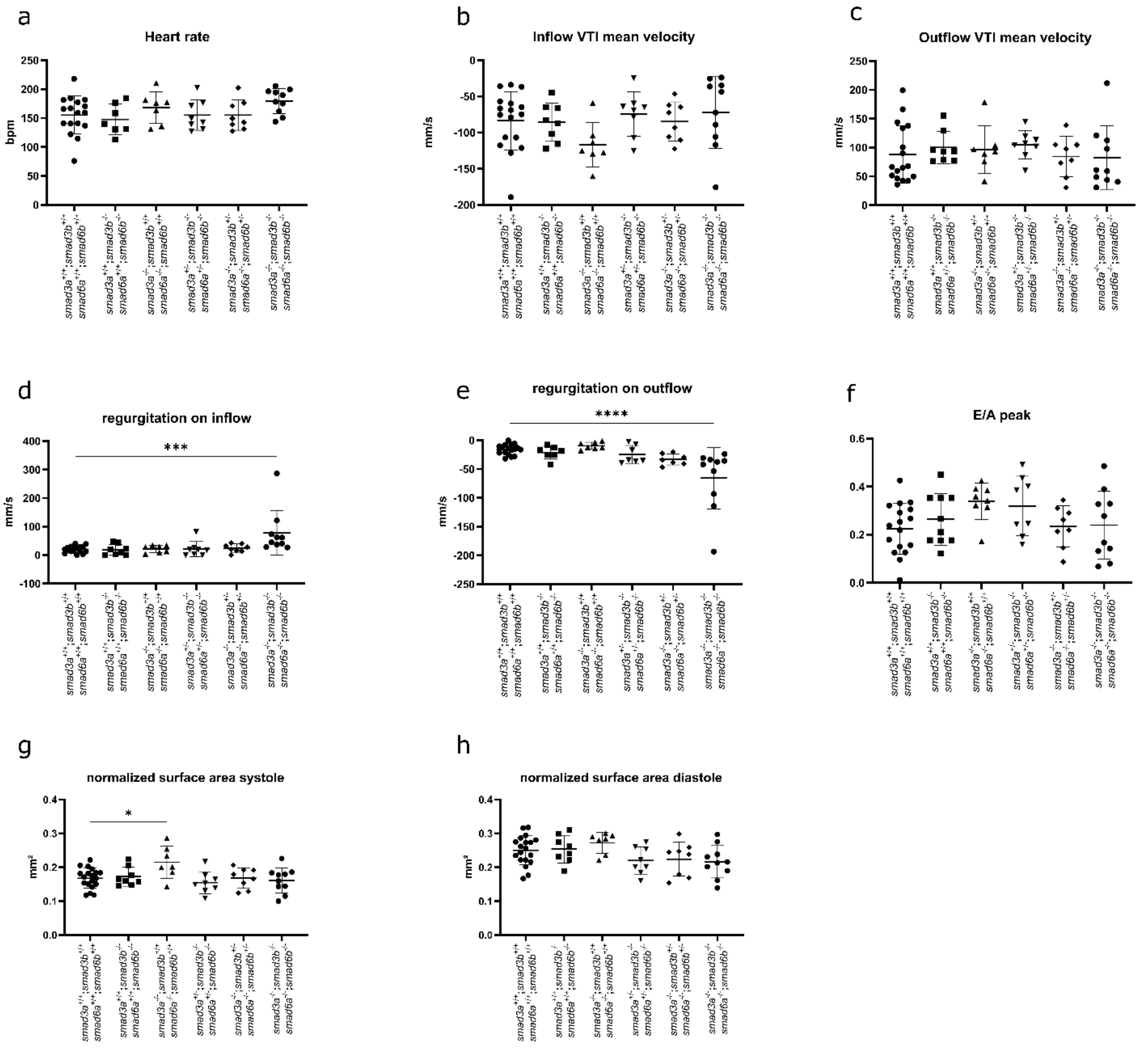


**Supplementary Figure 11: Ultrasound analysis in *smad3;smad6* mutant zebrafish models at 6-7 mpf. (a-h)** Quantification of ultrasound measurements in *smad3/smad6* mutant zebrafish. No significant differences are detected in heart rate (n=17-7-7-8-8-10), mean inflow velocity (n=17-8-7-8-8-10), mean outflow velocity (n=17-8-7-8-8-10), E/A peak (n=17-10-8-8-8-10) and normalized surface area in diastole (n=19-8-7-8-8-10). **(d, e)** Significant increase of regurgitation observed in the qKO model during inflow (n=17-8-

7-8-8-10) as well as outflow (n=17-7-77-7-11). **(g)** A small increase of normalized surface area of the ventricle in systole (n=19-8-7-8-8-10) is detected for *smad3a*$^{-/-}$;*smad6a*$^{-/-}$. **(a-h)** One-way ANOVA with Dunnett's multiple comparison test against WT controls (*smad3a*$^{+/+}$;*smad3b*$^{+/+}$;*smad6a*$^{+/+}$;*smad65*$^{+/+}$). Asterisks indicate significant differences. *p<0.05, ***p<0.001, ****p<0.0001. Data represented as average ± standard deviation.

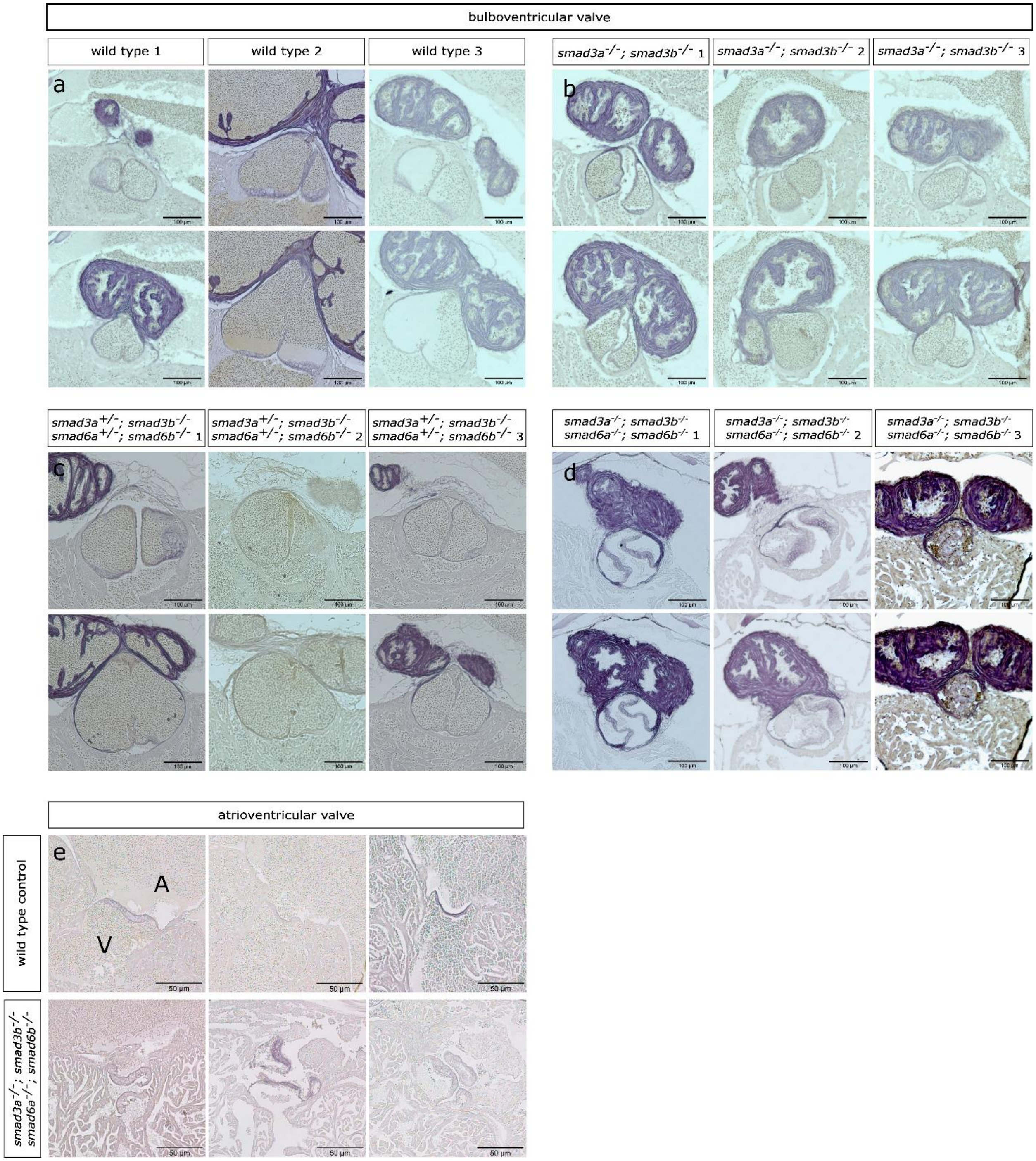


**Supplementary Figure 12: Hypertrophy in atrioventricular and bulboventricular valve in the qKO zebrafish model at 6 mpf. (a-d)** Overview of resorcin-Fuchsin stained sections of the bulboventricular valve from 3 WT zebrafish, 3 *smad3a*$^{-/-}$;*smad3b*$^{-/-}$, 3 *smad3a*$^{+/-}$;*smad3b*$^{-/-}$;*smad6a*$^{+/-}$;*smad6b*$^{-/-}$ and 3 qKO zebrafish. In the qKO model, the bulboventricular valve shows hypertrophy of the valve interstitial cells. **(e)** Hypertrophy of the valve interstitial cells and altered morphology of the valve leaflets is apparent in atrioventricular valves of the qKO zebrafish. **(a,e)** “A” atrium, “V” ventricle. Pictures obtained with ZEISS Axio Observer.Z1 microscope.

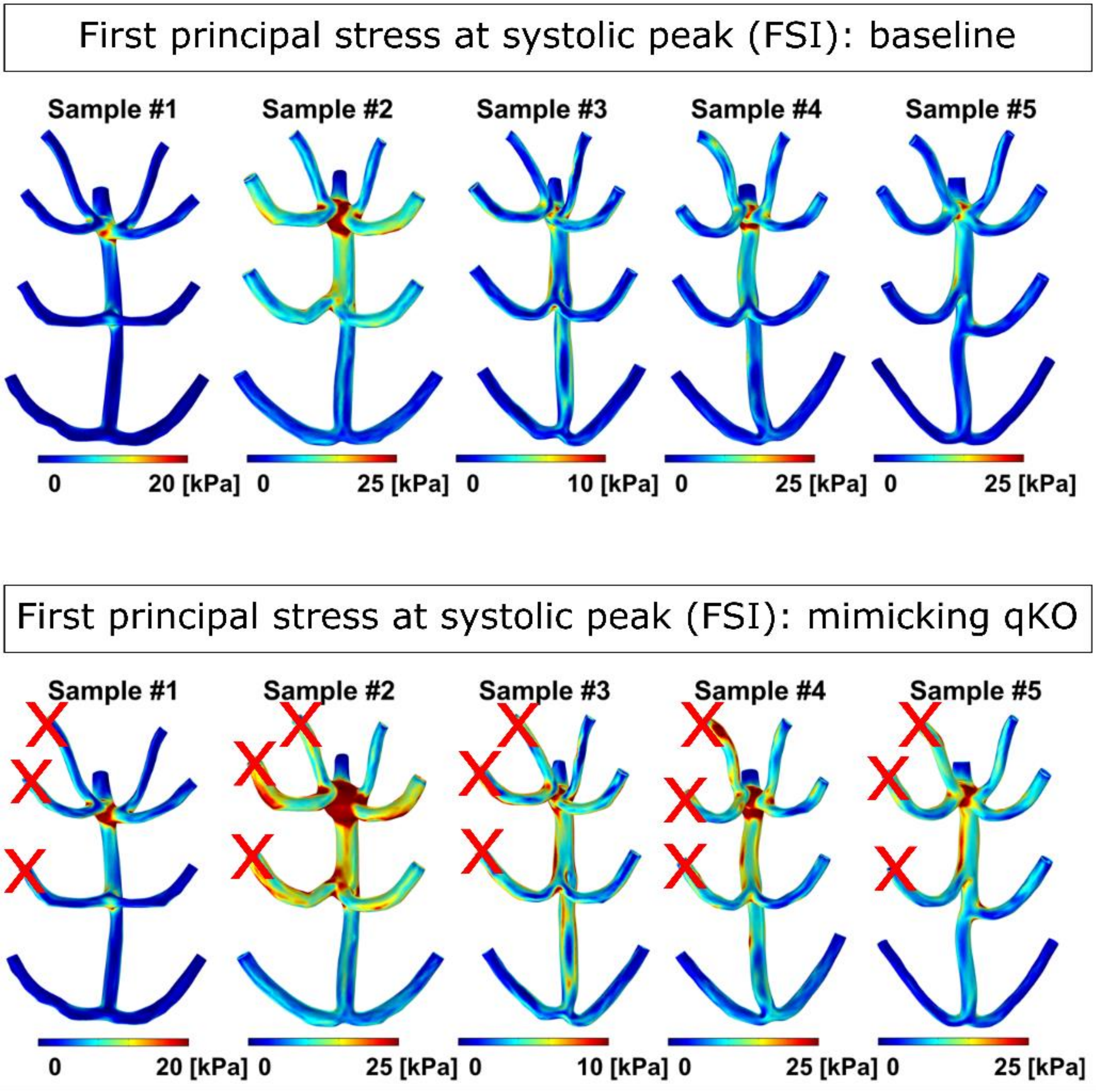


**Supplementary Figure 13: Modeling principal stress at systolic peak in qKO zebrafish.** First, principal stress at systolic peak is determined in wild type ventral aorta. In simulations corresponding to blood flow in qKO, blood flow of the right aortic arches 2, 3 and 4 is reduced to nearly 0 (indicated by red x) to mimic qKO aortic morphology. The principal stress at systolic peak in qKO simulations is mostly increased close to the branching points compared to WT simulations.

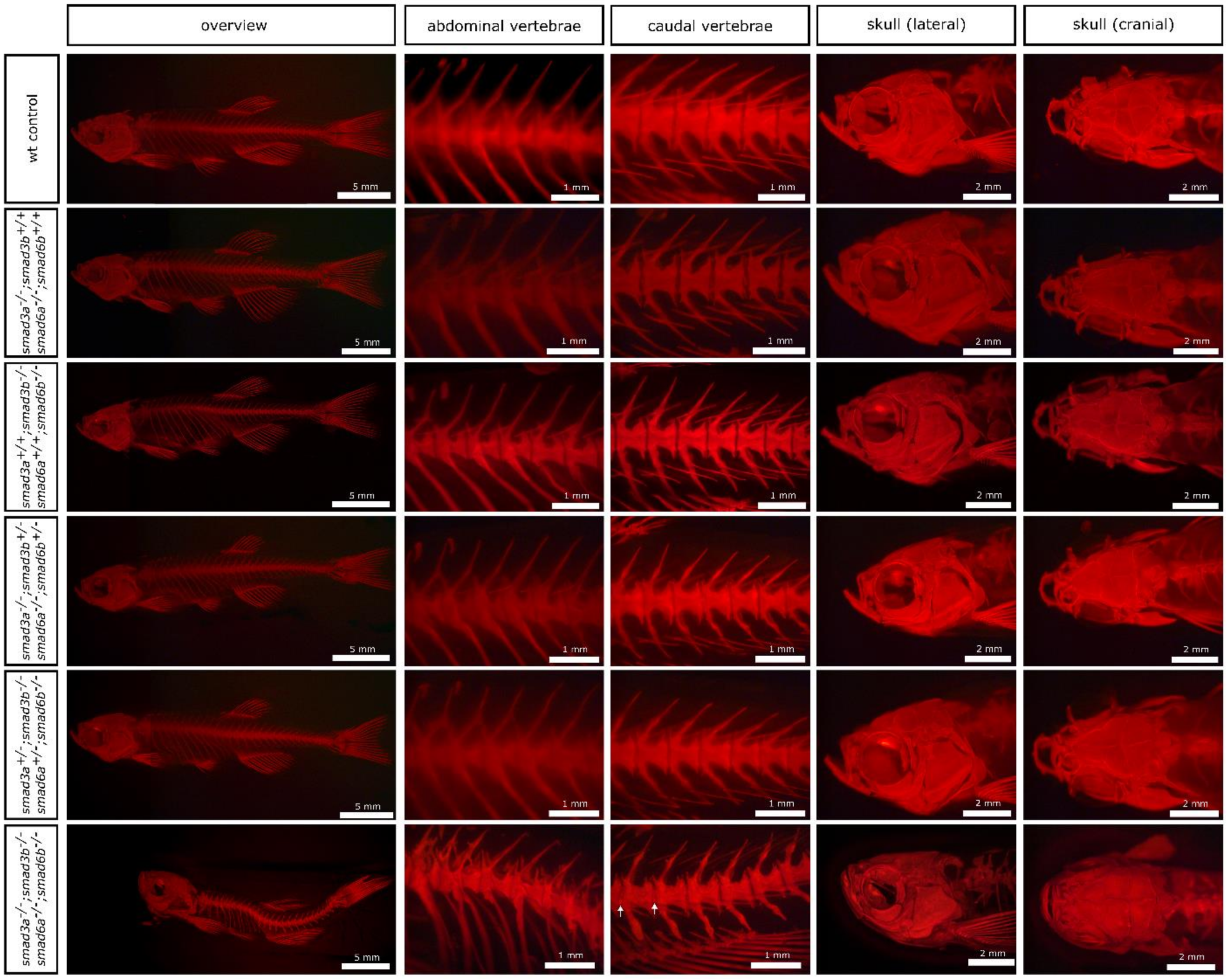


**Supplementary Figure 14: skeletal overview in 13 mpf *smad3/smad6* mutant zebrafish models.** Alizarin red staining for mineralized bone of a lateral overview picture, lateral view of abdominal and caudal vertebrae, and lateral and dorsal view of the skull, respectively of all viable *smad3/smad6* mutants. White arrowheads indicate notochord sheet mineralization. Whole-mount alizarin red staining for mineralized bone pictures taken with Leica M165 FC Fluorescent Stereo Microscope.

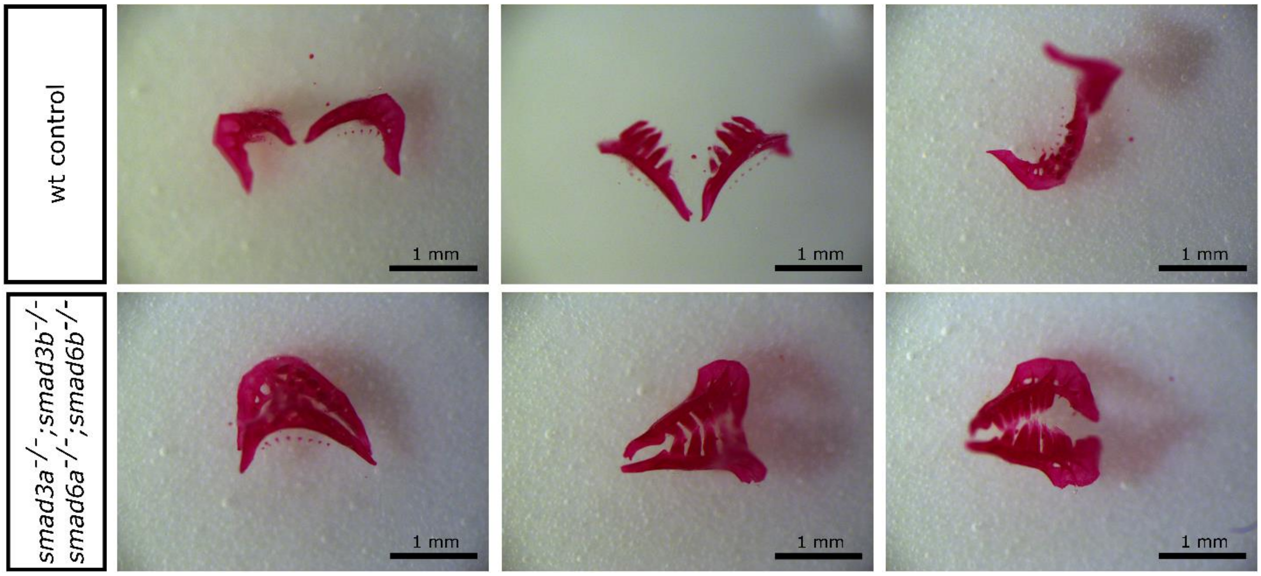


**Supplementary Figure 15: Dissected fifth ceratobranchial arch with pharyngeal teeth of 6 mpf WT and qKO zebrafish.** Alizarin red staining for mineralized bone of 3 WT and 3 qKO sets in multiple orientations. Alizarin red staining for mineralized bone pictures taken with Leica M165 FC Fluorescent Stereo Microscope.

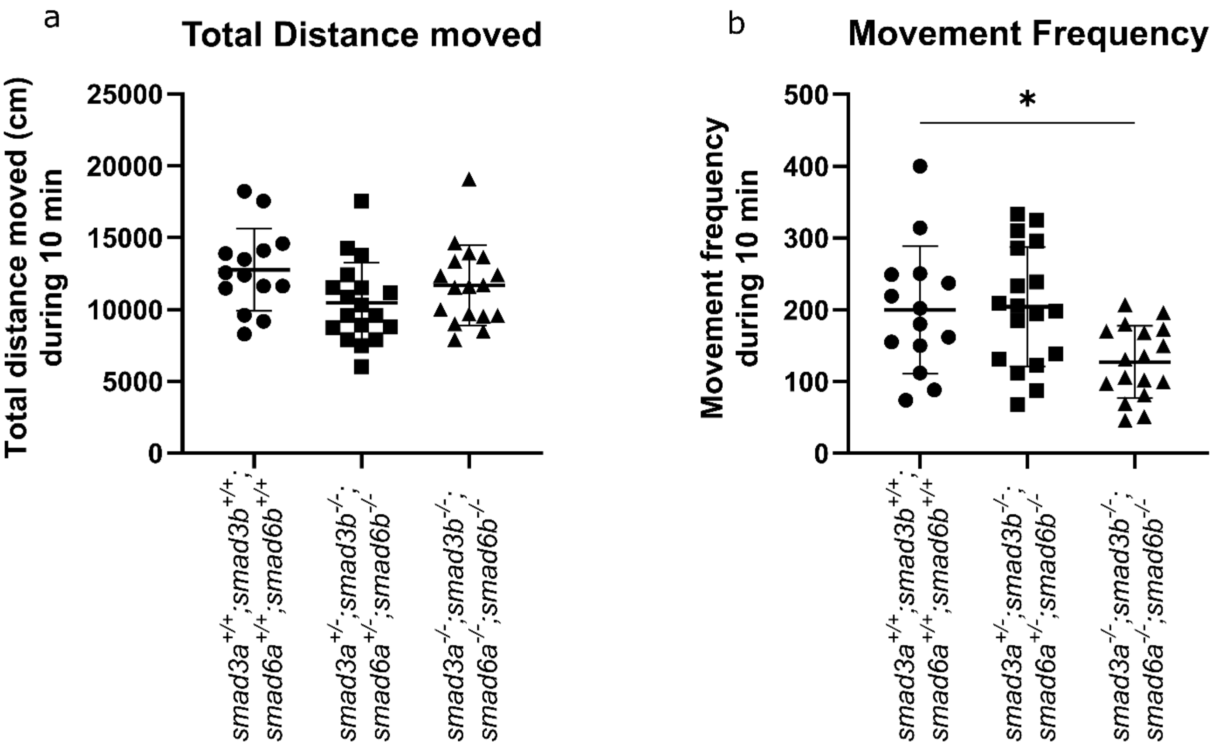


**Supplementary Figure 16: Decreased movement frequency in 9 mpf qKO zebrafish.** Swimming patterns were recorded in adult WT, *smad3a*$^{+/-}$*;smad3b*$^{-/-}$*;smad6a*$^{+/-}$*;smad6b*$^{-/-}$, and qKO zebrafish (n=14-18-17). **(a)** Total distance moved is similar between genotypes. **(b)** Significant decrease of movement frequency is observed in the qKO zebrafish. **(a, b)** One-way ANOVA with Dunnett's multiple comparison test against WT controls (*smad3a*$^{+/+}$*;smad3b*$^{+/+}$*;smad6a*$^{+/+}$*;smad6b*$^{+/+}$). *p<0.05. Data represented as average ± standard deviation.

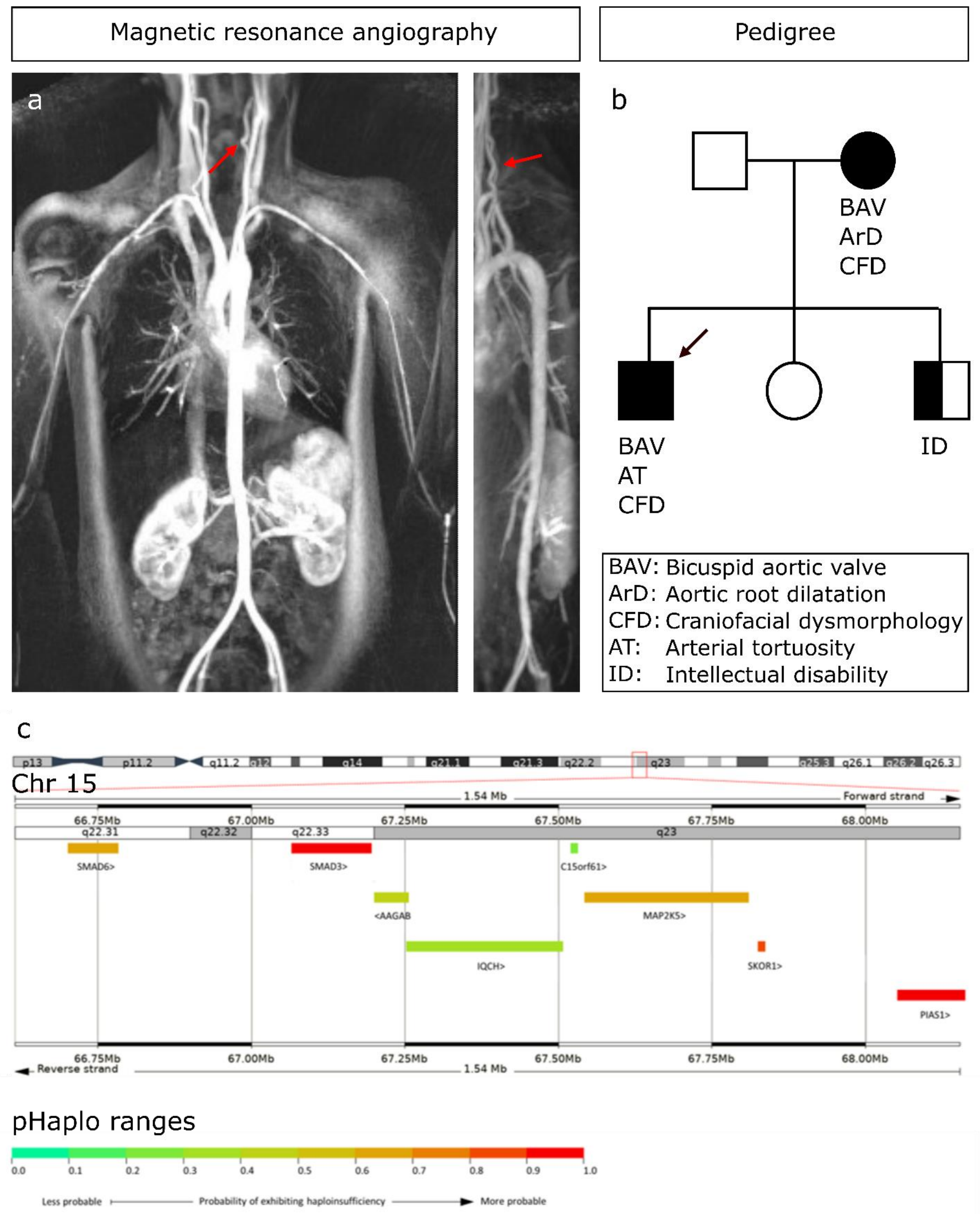


**Supplementary Figure 17: Case presentation of patient with *SMAD3* and *SMAD6* deletion. (a)** Magnetic resonance angiography shows arterial tortuosity in the vertebral arteries (red arrow). **(b)** Pedigree with index patient indicated with arrow. **(c)** 15q22.31q23 deletion present in the patient and his mother. pHaplo ranges are depicted for the deleted genes, which were calculated using DECIPHER.

### SUPPLEMENTARY TABLES

**Supplementary Table 1:** gBlock design of *smad3b, smad6a* and *smad6b.* Sequence consists of a first part matching with the random sequence – T7 promoter (red) – protospacer – tracrRNA (blue). Because of the sequence requirements of the T7 RNA Polymerase promoter, it is recommended that the first 2 nucleotides of the sgRNA transcript are GG (underlined and bold).

| Target gene | gBlock design (random sequence-T7 promoter-protospacer-tracrRNA) |
|---|---|
| smad3a | N/A |
| smad6a | ACCGCTAGCTAATACGACTCACTATA**GG**GTACAGGCGGCCCAC<br>ACGTTTTAGAGCTAGAAATAGCAAGTTAAAATAAGGCTAGTCC<br>GTTATCAACTTGAAAAAGTGGCACCGAGTCGGTGCTTTT |
| smad3b | CCGCTAGCTAATACGACTCACTATA**GG**GATGGAGTAATCATCT<br>AGGTTTTAGAGCTAGAAATAGCAAGTTAAAATAAGGCTAGTCC<br>GTTATCAACTTGAAAAAGTGGCACCGAGTCGGTGCTTTT |
| smad6b | CCGCTAGCTAATACGACTCACTATA**GG**GGGTACGGTATGTACG<br>GTGTTTTAGAGCTAGAAATAGCAAGTTAAAATAAGGCTAGTCC<br>GTTATCAACTTGAAAAAGTGGCACCGAGTCGGTGCTTTT |

**Supplementary Table 2:** Genotyping primers for *smad3a, smad3b, smad6a* and *smad6b.* All primers were used in a TD60-48 -1°C (2’ 94°C, 12 x (30’” 94°C, 30” 60*°C, 1’ 72°C), 25 x (40” 94°C, 40” 48°C, 30” 72°C), 10’ 72°C) PCR protocol. “*” each cycle, the temperature is lowered 1°C.

| **Genotyping primers** | **forward** | **reverse** |
|---|---|---|
| *smad3a* | GTGCCTCGCTGATTCTCATT | AGTTGGACGGGTCTGTGAAG |
| *smad6a* | CAACCGTGACGCCCCTAAAT | CCAAACTCCATCGGGCTCTT |
| *smad3b* | AATGTCATAATCGGTTATAATTGAGA | GGCATGTCTGGGTGAAAGTT |
| *smad6b* | TACGAGGAATCCCGCTGAGA | GAGGTTTCGACTCCTGGCTC |

**Supplementary Table 3:** Overview zebrafish lines and their respective mutations and generation. “cmg” is an in-house numbering used to identify a zebrafish line. All mutations result in the introduction of a premature termination codon.

| zebrafish lines | gene | mutation | generation |
|---|---|---|---|
| single KO | *smad3a*$^{sa2363}$ | c.682G>T | purchased (ZIRC) |
| | *smad3b*$^{cmg10}$ | c.455_459delinsATG | sgRNA injection |
| | *smad6a*$^{cmg15}$ | c.905_906delTG | sgRNA injection |
| | *smad6b*$^{cmg13}$ | c.283_287delACGGT | sgRNA injection |
| *smad3a*$^{-/-}$;*smad3b*$^{-/-}$ DKO | *smad3a*$^{sa2363}$ | c.682G>T | obtained via cross *smad3a*$^{sa2363/+}$ and *smad3b*$^{cmg10}$ $^{c.455\_459delinsATG\ /+}$ |
| | *smad3b*$^{cmg10}$ | c.455_459delinsATG | |
| *smad6a*$^{-/-}$;*smad6b*$^{-/-}$ DKO | *smad6a*$^{cmg15}$ | c.905_906delTG | obtained via cross *smad6a*$^{c.905\_906delTG\ /+}$ and *smad6b* $^{c.283\_287delACGGT\ /+}$ |
| | *smad6b*$^{cmg13}$ | c.283_287delACGGT | |
| *smad3a*$^{-/-}$;*smad6a*$^{-/-}$ DKO | *smad3a*$^{sa2363}$ | c.682G>T | sgRNA for *smad6a* injected in *smad3a*$^{sa2363/sa2363}$ |
| | *smad6a*$^{cmg59}$ | c.906_907delinsACCCATGTACCCATGT | |
| *smad3b*$^{-/-}$;*smad6b*$^{-/-}$ DKO | *smad3b*$^{cmg53}$ | c.457_458delGA | sgRNA against *smad3b* and *smad6b* co-injected. |
| | *smad6b*$^{cmg54}$ | c.274_287delACGGTATGTACGGT | |
| qKO | *smad3a*$^{sa2363}$ | c.682G>T | obtained via cross of *smad3a*$^{sa2363/+}$;*smad6a*$^{cmg59/+}$ and *smad3b*$^{cmg53/+}$;*smad6b*$^{cmg54/+}$ |
| | *smad6a*$^{cmg59}$ | c.906_907delinsACCCATGTACCCATGT | |
| | *smad3b*$^{cmg53}$ | c.457_458delGA | |
| | *smad6b*$^{cmg54}$ | c.274_287delACGGTATGTACGGT | |

**Supplementary Table 4**: List of all differentially expressed genes in the qKO compared to WT control cousins at 5 dpf sorted based on logFC. Gene ID, gene name, 10-base logarithm of fold change (logFC), 10-base logarithm counts per million (CPM), F-value, p-value and false discovery rate (FDR) are given respectively.

| gene_id | gene_name | logFC | logCPM | F | p-value | FDR |
|---|---|---|---|---|---|---|
| ENSDARG00000059039 | FO904903.1 | 10,11169874 | 0,230510502 | 313,3796196 | 1,70641E-08 | 9,76665E-06 |
| ENSDARG00000039164 | mhc1uma | 9,632267107 | -0,217608486 | 293,5927676 | 2,2941E-08 | 1,14997E-05 |
| ENSDARG00000102488 | FO704745.1 | 7,876269589 | -1,742219359 | 119,8582577 | 1,24379E-06 | 0,000135836 |
| ENSDARG00000100295 | ccl39.3 | 7,491454802 | -0,693376716 | 291,8115116 | 1,89703E-10 | 5,42884E-07 |
| ENSDARG00000100818 | si:ch73-97h19.2 | 7,028260146 | 0,855546221 | 539,1886432 | 3,59625E-12 | 2,05831E-08 |
| ENSDARG00000073821 | znf1177 | 6,564685259 | -0,93949797 | 58,00019693 | 3,7939E-06 | 0,000260052 |
| ENSDARG00000100188 | RNPS1 | 5,439639498 | 1,349172272 | 330,658362 | 8,51667E-11 | 3,24968E-07 |
| ENSDARG00000101600 | BX640576.1 | 4,574246906 | -1,853990953 | 64,09494427 | 1,88149E-06 | 0,000175101 |
| ENSDARG00000069189 | si:dkey-242h9.3 | 4,325289094 | 1,517574111 | 594,1339047 | 1,90997E-12 | 1,45756E-08 |
| ENSDARG00000097054 | CU927934.1 | 4,252798358 | 2,538815151 | 174,6395168 | 1,28839E-08 | 7,76222E-06 |
| ENSDARG00000098462 | CU570782.1 | 4,180372539 | -0,929314013 | 78,75431268 | 5,84249E-07 | 8,41245E-05 |
| ENSDARG00000035602 | dao.1 | 3,946022754 | 1,186227714 | 60,38882378 | 5,96889E-06 | 0,000334931 |
| ENSDARG00000015678 | si:dkeyp-75b4.9 | 3,813575819 | -1,60334318 | 37,46099726 | 3,28049E-05 | 0,001024748 |
| ENSDARG00000098037 | CU927934.3 | 3,667203273 | 2,240886022 | 143,7613036 | 3,04774E-08 | 1,36814E-05 |
| ENSDARG00000033760 | pmelb | 3,66239304 | 4,48567036 | 92,1706682 | 2,23377E-06 | 0,000190111 |
| ENSDARG00000059370 | nr1d4b | 3,655884365 | 4,287214313 | 30,83161834 | 0,000356381 | 0,00447558 |
| ENSDARG00000002311 | fabp11b | 3,642643189 | 2,087236528 | 54,29012393 | 1,6049E-05 | 0,000641231 |
| ENSDARG00000007480 | rpe65a | 3,629384817 | 5,260071197 | 185,3625114 | 3,70281E-08 | 1,49274E-05 |
| ENSDARG00000099428 | si:dkey-15h8.17 | 3,480678701 | 1,220894263 | 49,78767249 | 1,52397E-05 | 0,000618614 |
| ENSDARG00000095082 | ITLN1 (1 of many) | 3,177037287 | 3,370479885 | 177,4745423 | 1,04465E-08 | 6,46386E-06 |
| ENSDARG00000102401 | znf1001 | 3,170206984 | -0,908548484 | 45,70847885 | 1,17911E-05 | 0,00051223 |
| ENSDARG00000029057 | tm6sf2 | 2,981073567 | 2,545730716 | 109,3636175 | 1,67624E-07 | 3,91591E-05 |
| ENSDARG00000088524 | acot17 | 2,953511806 | 1,700540365 | 23,82965861 | 0,000606108 | 0,006343 |
| ENSDARG00000031161 | nr1d4a | 2,876018443 | 2,806928261 | 25,3368065 | 0,000553232 | 0,005994639 |
| ENSDARG00000089713 | BX004785.1 | 2,761127238 | -1,683994107 | 30,96192614 | 8,35537E-05 | 0,001801204 |
| ENSDARG00000094861 | si:dkey-163f12.11 | 2,707449788 | -2,112759454 | 24,50436671 | 0,000246499 | 0,003532483 |
| ENSDARG00000097323 | si:ch211-14i3.2 | 2,626532952 | 0,753909118 | 35,93506571 | 5,78232E-05 | 0,001443626 |
| ENSDARG00000093478 | si:ch211-197e7.3 | 2,605151536 | -1,27934524 | 29,64015124 | 0,000102817 | 0,002063014 |
| ENSDARG00000006008 | dct | 2,600034307 | 5,128677043 | 120,036933 | 2,6512E-07 | 5,41933E-05 |
| ENSDARG00000117500 | FO904870.1 | 2,59977983 | -0,963448794 | 47,70446737 | 9,40971E-06 | 0,000438749 |
| ENSDARG00000101021 | slc22a5 | 2,56351435 | 1,116339492 | 108,8007563 | 8,74327E-08 | 2,70498E-05 |
| ENSDARG00000094576 | CT025748.1 | 2,550961568 | -1,936792627 | 23,16618813 | 0,000315969 | 0,004135961 |
| ENSDARG00000104193 | CABZ01019904.1 | 2,527691561 | 0,165379291 | 29,66481555 | 0,00012768 | 0,002382318 |
| ENSDARG00000099831 | BX005396.2 | 2,483224549 | -1,191991052 | 69,68191817 | 1,17561E-06 | 0,000130022 |
| ENSDARG00000009482 | slc38a8a | 2,458303064 | -1,838869005 | 23,50555541 | 0,000296422 | 0,003966269 |
| ENSDARG00000043802 | ms4a17a.8 | 2,450918459 | 2,309752192 | 34,98869099 | 0,000101798 | 0,002046144 |
| ENSDARG00000105044 | si:ch211-51i16.3 | 2,42893437 | -1,315263071 | 17,27386564 | 0,001110776 | 0,009401148 |
| ENSDARG00000103504 | CR855311.4 | 2,415630672 | -0,438266179 | 54,80054493 | 4,46639E-06 | 0,000279381 |
| ENSDARG00000111795 | CR855311.5 | 2,392659617 | 0,378583602 | 38,31227285 | 3,26658E-05 | 0,001024748 |
| ENSDARG00000097670 | BX005048.1 | 2,381050589 | 1,49819762 | 230,6142295 | 8,47429E-10 | 1,33247E-06 |
| ENSDARG00000101818 | BX294178.1 | 2,380949766 | -0,452231631 | 47,10514449 | 1,00611E-05 | 0,00045976 |
| ENSDARG00000089749 | aqp8b | 2,367558218 | 1,159942443 | 42,36400428 | 2,41926E-05 | 0,000845598 |
| ENSDARG00000117510 | CU302321.1 | 2,362676571 | 0,962683135 | 50,43425318 | 7,85437E-06 | 0,000393475 |
| ENSDARG00000053928 | si:dkey-117a8.3 | 2,361329158 | -2,05875373 | 23,46980234 | 0,000298414 | 0,003978973 |
| ENSDARG00000042090 | si:ch73-55i23.1 | 2,355095429 | -0,232906236 | 14,63922489 | 0,00244975 | 0,01603791 |
| ENSDARG00000024771 | slc24a5 | 2,352595088 | 0,969695388 | 110,170192 | 8,11235E-08 | 2,69166E-05 |
| ENSDARG00000102364 | si:dkey-202l22.6 | 2,313722692 | 3,6415048 | 12,74155898 | 0,006004096 | 0,029314942 |
| ENSDARG00000092680 | si:dkey-58f10.12 | 2,279848273 | 0,999261385 | 57,35324326 | 3,69542E-06 | 0,000255631 |
| ENSDARG00000068947 | si:ch211-264e16.1 | 2,274324517 | 0,882464047 | 63,56064405 | 1,97157E-06 | 0,000179673 |
| ENSDARG00000079525 | slc39a5 | 2,27156517 | 1,81351407 | 42,38080037 | 2,86285E-05 | 0,000938997 |
| ENSDARG00000041490 | afp4 | 2,256998235 | 1,464457976 | 12,53175362 | 0,005287891 | 0,026786639 |
| ENSDARG00000042221 | mthfd1l | 2,239704099 | 5,097232498 | 36,54922454 | 0,000139814 | 0,002493266 |
| ENSDARG00000016818 | abcg2d | 2,238061319 | 0,514098011 | 63,06295118 | 2,05997E-06 | 0,000182089 |
| ENSDARG00000093640 | ugt5a2 | 2,235954013 | 1,051990781 | 35,68404917 | 5,64032E-05 | 0,001418599 |
| ENSDARG00000102101 | si:ch73-174h16.5 | 2,18818132 | 0,85663614 | 59,54422066 | 2,83297E-06 | 0,00022287 |
| ENSDARG00000057253 | pimr141 | 2,144677089 | -1,096884364 | 31,82343872 | 7,32365E-05 | 0,001666675 |
| ENSDARG00000102207 | zgc:86738 | 2,129269026 | 2,418366213 | 23,23524484 | 0,000610296 | 0,006368326 |
| ENSDARG00000077652 | lratb.2 | 2,124867063 | 1,008206474 | 57,49470419 | 3,43565E-06 | 0,000245431 |
| ENSDARG00000044253 | paqr3b | 2,098566189 | 2,713567613 | 59,2868118 | 5,07465E-06 | 0,000300828 |

| | | | | | | |
|---|---|---|---|---|---|---|
| ENSDARG00000010454 | guca1a | 2,092345721 | 1,293498479 | 9,945779981 | 0,010166418 | 0,042480646 |
| ENSDARG00000080188 | RF00571 | 2,089273008 | -1,52438841 | 15,59308349 | 0,001590759 | 0,011987766 |
| ENSDARG00000056151 | tyrp1b | 2,079741379 | 6,525946219 | 140,9247361 | 4,29098E-08 | 1,62231E-05 |
| ENSDARG00000039963 | fgfbp1b | 2,046639146 | 0,656922691 | 43,35622657 | 1,56367E-05 | 0,000630259 |
| ENSDARG00000075641 | steap3 | 2,028509826 | 0,287461961 | 26,97372708 | 0,000174299 | 0,002847897 |
| ENSDARG00000100164 | si:dkey-16p6.1 | 2,02545488 | 1,482669466 | 147,6985961 | 1,36761E-08 | 8,02823E-06 |
| ENSDARG00000035606 | aldh3b2 | 2,022613366 | -0,10353371 | 52,17164376 | 5,82986E-06 | 0,000330368 |
| ENSDARG00000039077 | tyr | 2,013963601 | 2,762773652 | 109,2937502 | 8,6913E-08 | 2,70498E-05 |
| ENSDARG00000028740 | msnb | 2,01101972 | 1,685682312 | 59,66388872 | 3,03618E-06 | 0,000231011 |
| ENSDARG00000058409 | hce2l1 | 2,010587406 | -0,547753156 | 11,88744118 | 0,004697621 | 0,024774786 |
| ENSDARG00000089824 | lpar4 | 2,010285454 | -0,706771354 | 16,88604969 | 0,001211574 | 0,009947658 |
| ENSDARG00000092233 | vtg1 | 1,959743191 | -1,673906463 | 12,14045154 | 0,003899819 | 0,021818784 |
| ENSDARG00000091637 | pip4k2cb | 1,95296651 | -0,50227672 | 30,79375398 | 8,57578E-05 | 0,001827163 |
| ENSDARG00000054048 | plin1 | 1,951972477 | -1,301029969 | 20,59367668 | 0,000523782 | 0,005796613 |
| ENSDARG00000103239 | si:dkey-14o6.8 | 1,943790185 | -1,227231345 | 44,39088357 | 1,37437E-05 | 0,000573131 |
| ENSDARG00000104407 | BX649453.1 | 1,928635071 | 0,288988469 | 87,12844472 | 3,2514E-07 | 6,20313E-05 |
| ENSDARG00000095858 | CU683879.1 | 1,92378479 | -1,345916991 | 17,73356784 | 0,000965907 | 0,008627969 |
| ENSDARG00000098890 | cyp2aa9 | 1,910059772 | 2,709676748 | 59,13152968 | 4,33105E-06 | 0,000275431 |
| ENSDARG00000102375 | si:ch211-204c21.1 | 1,905073724 | 0,128457631 | 34,88452321 | 4,67855E-05 | 0,001283985 |
| ENSDARG00000078052 | emilin3a | 1,883657231 | 3,799408294 | 169,4450693 | 5,85223E-09 | 4,78503E-06 |
| ENSDARG00000113320 | si:dkey-88n24.5 | 1,879250803 | -0,491102446 | 21,5385392 | 0,000433059 | 0,00509932 |
| ENSDARG00000069817 | crygm2d17 | 1,864646539 | 7,298605916 | 128,4900107 | 5,29247E-08 | 1,90485E-05 |
| ENSDARG00000093977 | si:ch211-220f16.1 | 1,844707541 | -1,481335301 | 18,66586531 | 0,000786193 | 0,00751215 |
| ENSDARG00000029204 | tyrp1a | 1,827657199 | 3,971372171 | 175,7864685 | 4,65704E-09 | 4,26473E-06 |
| ENSDARG00000086114 | CR792417.1 | 1,821374344 | -1,416202208 | 11,14937397 | 0,00524651 | 0,026646318 |
| ENSDARG00000061303 | oca2 | 1,812838951 | 1,751882656 | 106,5263679 | 9,92011E-08 | 2,80384E-05 |
| ENSDARG00000031952 | mb | 1,806528144 | 5,572201185 | 149,6674995 | 1,91402E-08 | 1,02482E-05 |
| ENSDARG00000093431 | iqcb1 | 1,799721471 | -0,253683217 | 22,8318496 | 0,000336693 | 0,004328047 |
| ENSDARG00000102496 | CR847898.4 | 1,791041203 | -1,044238744 | 12,48538875 | 0,003607928 | 0,02069136 |
| ENSDARG00000099277 | znf1039 | 1,786471535 | -0,9020384 | 27,34805668 | 0,000149746 | 0,002605552 |
| ENSDARG00000045808 | rlbp1b | 1,774933457 | 3,871764109 | 105,3021569 | 1,32032E-07 | 3,3217E-05 |
| ENSDARG00000056386 | TMC1 | 1,773498309 | 1,488489337 | 63,70632382 | 1,94653E-06 | 0,000178829 |
| ENSDARG00000001993 | myhb | 1,740384151 | 7,650650874 | 203,3353345 | 1,87372E-09 | 2,14484E-06 |
| ENSDARG00000012297 | cnga3b | 1,733555463 | 2,21244235 | 31,95555754 | 0,000104213 | 0,002082742 |
| ENSDARG00000110946 | znf1110 | 1,731349371 | -2,116047497 | 18,64715551 | 0,000789398 | 0,007530201 |
| ENSDARG00000102241 | zgc:136410 | 1,72292316 | 3,319985993 | 21,42249613 | 0,000854702 | 0,007925289 |
| ENSDARG00000045758 | cracr2ab | 1,705196133 | 1,761373601 | 57,20896026 | 3,53088E-06 | 0,000247969 |
| ENSDARG00000076554 | cdkn1a | 1,700454447 | 3,834769908 | 20,28106017 | 0,001113428 | 0,009413232 |
| ENSDARG00000104737 | si:ch211-223a21.3 | 1,686371081 | -0,149989641 | 33,86105013 | 5,41658E-05 | 0,001390214 |
| ENSDARG00000085485 | RF00568 | 1,682547267 | -1,532757851 | 23,07679981 | 0,000321362 | 0,004185021 |
| ENSDARG00000087059 | selenou1b | 1,662286855 | 0,778674428 | 46,1493218 | 1,12105E-05 | 0,00049547 |
| ENSDARG00000117305 | BX640453.1 | 1,662094316 | -1,183717009 | 17,09154311 | 0,001117359 | 0,009430845 |
| ENSDARG00000117211 | CR936462.1 | 1,639454415 | 1,78367913 | 53,905279 | 4,88472E-06 | 0,000294485 |
| ENSDARG00000115857 | znf1001 | 1,638356823 | -1,385433891 | 14,77918718 | 0,001943488 | 0,013715846 |
| ENSDARG00000092324 | CR383669.1 | 1,631922057 | -1,707596336 | 11,14141492 | 0,005181261 | 0,026448115 |
| ENSDARG00000068958 | frmd7 | 1,630437033 | -1,619673622 | 16,80274136 | 0,001194309 | 0,009867381 |
| ENSDARG00000101064 | BX511223.1 | 1,619610821 | -2,166868263 | 14,72377049 | 0,001970622 | 0,013834841 |
| ENSDARG00000100803 | zgc:173517 | 1,606127519 | 0,771824428 | 63,23879767 | 2,02823E-06 | 0,00018131 |
| ENSDARG00000089094 | si:dkey-160o24.3 | 1,603757152 | 0,690891454 | 31,40089013 | 7,81014E-05 | 0,001735973 |
| ENSDARG00000117309 | BX927081.3 | 1,599583346 | 0,562637667 | 50,00634378 | 7,32097E-06 | 0,000383538 |
| ENSDARG00000075500 | smim1 | 1,591244843 | -0,045561459 | 25,38294068 | 0,000210461 | 0,0032122 |
| ENSDARG00000039534 | rrh | 1,590664835 | 1,300941844 | 56,3462708 | 3,83736E-06 | 0,000260127 |
| ENSDARG00000103763 | l3hypdh | 1,589242352 | 1,453494205 | 113,7844957 | 6,6837E-08 | 2,25024E-05 |
| ENSDARG00000095893 | si:dkey-85n7.7 | 1,588444672 | -1,754181444 | 9,920965781 | 0,00747234 | 0,034139243 |
| ENSDARG00000011785 | tbx6 | 1,588136337 | 2,501464167 | 118,7676423 | 5,16293E-08 | 1,90485E-05 |
| ENSDARG00000076950 | kel | 1,58611483 | -0,243140321 | 14,97785537 | 0,001891614 | 0,013494906 |
| ENSDARG00000103555 | si:ch211-8c17.4 | 1,574395282 | -1,877671055 | 11,39638921 | 0,004812283 | 0,025147774 |
| ENSDARG00000097838 | si:dkey-42l23.1 | 1,565843513 | -2,042152198 | 8,420609737 | 0,01209803 | 0,047753843 |
| ENSDARG00000099687 | CT009597.1 | 1,542999368 | -2,0573889 | 10,07244735 | 0,007131913 | 0,033058923 |
| ENSDARG00000112862 | CU182107.2 | 1,542819243 | -1,800318493 | 12,69919734 | 0,003342032 | 0,019633689 |
| ENSDARG00000041381 | arntl2 | 1,535096639 | 4,226319123 | 19,20229544 | 0,001341001 | 0,010667432 |
| ENSDARG00000090687 | rcbtb2 | 1,512853026 | 3,335261205 | 99,28818208 | 1,5077E-07 | 3,71153E-05 |
| ENSDARG00000079519 | eys | 1,504833376 | -0,708433445 | 21,86184953 | 0,000406279 | 0,004910958 |
| ENSDARG00000095527 | CR392363.2 | 1,499877616 | -1,72838744 | 10,75825623 | 0,0057992 | 0,028632067 |
| ENSDARG00000096603 | bmb | 1,490907443 | 2,293401339 | 34,33842728 | 6,09847E-05 | 0,001496445 |
| ENSDARG00000077862 | si:dkey-169i5.4 | 1,489121898 | 1,973733811 | 30,17557224 | 0,000110905 | 0,002173188 |
| ENSDARG00000001975 | hsd11b2 | 1,488157868 | 4,741457258 | 191,014788 | 2,77267E-09 | 2,75989E-06 |
| ENSDARG00000106564 | CT027676.1 | 1,479418463 | -0,441154732 | 17,79650405 | 0,00095238 | 0,008533112 |
| ENSDARG00000039099 | aep1 | 1,47459467 | 6,225360593 | 101,2100234 | 1,85637E-07 | 4,20788E-05 |
| ENSDARG00000095025 | CR388166.2 | 1,460519251 | -0,037549889 | 26,10570016 | 0,000185316 | 0,002945598 |
| ENSDARG00000053066 | tubb1 | 1,454393422 | 1,652459377 | 19,41256556 | 0,000808285 | 0,007634525 |

| | | | | | | |
|---|---|---|---|---|---|---|
| ENSDARG00000042811 | fgf1b | 1,450038138 | 0,941454863 | 46,421893 | 1,0868E-05 | 0,000484071 |
| ENSDARG00000079403 | si:dkey-204l11.1 | 1,441466669 | 0,242844002 | 11,92188307 | 0,004435564 | 0,023879699 |
| ENSDARG00000101200 | zgc:112964 | 1,434076643 | 2,57231763 | 22,54902676 | 0,000490055 | 0,005555483 |
| ENSDARG00000114317 | BX927253.3 | 1,431186108 | -0,245066686 | 15,20287733 | 0,001749715 | 0,012802162 |
| ENSDARG00000042370 | ptafr | 1,430039941 | -1,007068572 | 12,00447013 | 0,00404868 | 0,022410655 |
| ENSDARG00000107748 | RF00001 | 1,428650751 | -1,831007564 | 10,09669862 | 0,0070791 | 0,032874021 |
| ENSDARG00000093019 | si:dkey-83k24.5 | 1,427617087 | 2,262711821 | 33,08978443 | 7,03846E-05 | 0,001619673 |
| ENSDARG00000103398 | fabp1b.2 | 1,426550209 | -0,596548789 | 17,65854708 | 0,000982323 | 0,00868313 |
| ENSDARG00000098721 | BX649331.1 | 1,424709004 | -0,085834386 | 30,4102112 | 9,10396E-05 | 0,001900641 |
| ENSDARG00000105208 | zgc:85843 | 1,42151018 | 3,171705212 | 28,02447902 | 0,000200556 | 0,003115018 |
| ENSDARG00000029930 | SPDEF | 1,410536551 | 0,08008742 | 34,88703042 | 4,67689E-05 | 0,001283985 |
| ENSDARG00000014358 | optc | 1,404802754 | 0,23782291 | 36,95554727 | 3,51188E-05 | 0,001067743 |
| ENSDARG00000038643 | alas2 | 1,403173605 | 5,024123277 | 15,22049163 | 0,003101687 | 0,018735637 |
| ENSDARG00000091298 | pmela | 1,400806963 | 6,204386236 | 135,3061661 | 2,33969E-08 | 1,14997E-05 |
| ENSDARG00000102580 | CU571310.1 | 1,396041244 | 0,218205067 | 23,52515593 | 0,000295336 | 0,003963279 |
| ENSDARG00000001913 | palmda | 1,388797459 | 4,794255795 | 215,8979869 | 1,28498E-09 | 1,73049E-06 |
| ENSDARG00000103515 | vcana | 1,383993283 | 0,358160774 | 27,95616805 | 0,000135244 | 0,002456626 |
| ENSDARG00000017400 | klf1 | 1,377930772 | 1,646460176 | 17,3366108 | 0,001256949 | 0,010208088 |
| ENSDARG00000002956 | si:dkey-32n7.4 | 1,376590445 | 2,315978028 | 62,00633598 | 2,2632E-06 | 0,000191902 |
| ENSDARG00000099121 | si:dkey-6d5.1 | 1,371900221 | -1,179835499 | 28,20092546 | 0,000129869 | 0,00240746 |
| ENSDARG00000100184 | BX908782.1 | 1,365567937 | 0,626299488 | 21,27681034 | 0,000456236 | 0,005283288 |
| ENSDARG00000117535 | AL929493.2 | 1,365367718 | 1,16976546 | 57,17677695 | 3,5418E-06 | 0,000247969 |
| ENSDARG00000044550 | hif1al2 | 1,360484524 | 2,131134874 | 19,48072357 | 0,000830104 | 0,007760069 |
| ENSDARG00000098245 | cidea | 1,359493275 | -0,63985712 | 21,80972688 | 0,000410465 | 0,004927926 |
| ENSDARG00000012395 | mmp13a | 1,354440048 | 1,224486375 | 13,89225122 | 0,002806135 | 0,017548114 |
| ENSDARG00000105439 | si:dkey-273g18.1 | 1,352554039 | -1,510161662 | 10,03862215 | 0,007206341 | 0,033302779 |
| ENSDARG00000105092 | FP236157.3 | 1,352176115 | -1,959882047 | 8,27350509 | 0,012709931 | 0,049460904 |
| ENSDARG00000103347 | cyp2aa3 | 1,349229514 | 2,318692999 | 53,28717136 | 5,19989E-06 | 0,000304466 |
| ENSDARG00000092272 | BX511161.1 | 1,349200244 | 0,532246076 | 19,95327605 | 0,00059777 | 0,006292298 |
| ENSDARG00000097634 | CU306817.1 | 1,34756501 | -1,282106829 | 12,93140143 | 0,003138468 | 0,018874634 |
| ENSDARG00000100105 | impg2b | 1,345483668 | 2,049575102 | 40,04599003 | 2,33959E-05 | 0,000831717 |
| ENSDARG00000101164 | nansb | 1,343293577 | 1,39479642 | 38,0610709 | 3,02822E-05 | 0,000981985 |
| ENSDARG00000016718 | mmp11b | 1,340101035 | 3,373197769 | 87,58268537 | 3,15417E-07 | 6,11962E-05 |
| ENSDARG00000044524 | def6b | 1,339852351 | 1,076774437 | 68,96464974 | 1,2465E-06 | 0,000135836 |
| ENSDARG00000052958 | nppb | 1,331163234 | -0,013616957 | 8,485377216 | 0,012513428 | 0,048904476 |
| ENSDARG00000102999 | si:dkey-246j6.1 | 1,327380107 | -0,493668257 | 32,32368915 | 6,79215E-05 | 0,001584604 |
| ENSDARG00000117113 | zgc:173607 | 1,317893771 | -1,563679479 | 12,14015761 | 0,003898027 | 0,021818784 |
| ENSDARG00000060962 | btr04 | 1,317479253 | 1,596914019 | 32,54089066 | 6,57531E-05 | 0,001555115 |
| ENSDARG00000091043 | pcdh2ab6 | 1,31589458 | 1,278993641 | 34,98452674 | 4,61286E-05 | 0,001275445 |
| ENSDARG00000095464 | gstt2 | 1,314898391 | 4,124537609 | 81,81399362 | 4,77163E-07 | 7,85912E-05 |
| ENSDARG00000007377 | odc1 | 1,307770327 | 5,774901615 | 81,75372894 | 6,29171E-07 | 8,82877E-05 |
| ENSDARG00000109267 | CABZ01024501.1 | 1,30413885 | 1,197224897 | 35,27674367 | 4,42692E-05 | 0,001252779 |
| ENSDARG00000101322 | tfr1a | 1,301644175 | 4,206266204 | 36,11570345 | 6,35423E-05 | 0,001523284 |
| ENSDARG00000102884 | si:ch211-145h19.5 | 1,298122508 | -1,043512815 | 14,65331082 | 0,002005756 | 0,013979662 |
| ENSDARG00000061480 | si:ch211-132b12.1 | 1,29564918 | 1,038434084 | 21,69575444 | 0,000425116 | 0,005034972 |
| ENSDARG00000102480 | BX005417.2 | 1,293360728 | -0,474675345 | 18,16120658 | 0,000878161 | 0,008054736 |
| ENSDARG00000023587 | kcnk5a | 1,289900219 | 2,915091015 | 11,36602508 | 0,006602885 | 0,031375352 |
| ENSDARG00000103950 | pcdh1gc6 | 1,288670882 | 4,307739788 | 36,17609 | 6,33569E-05 | 0,001520508 |
| ENSDARG00000105464 | si:ch211-285c6.6 | 1,278278097 | -1,457332088 | 14,36175693 | 0,002159045 | 0,014680477 |
| ENSDARG00000055160 | chadla | 1,27573067 | 1,297062679 | 47,55798875 | 9,56432E-06 | 0,000444149 |
| ENSDARG00000094708 | sftpba | 1,27443153 | 2,121693555 | 48,01918764 | 9,08713E-06 | 0,000430726 |
| ENSDARG00000100288 | impg2b | 1,270843972 | 1,520107267 | 36,00159089 | 4,00175E-05 | 0,001168849 |
| ENSDARG00000034940 | slc1a7a | 1,267860426 | 2,545413125 | 25,15298153 | 0,000265697 | 0,003693296 |
| ENSDARG00000044767 | tspan10 | 1,264587306 | 1,102605655 | 35,14346622 | 4,51063E-05 | 0,00126552 |
| ENSDARG00000104654 | leap2 | 1,264166042 | 4,78866535 | 47,86569288 | 1,50907E-05 | 0,000613651 |
| ENSDARG00000114704 | FP016205.1 | 1,263153303 | 2,465120998 | 42,88731351 | 1,64428E-05 | 0,000653545 |
| ENSDARG00000037932 | lin7b | 1,257030203 | 4,684085028 | 46,1763368 | 1,78991E-05 | 0,000687735 |
| ENSDARG00000092664 | si:dkey-93m18.3 | 1,255553054 | -1,967182283 | 14,00025818 | 0,002368283 | 0,015683965 |
| ENSDARG00000003732 | mitfa | 1,254467215 | 1,270486462 | 40,32837409 | 2,25708E-05 | 0,000808665 |
| ENSDARG00000060246 | slc16a6b | 1,254246326 | 4,518236313 | 15,54992519 | 0,002660567 | 0,01691973 |
| ENSDARG00000102430 | opn6a | 1,247355788 | 3,970890374 | 47,83651357 | 1,17831E-05 | 0,00051223 |
| ENSDARG00000089974 | SLC15A5 | 1,245093726 | -0,683870882 | 15,87665543 | 0,001485672 | 0,011383193 |
| ENSDARG00000113834 | BX537305.1 | 1,244682605 | -0,059701615 | 19,51330833 | 0,000655603 | 0,006691555 |
| ENSDARG00000086838 | itga2.3 | 1,24106563 | 1,015649913 | 31,97030158 | 7,1628E-05 | 0,001638213 |
| ENSDARG00000068407 | six9 | 1,240985927 | -0,191371015 | 21,57407415 | 0,000430018 | 0,005077278 |
| ENSDARG00000068114 | f11r.2 | 1,240790375 | 2,356383521 | 29,35700617 | 0,0001167 | 0,002247039 |
| ENSDARG00000079412 | ftr02 | 1,240661281 | 1,275145526 | 53,34147254 | 5,17129E-06 | 0,000303568 |
| ENSDARG00000096810 | BX279523.1 | 1,240275157 | 0,441533525 | 62,61784887 | 2,14291E-06 | 0,000185316 |
| ENSDARG00000096186 | si:dkey-247i3.5 | 1,237048077 | 0,06949515 | 32,98898039 | 6,15265E-05 | 0,001499917 |
| ENSDARG00000014395 | selenoo2 | 1,236436006 | 0,317409576 | 25,49161197 | 0,000206441 | 0,003171981 |
| ENSDARG00000080775 | RF00571 | 1,236028153 | -1,728281137 | 10,33210363 | 0,006589345 | 0,03133547 |

| | | | | | | |
|---|---|---|---|---|---|---|
| ENSDARG00000097973 | si:ch1073-190k2.1 | 1,232854902 | 2,034445789 | 18,99994329 | 0,000853868 | 0,007923974 |
| ENSDARG00000097929 | BX005392.3 | 1,228570235 | -0,717440755 | 8,932205061 | 0,010221369 | 0,042620877 |
| ENSDARG00000055163 | epoa | 1,222125337 | -1,015931861 | 12,86930186 | 0,003191468 | 0,019077149 |
| ENSDARG00000040427 | acot19 | 1,215605702 | 1,639462902 | 11,0885431 | 0,006080339 | 0,029578145 |
| ENSDARG00000043798 | ms4a17a.1 | 1,215225355 | -0,895040392 | 14,61817789 | 0,002023545 | 0,014055783 |
| ENSDARG00000086998 | zgc:64002 | 1,209434893 | 0,971320344 | 23,04251136 | 0,000323459 | 0,004205151 |
| ENSDARG00000099247 | si:dkey-68o6.5 | 1,208827635 | -1,714224355 | 9,626567627 | 0,008189393 | 0,036426648 |
| ENSDARG00000101393 | si:dkey-31g6.6 | 1,207310866 | 5,331969871 | 16,156343 | 0,002399747 | 0,015810019 |
| ENSDARG00000115701 | crygm2d9 | 1,202775206 | 6,623560238 | 90,70928604 | 2,63053E-07 | 5,41933E-05 |
| ENSDARG00000101481 | rbp5 | 1,197239864 | 4,688894362 | 86,87250964 | 3,3077E-07 | 6,24314E-05 |
| ENSDARG00000053345 | si:dkey-56i24.1 | 1,192166524 | -0,113685743 | 12,01196362 | 0,004040186 | 0,02237446 |
| ENSDARG00000112831 | znf1116 | 1,190534656 | -2,047663395 | 8,458685316 | 0,01194528 | 0,047363222 |
| ENSDARG00000077372 | tfr1b | 1,187445117 | 5,953585471 | 12,22386418 | 0,005934092 | 0,029084801 |
| ENSDARG00000105570 | crestin | 1,186862578 | 0,211299499 | 23,67445817 | 0,000287215 | 0,003876829 |
| ENSDARG00000063079 | ago3b | 1,185574903 | 2,53732232 | 27,62589102 | 0,000157205 | 0,002683848 |
| ENSDARG00000089124 | hbae1.3 | 1,181766363 | 4,58477101 | 19,08963372 | 0,001189127 | 0,009835216 |
| ENSDARG00000100112 | znf1118 | 1,179233951 | 1,24393951 | 49,26576487 | 7,92851E-06 | 0,00039562 |
| ENSDARG00000043035 | capn3b | 1,176191864 | 4,47315286 | 53,7194902 | 6,14901E-06 | 0,00033941 |
| ENSDARG00000100723 | znf1097 | 1,17605054 | -0,955956891 | 10,12092006 | 0,007026805 | 0,032717443 |
| ENSDARG00000101790 | si:ch73-299h12.3 | 1,172044702 | 1,391728864 | 48,09863141 | 9,00773E-06 | 0,000428738 |
| ENSDARG00000084681 | RF00001 | 1,165574348 | -1,844873168 | 12,29838851 | 0,00373047 | 0,021176636 |
| ENSDARG00000000551 | slc1a4 | 1,164529579 | 5,075434829 | 41,74480358 | 3,06856E-05 | 0,000992836 |
| ENSDARG00000052700 | SLC9B2 | 1,160416536 | 4,373354267 | 28,2969325 | 0,000194544 | 0,003042275 |
| ENSDARG00000059792 | trpm5 | 1,157423369 | 2,468836471 | 41,01310457 | 2,07056E-05 | 0,000758453 |
| ENSDARG00000101385 | f8 | 1,154294777 | 3,628479176 | 23,56393924 | 0,000408121 | 0,004919583 |
| ENSDARG00000078618 | inpp5kb | 1,149692331 | 2,265731664 | 36,27948275 | 3,85136E-05 | 0,001139188 |
| ENSDARG00000035555 | gbgt1l3 | 1,144755708 | 1,65951742 | 22,31163091 | 0,00037591 | 0,004629419 |
| ENSDARG00000040938 | ssuh2.1 | 1,142894772 | -1,412086533 | 11,68147812 | 0,004435227 | 0,023879699 |
| ENSDARG00000071450 | tdrd5 | 1,142055527 | -1,027311448 | 9,460779881 | 0,008628255 | 0,037900091 |
| ENSDARG00000008306 | rdh5 | 1,140481352 | 2,949243667 | 45,1932063 | 1,2514E-05 | 0,000534508 |
| ENSDARG00000005176 | zgc:101040 | 1,139977817 | 3,123266642 | 44,85163209 | 1,30702E-05 | 0,000551057 |
| ENSDARG00000033539 | paics | 1,138387068 | 7,287431652 | 18,60453506 | 0,001303232 | 0,010468841 |
| ENSDARG00000012881 | slc4a1a | 1,136391334 | 5,888092738 | 13,10700327 | 0,00470422 | 0,024792451 |
| ENSDARG00000097816 | si:dkey-117a8.1 | 1,135997377 | -1,236842821 | 9,354272045 | 0,008924609 | 0,03885888 |
| ENSDARG00000037961 | rcn3 | 1,134475225 | 5,086088564 | 54,43496039 | 6,23499E-06 | 0,000341673 |
| ENSDARG00000089297 | si:dkeyp-115d7.2 | 1,133774767 | -0,919386101 | 12,81990138 | 0,00323437 | 0,019250309 |
| ENSDARG00000093862 | si:ch211-274k16.2 | 1,129020704 | -0,21419943 | 20,25312697 | 0,000561722 | 0,006057491 |
| ENSDARG00000051874 | stra6 | 1,121849035 | 4,652486781 | 61,77239479 | 2,57496E-06 | 0,000208308 |
| ENSDARG00000019651 | zgc:162239 | 1,11696198 | -1,381119421 | 13,63534579 | 0,002603637 | 0,016696826 |
| ENSDARG00000068288 | lamc2 | 1,115883267 | 1,498759072 | 61,34595597 | 2,40191E-06 | 0,000199237 |
| ENSDARG00000010047 | neu3.2 | 1,115712286 | 1,62081535 | 21,11377832 | 0,000474829 | 0,005451724 |
| ENSDARG00000104105 | ttf2 | 1,110561559 | 3,747558991 | 106,9083191 | 9,71032E-08 | 2,80384E-05 |
| ENSDARG00000053062 | CR628323.2 | 1,106484175 | -2,003536466 | 10,53311992 | 0,006202036 | 0,030006215 |
| ENSDARG00000097815 | CU928129.1 | 1,101206836 | 0,976504677 | 42,57859528 | 1,70698E-05 | 0,000665752 |
| ENSDARG00000042014 | cyp11c1 | 1,100725364 | 2,803694746 | 67,31631014 | 1,42885E-06 | 0,000145321 |
| ENSDARG00000100793 | CR450785.1 | 1,098079598 | 1,937151346 | 45,14499089 | 1,25842E-05 | 0,000536506 |
| ENSDARG00000018459 | msrb2 | 1,087252734 | 5,55554399 | 49,67864043 | 1,04176E-05 | 0,000469489 |
| ENSDARG00000091511 | gpx7 | 1,087236952 | 2,644142028 | 48,91293678 | 8,23824E-06 | 0,000406479 |
| ENSDARG00000103417 | si:dkey-269o24.6 | 1,086518069 | 0,170919389 | 20,97002012 | 0,000485245 | 0,005524291 |
| ENSDARG00000016733 | psat1 | 1,086423082 | 5,444211429 | 46,59079045 | 1,53263E-05 | 0,000621029 |
| ENSDARG00000077559 | LO017815.1 | 1,085829006 | 1,559743306 | 27,92103011 | 0,000136037 | 0,002460837 |
| ENSDARG00000005626 | RHOU (1 of many) | 1,080999463 | 1,490410839 | 35,22667023 | 4,45816E-05 | 0,001256775 |
| ENSDARG00000043918 | FO904943.1 | 1,080024274 | -0,396123588 | 12,26561433 | 0,00376448 | 0,021290516 |
| ENSDARG00000089139 | im:7145024 | 1,079080139 | 1,68991564 | 58,37010918 | 3,16178E-06 | 0,000232441 |
| ENSDARG00000098604 | si:dkey-14o6.4 | 1,075077192 | -1,729223567 | 9,923401208 | 0,007466719 | 0,034132946 |
| ENSDARG00000101240 | CU651662.1 | 1,073111235 | 0,872724187 | 19,1210626 | 0,00071266 | 0,007064387 |
| ENSDARG00000102594 | zgc:194627 | 1,070260265 | 2,706393683 | 35,31786962 | 4,40144E-05 | 0,001251091 |
| ENSDARG00000102845 | dclk2b | 1,06754128 | 2,30816415 | 28,00271722 | 0,000134202 | 0,002448149 |
| ENSDARG00000077387 | tcte1 | 1,064774245 | -0,676954194 | 10,44443832 | 0,006369555 | 0,030616125 |
| ENSDARG00000112541 | znf1093 | 1,059729007 | 0,356071678 | 28,95873789 | 0,000114722 | 0,002218276 |
| ENSDARG00000103914 | THEM6 | 1,058899411 | 2,184578816 | 41,9344036 | 1,84689E-05 | 0,000697734 |
| ENSDARG00000058839 | susd1 | 1,055724667 | 0,562794011 | 27,89891118 | 0,000136538 | 0,002467171 |
| ENSDARG00000102959 | zgc:171220 | 1,055039787 | 0,941670966 | 32,34995737 | 6,76549E-05 | 0,001582116 |
| ENSDARG00000105016 | RF00213 | 1,054325125 | -1,343610751 | 8,43413468 | 0,012043512 | 0,047653675 |
| ENSDARG00000093753 | BX004774.2 | 1,054305569 | 3,355663193 | 20,9930551 | 0,000609211 | 0,006359908 |
| ENSDARG00000101508 | nr6a1a | 1,053691423 | 0,159012951 | 12,61150305 | 0,003422854 | 0,019914315 |
| ENSDARG00000117132 | BX663613.1 | 1,053037405 | -0,260663883 | 15,35876877 | 0,001684105 | 0,012473598 |
| ENSDARG00000111008 | CU694317.1 | 1,052085698 | -1,993109911 | 8,547976912 | 0,011595784 | 0,046452123 |
| ENSDARG00000023722 | hcrtr2 | 1,05070975 | -0,953064636 | 11,29835965 | 0,004950421 | 0,025618207 |
| ENSDARG00000032242 | tnnt2c | 1,050444651 | 3,476229935 | 55,13412576 | 4,32117E-06 | 0,000275431 |
| ENSDARG00000012945 | tnfsf13b | 1,049456335 | -0,109241411 | 14,64323082 | 0,002010841 | 0,014005538 |

| | | | | | | |
|---|---|---|---|---|---|---|
| ENSDARG00000098237 | fbn2b | 1,04885657 | 6,772470389 | 34,76549818 | 6,94331E-05 | 0,001607283 |
| ENSDARG00000099252 | ipcef1 | 1,048605248 | 1,766819672 | 35,75927285 | 4,13843E-05 | 0,001194769 |
| ENSDARG00000044326 | BX950188.1 | 1,047752241 | 0,853499396 | 10,53543693 | 0,006366612 | 0,030616125 |
| ENSDARG00000020028 | cps1 | 1,047259088 | 2,853742786 | 28,55680175 | 0,000129071 | 0,002398493 |
| ENSDARG00000095150 | si:dkey-114c15.5 | 1,04613074 | -0,376746524 | 19,5722842 | 0,000647488 | 0,006623586 |
| ENSDARG00000112473 | BX572103.1 | 1,045142378 | -0,473638764 | 12,17410319 | 0,003861359 | 0,021699056 |
| ENSDARG00000030270 | tnnt3a | 1,04473018 | 8,156552224 | 105,342566 | 1,06042E-07 | 2,89015E-05 |
| ENSDARG00000102554 | zgc:112433 | 1,043547664 | 1,698257627 | 40,90365998 | 2,09914E-05 | 0,000766472 |
| ENSDARG00000087736 | si:dkey-118k5.3 | 1,04286865 | 0,740459406 | 15,85447018 | 0,001493598 | 0,011421076 |
| ENSDARG00000104015 | fgfr1bl | 1,042615752 | 1,833513015 | 15,37241786 | 0,001792887 | 0,012985242 |
| ENSDARG00000094362 | CR762497.1 | 1,042367543 | 0,802066138 | 9,883223487 | 0,007802888 | 0,035269362 |
| ENSDARG00000038785 | abcf2a | 1,039055101 | 5,877678798 | 58,97882684 | 3,37749E-06 | 0,000243196 |
| ENSDARG00000000423 | si:ch73-314g15.3 | 1,033136826 | 2,139361043 | 77,63844535 | 6,34266E-07 | 8,82877E-05 |
| ENSDARG00000035151 | drll.1 | 1,029518033 | 0,263009184 | 11,53104615 | 0,004629743 | 0,024546861 |
| ENSDARG00000068941 | zgc:113983 | 1,028886375 | -0,737323018 | 17,48758463 | 0,00102095 | 0,008905112 |
| ENSDARG00000109251 | si:dkey-4e4.1 | 1,02675053 | -0,620778086 | 14,86405114 | 0,001902767 | 0,013545381 |
| ENSDARG00000071212 | p3h1 | 1,025431467 | 4,648136427 | 78,8398134 | 5,80608E-07 | 8,41245E-05 |
| ENSDARG00000059053 | slc13a4 | 1,021384956 | 4,44585764 | 50,23816036 | 7,6654E-06 | 0,000389117 |
| ENSDARG00000088316 | si:ch73-302o18.2 | 1,020419542 | -1,615564463 | 9,351581157 | 0,008932248 | 0,038881951 |
| ENSDARG00000103031 | znf1029 | 1,018550624 | -0,256561343 | 12,16104352 | 0,003875419 | 0,021746038 |
| ENSDARG00000061265 | lmln | 1,017003357 | 2,23885541 | 22,37368326 | 0,000376628 | 0,004634243 |
| ENSDARG00000003701 | cldng | 1,015815467 | -1,294747796 | 8,690616637 | 0,011061854 | 0,044938845 |
| ENSDARG00000059751 | abtb2a | 1,015127517 | 2,874245857 | 47,35054295 | 9,78832E-06 | 0,000450893 |
| ENSDARG00000053520 | LO017829.1 | 1,014927975 | 2,448073415 | 35,03227946 | 4,58187E-05 | 0,001271483 |
| ENSDARG00000078331 | best1 | 1,01451241 | 0,303308981 | 17,41039673 | 0,001038962 | 0,008996212 |
| ENSDARG00000070480 | agr2 | 1,013695584 | 5,938824208 | 32,35874122 | 0,000100706 | 0,002029538 |
| ENSDARG00000034572 | gpr143 | 1,007325946 | 2,411376986 | 37,79197068 | 3,13849E-05 | 0,00100304 |
| ENSDARG00000105718 | CABZ01068273.1 | 1,002504985 | -1,049808829 | 8,662604494 | 0,011164413 | 0,045206594 |
| ENSDARG00000071905 | CR847851.1 | 1,001340412 | -1,454257077 | 8,472567025 | 0,011890149 | 0,04722312 |
| ENSDARG00000099538 | si:ch211-188p14.3 | -1,000006697 | -0,03612783 | 10,40108344 | 0,006453359 | 0,030908619 |
| ENSDARG00000007024 | uox | -1,00131504 | 5,83923402 | 34,77414846 | 6,86638E-05 | 0,001595031 |
| ENSDARG00000097019 | dhrs13b.1 | -1,001618576 | 0,085766701 | 25,62121805 | 0,000201761 | 0,003125242 |
| ENSDARG00000055534 | grk7b | -1,001943231 | 2,22700528 | 29,93019657 | 9,81863E-05 | 0,002003456 |
| ENSDARG00000062947 | amn | -1,002103361 | 2,18123716 | 21,54997895 | 0,000442183 | 0,005175526 |
| ENSDARG00000001818 | c3b.2 | -1,003106753 | 4,911640436 | 50,96674986 | 7,38379E-06 | 0,000385289 |
| ENSDARG00000086458 | hdac10 | -1,003416741 | 3,556127917 | 81,74743737 | 4,71009E-07 | 7,83027E-05 |
| ENSDARG00000089434 | etv7 | -1,003832312 | 0,155987227 | 18,91244592 | 0,000745328 | 0,007292053 |
| ENSDARG00000092920 | si:ch211-106h4.12 | -1,004010366 | 4,146256965 | 68,40619308 | 1,3051E-06 | 0,000137098 |
| ENSDARG00000105145 | si:ch211-282j17.12 | -1,005195699 | -1,027418336 | 12,22600279 | 0,003806066 | 0,021467373 |
| ENSDARG00000042214 | tmem243a | -1,005420415 | 1,175674687 | 24,81574638 | 0,000232969 | 0,003388556 |
| ENSDARG00000079892 | si:dkey-183c2.4 | -1,005782793 | 0,48215097 | 18,35357623 | 0,000841712 | 0,007839774 |
| ENSDARG00000105511 | BX248521.2 | -1,006115205 | 0,359090495 | 19,11815237 | 0,000713104 | 0,007064387 |
| ENSDARG00000099149 | hs3st3b1a | -1,009131547 | 2,505766688 | 46,63146075 | 1,06129E-05 | 0,000474577 |
| ENSDARG00000105274 | CABZ01074130.1 | -1,009152892 | 7,324879484 | 31,51113622 | 0,000105761 | 0,002098172 |
| ENSDARG00000000212 | krt97 | -1,009675191 | 8,124398669 | 21,96128232 | 0,000548485 | 0,005959669 |
| ENSDARG00000008866 | cabp4 | -1,010702353 | 0,991179978 | 11,73328712 | 0,004468133 | 0,023956308 |
| ENSDARG00000091584 | si:ch73-217n20.1 | -1,010774065 | 3,910782714 | 66,45669876 | 1,53602E-06 | 0,000151577 |
| ENSDARG00000044982 | dhrs3a | -1,011730194 | 3,898565715 | 46,63442142 | 1,07097E-05 | 0,000477949 |
| ENSDARG00000002758 | dedd1 | -1,011807973 | 6,609253055 | 35,60635784 | 5,89543E-05 | 0,001460713 |
| ENSDARG00000079903 | si:ch73-343l4.2 | -1,01221719 | -0,897921929 | 15,65662403 | 0,001566485 | 0,011849349 |
| ENSDARG00000002298 | ankrd22 | -1,012919646 | 1,186358927 | 25,50010493 | 0,00020613 | 0,003169337 |
| ENSDARG00000022951 | pth2 | -1,012920421 | 0,545451453 | 24,75325633 | 0,000235614 | 0,003418345 |
| ENSDARG00000021573 | slc16a7 | -1,013137332 | 1,921851881 | 21,39773706 | 0,000445829 | 0,005206482 |
| ENSDARG00000089952 | si:ch211-147k9.8 | -1,014450461 | -1,735800526 | 10,33098852 | 0,00659157 | 0,03133547 |
| ENSDARG00000071558 | fblim1 | -1,015297291 | 3,58847564 | 62,60661169 | 2,14505E-06 | 0,000185316 |
| ENSDARG00000074036 | ky | -1,017748013 | 0,570556465 | 14,89647826 | 0,001887468 | 0,013478473 |
| ENSDARG00000031876 | pard6ga | -1,018228071 | -1,370237143 | 11,52001067 | 0,004644398 | 0,02457902 |
| ENSDARG00000019260 | dhrs9 | -1,018440859 | 4,588200803 | 28,3849391 | 0,000177359 | 0,002869586 |
| ENSDARG00000052515 | calcoco2 | -1,019077507 | 3,043521186 | 61,19954925 | 2,43397E-06 | 0,000199726 |
| ENSDARG00000036722 | slc9a3.2 | -1,019445416 | 2,511456414 | 26,69750712 | 0,000168823 | 0,002797724 |
| ENSDARG00000102129 | crygn1 | -1,020307884 | 2,801424594 | 20,3418109 | 0,000623145 | 0,006463895 |
| ENSDARG00000092469 | fndc7rs1 | -1,021805541 | 1,673715068 | 18,64611029 | 0,000793987 | 0,007561372 |
| ENSDARG00000036140 | crybgx | -1,021845686 | 7,589055396 | 19,97172877 | 0,000864501 | 0,007968089 |
| ENSDARG00000054837 | zgc:136870 | -1,022081114 | 0,860767201 | 14,71992509 | 0,001972521 | 0,013836834 |
| ENSDARG00000101695 | si:dkey-26m3.3 | -1,026646859 | 1,31156749 | 14,56677056 | 0,002094891 | 0,014376627 |
| ENSDARG00000075067 | aipl1 | -1,026668251 | 2,183753763 | 27,62326851 | 0,000142972 | 0,002529517 |
| ENSDARG00000079216 | dpep2 | -1,026737672 | 1,090634992 | 25,92743966 | 0,000191181 | 0,003010239 |
| ENSDARG00000044775 | fut7 | -1,028597079 | -1,153803438 | 15,09840076 | 0,001795339 | 0,012998887 |
| ENSDARG00000044365 | angptl3 | -1,028738045 | 4,759164112 | 57,14489719 | 3,84044E-06 | 0,000260127 |
| ENSDARG00000090899 | sycn.3 | -1,029165054 | 2,638516327 | 8,772645417 | 0,012803379 | 0,049732186 |
| ENSDARG00000094272 | map3k19 | -1,02924249 | 0,49958014 | 24,83498509 | 0,000232161 | 0,003381107 |

| | | | | | | |
|---|---|---|---|---|---|---|
| ENSDARG00000104717 | tbxa2r | -1,031269764 | -1,025290714 | 13,24695499 | 0,002884386 | 0,017876325 |
| ENSDARG00000043249 | irf1b | -1,033016171 | 2,954290638 | 11,67533483 | 0,005433743 | 0,027299761 |
| ENSDARG00000102835 | CABZ01021435.1 | -1,033230065 | -0,041192304 | 11,41404304 | 0,004787883 | 0,025065476 |
| ENSDARG00000069806 | sstr2b | -1,033267878 | -0,211256471 | 12,97711454 | 0,003100104 | 0,018735637 |
| ENSDARG00000005894 | slc7a9 | -1,033511457 | 2,645405441 | 13,61965484 | 0,003104909 | 0,018735844 |
| ENSDARG00000007582 | si:ch1073-155h21.1 | -1,033534057 | 0,074903931 | 12,73538147 | 0,003309329 | 0,019521713 |
| ENSDARG00000099509 | btr23 | -1,035664306 | 1,199524778 | 9,534300038 | 0,008986673 | 0,039016196 |
| ENSDARG00000104966 | si:dkey-238k10.3 | -1,037679163 | -0,100278972 | 19,04076083 | 0,000725035 | 0,007151088 |
| ENSDARG00000071495 | sult3st5 | -1,037803116 | -0,245639951 | 11,07121805 | 0,00528853 | 0,026786639 |
| ENSDARG00000091993 | dicp1.1 | -1,039005386 | -0,546367922 | 15,99478014 | 0,001444278 | 0,011160238 |
| ENSDARG00000088301 | si:dkey-254e13.6 | -1,039913997 | -0,302660128 | 21,04878913 | 0,000477599 | 0,005472546 |
| ENSDARG00000056339 | stk31 | -1,040584819 | 0,635830396 | 10,25514835 | 0,006872701 | 0,032209544 |
| ENSDARG00000078250 | zgc:194398 | -1,041673124 | 4,150757195 | 21,56849867 | 0,000589242 | 0,006233873 |
| ENSDARG00000041413 | adgrg11 | -1,043134094 | 1,559666207 | 18,28827489 | 0,000859229 | 0,007943787 |
| ENSDARG00000100292 | fam166b | -1,04496125 | -0,070786395 | 25,20216425 | 0,000217351 | 0,003274826 |
| ENSDARG00000099738 | ccr9b | -1,045698804 | -0,081658139 | 17,74040865 | 0,000964426 | 0,008618102 |
| ENSDARG00000105095 | AL929028.1 | -1,047117276 | 1,042608504 | 19,12037732 | 0,000712764 | 0,007064387 |
| ENSDARG00000114349 | si:dkey-20i20.15 | -1,047507167 | 0,293466482 | 39,8360813 | 2,40317E-05 | 0,000842546 |
| ENSDARG00000044612 | c1qb | -1,048053994 | -0,296377924 | 20,00383615 | 0,000591508 | 0,00624919 |
| ENSDARG00000077382 | hcn5 | -1,048225093 | 1,120800794 | 42,01185644 | 1,82939E-05 | 0,000694561 |
| ENSDARG00000056643 | slc22a7b.1 | -1,048816662 | 2,226435937 | 55,22697403 | 4,28173E-06 | 0,000275353 |
| ENSDARG00000100900 | si:ch211-183d5.2 | -1,050048787 | -0,828423588 | 10,00663777 | 0,007277548 | 0,033530326 |
| ENSDARG00000060953 | si:ch1073-214b20.2 | -1,050408554 | -1,656443543 | 11,54860092 | 0,004606542 | 0,024486227 |
| ENSDARG00000061717 | cytip | -1,051362403 | 0,036750889 | 15,01930476 | 0,001830794 | 0,013205482 |
| ENSDARG00000075141 | gprc5bb | -1,052219826 | 1,364391848 | 49,20779651 | 7,97847E-06 | 0,000396224 |
| ENSDARG00000078093 | zgc:172065 | -1,052251221 | 1,150557538 | 17,0553586 | 0,001126679 | 0,009488807 |
| ENSDARG00000116915 | si:ch211-133h13.1 | -1,052496318 | 0,7333155 | 30,12614711 | 9,51936E-05 | 0,001954585 |
| ENSDARG00000032206 | cthl | -1,052636779 | 2,70692097 | 22,267005 | 0,000418663 | 0,004997324 |
| ENSDARG00000058045 | tlr21 | -1,053608057 | 0,748256514 | 10,23735753 | 0,006959894 | 0,032479083 |
| ENSDARG00000056458 | lhfpl5b | -1,053894275 | -1,17391757 | 10,571159 | 0,006131749 | 0,029764609 |
| ENSDARG00000075515 | cd37 | -1,054475738 | -0,095903125 | 23,23537546 | 0,000311866 | 0,004097397 |
| ENSDARG00000089582 | si:dkey-265e15.2 | -1,054623867 | 1,625741048 | 50,78421603 | 6,73975E-06 | 0,000360514 |
| ENSDARG00000035798 | gngt1 | -1,054867098 | 6,626908811 | 43,05669176 | 2,18159E-05 | 0,000786165 |
| ENSDARG00000098620 | si:ch211-188p14.2 | -1,057822082 | 0,632010223 | 8,558655634 | 0,012011762 | 0,047552703 |
| ENSDARG00000055437 | METTL21C | -1,057877416 | 0,799828321 | 16,05119641 | 0,001424979 | 0,011066309 |
| ENSDARG00000093957 | si:dkey-251i10.2 | -1,058279837 | 4,606002572 | 19,24380049 | 0,001057184 | 0,009102362 |
| ENSDARG00000117363 | CABZ01071845.1 | -1,058386703 | -0,233893991 | 16,17671021 | 0,001383102 | 0,010850559 |
| ENSDARG00000096863 | si:ch73-380l3.4 | -1,058902018 | -1,532741598 | 9,39378033 | 0,00881332 | 0,038513484 |
| ENSDARG00000095275 | si:dkey-269p2.1 | -1,059025208 | -0,847685721 | 13,70564206 | 0,002556269 | 0,016499358 |
| ENSDARG00000075929 | si:dkey-219c10.4 | -1,059439772 | 1,091212549 | 24,24642211 | 0,000258396 | 0,00362105 |
| ENSDARG00000077096 | fndc7a | -1,059852333 | 2,997044314 | 27,68927221 | 0,000154403 | 0,002646511 |
| ENSDARG00000070666 | rhol | -1,059864888 | 3,267232981 | 18,89010127 | 0,000949672 | 0,008512836 |
| ENSDARG00000061710 | map3k8 | -1,059993801 | 0,898345185 | 41,5146029 | 1,94517E-05 | 0,000722933 |
| ENSDARG00000097974 | CU855685.1 | -1,063418633 | -0,405411036 | 13,17773455 | 0,002938021 | 0,018086326 |
| ENSDARG00000077504 | si:ch211-103n10.5 | -1,066064374 | 1,282341936 | 9,553023537 | 0,009074507 | 0,039270868 |
| ENSDARG00000053502 | cryaa | -1,067040976 | 5,398778482 | 52,79625791 | 6,97656E-06 | 0,000368022 |
| ENSDARG00000109371 | pdgfaa | -1,067677193 | -0,032383198 | 11,80663509 | 0,004280579 | 0,023338792 |
| ENSDARG00000045644 | ca12 | -1,068162065 | 1,016615703 | 28,84804756 | 0,000116802 | 0,00224711 |
| ENSDARG00000063475 | abcg1 | -1,068337111 | 3,881286455 | 48,75591598 | 8,72358E-06 | 0,000421869 |
| ENSDARG00000089429 | si:dkey-205h13.2 | -1,06841227 | 7,445650292 | 38,60647247 | 3,84959E-05 | 0,001139188 |
| ENSDARG00000105528 | si:dkey-86l18.10 | -1,068634845 | 2,416207977 | 27,16983918 | 0,000158423 | 0,002696535 |
| ENSDARG00000103650 | si:ch73-329n5.1 | -1,069551365 | 0,044260172 | 14,3584312 | 0,00216087 | 0,014688524 |
| ENSDARG00000098570 | si:ch211-8c17.2 | -1,071057686 | -1,12854199 | 14,7985618 | 0,001934103 | 0,013677847 |
| ENSDARG00000021208 | serpind1 | -1,073617477 | 3,550184718 | 62,18833846 | 2,2266E-06 | 0,000190111 |
| ENSDARG00000097208 | alkbh3 | -1,073697146 | 3,868883728 | 86,06462054 | 3,49293E-07 | 6,50138E-05 |
| ENSDARG00000101675 | zgc:123107 | -1,073728187 | -1,905707518 | 13,58787675 | 0,002636198 | 0,016839594 |
| ENSDARG00000000656 | psmb9a | -1,073951589 | -0,821930783 | 14,06143654 | 0,002331279 | 0,015492686 |
| ENSDARG00000015065 | c4 | -1,076200414 | 1,197899483 | 17,87721733 | 0,00093535 | 0,008414106 |
| ENSDARG00000099889 | dusp27 | -1,076379524 | 5,235962306 | 68,3844779 | 1,43747E-06 | 0,000145321 |
| ENSDARG00000079926 | syt15 | -1,077476277 | 1,22788161 | 28,35955174 | 0,000126516 | 0,002376098 |
| ENSDARG00000069951 | eef1a1l2 | -1,07877913 | 4,373645135 | 93,01703657 | 2,2163E-07 | 4,91838E-05 |
| ENSDARG00000060070 | adcy7 | -1,079743427 | 4,383848639 | 165,5524043 | 6,76007E-09 | 4,99242E-06 |
| ENSDARG00000058005 | hgd | -1,080657209 | 6,057104732 | 105,8060905 | 1,03301E-07 | 2,87701E-05 |
| ENSDARG00000079532 | zgc:194242 | -1,081093589 | 0,155444679 | 11,91522798 | 0,004151439 | 0,022835909 |
| ENSDARG00000068586 | CU302253.1 | -1,081274297 | 4,744138496 | 20,62890664 | 0,000820041 | 0,007710065 |
| ENSDARG00000026663 | tlr19 | -1,081311655 | -0,398335757 | 11,62542105 | 0,004506585 | 0,02411165 |
| ENSDARG00000102442 | folr | -1,081323935 | 3,326250262 | 12,20666816 | 0,005017811 | 0,025850083 |
| ENSDARG00000015752 | zap70 | -1,082184523 | -0,907623274 | 9,976198561 | 0,007346075 | 0,033764513 |
| ENSDARG00000087181 | cbx7b | -1,082927214 | 0,016546206 | 11,71174076 | 0,004397249 | 0,023754276 |
| ENSDARG00000101665 | pcdh1g11 | -1,084927117 | 1,139759744 | 31,73704823 | 7,4202E-05 | 0,001681962 |
| ENSDARG00000061292 | pi15b | -1,086218588 | 1,11043273 | 34,13151873 | 5,20927E-05 | 0,001364945 |

| | | | | | | |
|---|---|---|---|---|---|---|
| ENSDARG00000061968 | myo1ha | -1,087819237 | -0,474352376 | 22,95168 | 0,00032909 | 0,004251802 |
| ENSDARG00000075248 | zgc:172133 | -1,088739908 | -1,277185152 | 9,181670126 | 0,009430351 | 0,04039737 |
| ENSDARG00000032010 | slc15a2 | -1,08963739 | 4,31175789 | 49,69570716 | 8,38951E-06 | 0,000410485 |
| ENSDARG00000076800 | tnfrsf18 | -1,089693449 | -1,972244578 | 9,851249925 | 0,007635326 | 0,034717608 |
| ENSDARG00000010207 | smad3b | -1,090604549 | 3,346952583 | 163,2752359 | 7,36593E-09 | 5,11017E-06 |
| ENSDARG00000086518 | si:ch211-155o21.3 | -1,092159908 | -0,408305456 | 22,64990371 | 0,000348627 | 0,00442432 |
| ENSDARG00000008790 | actr3b | -1,092463449 | 2,041159725 | 32,06637942 | 7,05977E-05 | 0,001622755 |
| ENSDARG00000117607 | FP102158.1 | -1,092673075 | -1,195298849 | 13,51759005 | 0,002685278 | 0,017024854 |
| ENSDARG00000011671 | pde6b | -1,092869565 | 4,9710694 | 38,19416079 | 4,46299E-05 | 0,001256775 |
| ENSDARG00000079727 | selenop2 | -1,094518642 | 5,50027216 | 53,53252965 | 6,68326E-06 | 0,000358329 |
| ENSDARG00000105522 | si:ch211-196f2.3 | -1,095341322 | 3,897390494 | 11,45555048 | 0,006561649 | 0,031250759 |
| ENSDARG00000112287 | zgc:66156 | -1,096334318 | 6,289210453 | 28,19007214 | 0,000213005 | 0,003231642 |
| ENSDARG00000070440 | atp6v1c2 | -1,0979093 | -1,077112592 | 14,30410924 | 0,002190933 | 0,014834895 |
| ENSDARG00000011701 | ctsll | -1,098571774 | -0,207740851 | 20,18838914 | 0,000569287 | 0,006104572 |
| ENSDARG00000079031 | si:ch211-22d5.2 | -1,099167902 | 2,399683073 | 17,27178047 | 0,001248142 | 0,010161797 |
| ENSDARG00000087388 | si:dkey-29b11.3 | -1,099864246 | -0,269731005 | 17,52503693 | 0,00101234 | 0,008852756 |
| ENSDARG00000041828 | si:dkeyp-28d2.4 | -1,100298498 | -1,107849204 | 12,30800869 | 0,003720554 | 0,021133326 |
| ENSDARG00000116457 | zmp:0000001088 | -1,10251177 | 1,245799772 | 20,18598737 | 0,00056957 | 0,006104747 |
| ENSDARG00000100439 | zgc:153383 | -1,102652095 | 1,542088243 | 16,58016316 | 0,001312529 | 0,010516735 |
| ENSDARG00000096721 | si:rp71-1c10.11 | -1,104547858 | 3,998710253 | 47,70199699 | 1,02897E-05 | 0,000467407 |
| ENSDARG00000087706 | dicp3.3 | -1,104643652 | -0,238566067 | 17,58705813 | 0,000998266 | 0,00876988 |
| ENSDARG00000002193 | rho | -1,109163629 | 11,03183866 | 24,5031785 | 0,000309432 | 0,004078372 |
| ENSDARG00000086256 | si:ch211-236p5.2 | -1,110100506 | 1,990023742 | 32,67126173 | 6,44898E-05 | 0,001539551 |
| ENSDARG00000095475 | BX649377.1 | -1,11054025 | -0,763743645 | 8,923846933 | 0,010249179 | 0,042654916 |
| ENSDARG00000070229 | zgc:158258 | -1,111024396 | 2,703174878 | 21,19860565 | 0,000543227 | 0,005916569 |
| ENSDARG00000117258 | FP236157.4 | -1,111442245 | 1,003212005 | 19,89332416 | 0,000605293 | 0,006339241 |
| ENSDARG00000038729 | s100z | -1,11160568 | 2,150375693 | 23,96066473 | 0,000284625 | 0,003858032 |
| ENSDARG00000026017 | xpnpep2 | -1,112368758 | 3,011919962 | 20,57739489 | 0,000656128 | 0,006691555 |
| ENSDARG00000045414 | elovl2 | -1,114679694 | 4,87793552 | 15,76110908 | 0,002464356 | 0,016110501 |
| ENSDARG00000086130 | si:ch211-42i6.2 | -1,115495282 | -1,020823733 | 15,99446611 | 0,001444387 | 0,011160238 |
| ENSDARG00000101049 | CABZ01075274.1 | -1,115901033 | 0,766805767 | 27,57755169 | 0,000144072 | 0,00253775 |
| ENSDARG00000016161 | ccdc170 | -1,116168043 | 0,113849462 | 27,57631283 | 0,000144102 | 0,00253775 |
| ENSDARG00000060319 | scn4bb | -1,116252724 | 0,960134511 | 20,29225968 | 0,000557205 | 0,00602581 |
| ENSDARG00000039145 | plaub | -1,116433741 | 0,972546201 | 18,60758836 | 0,000796225 | 0,007570093 |
| ENSDARG00000076790 | crygm2d16 | -1,116978365 | 7,793563728 | 41,89859252 | 2,4636E-05 | 0,000855867 |
| ENSDARG00000063347 | hrc | -1,117642121 | -0,447604009 | 13,20802061 | 0,002914413 | 0,018003934 |
| ENSDARG00000076306 | loxl3a | -1,118916788 | 2,629004656 | 115,329066 | 6,16307E-08 | 2,10593E-05 |
| ENSDARG00000105352 | zgc:165555 | -1,119479312 | -1,443850722 | 8,920539918 | 0,010260206 | 0,042693049 |
| ENSDARG00000044318 | itgb7 | -1,119564758 | 1,185567436 | 37,4859438 | 3,26954E-05 | 0,001024748 |
| ENSDARG00000117555 | FP236157.6 | -1,120395556 | 2,106939591 | 18,66479699 | 0,000868262 | 0,007989547 |
| ENSDARG00000105450 | si:ch211-63p21.1 | -1,120921356 | 4,280885913 | 24,19605862 | 0,000383636 | 0,004694256 |
| ENSDARG00000022183 | gsto1 | -1,122427776 | 3,365132542 | 13,46571829 | 0,003674296 | 0,020946047 |
| ENSDARG00000100312 | hoxa13a | -1,122972427 | 0,649913159 | 21,81247339 | 0,000410243 | 0,004927926 |
| ENSDARG00000113315 | zgc:153932 | -1,123505428 | 2,106362769 | 25,62681333 | 0,000208158 | 0,003187666 |
| ENSDARG00000102225 | si:ch211-205a14.7 | -1,12565101 | -0,885054266 | 11,4544226 | 0,00473261 | 0,02489839 |
| ENSDARG00000070362 | cx30.9 | -1,125678299 | -1,11898744 | 11,16086831 | 0,005151982 | 0,026369209 |
| ENSDARG00000059361 | slc39a4 | -1,12597746 | 0,550480017 | 17,76652844 | 0,000958795 | 0,008583992 |
| ENSDARG00000094045 | si:dkeyp-73d8.8 | -1,126934611 | 0,74019022 | 17,50932543 | 0,001015942 | 0,008877469 |
| ENSDARG00000102119 | si:ch73-359m17.2 | -1,129455218 | 1,492532173 | 36,20173409 | 3,89277E-05 | 0,001144181 |
| ENSDARG00000052413 | crfb12 | -1,129767157 | 2,958869364 | 80,77924428 | 5,04621E-07 | 8,02278E-05 |
| ENSDARG00000079307 | si:dkey-205h13.1 | -1,131592254 | 3,444546454 | 32,49052561 | 8,02363E-05 | 0,001759037 |
| ENSDARG00000053973 | fetub | -1,135420199 | 9,704121701 | 84,85557705 | 3,79286E-07 | 6,91397E-05 |
| ENSDARG00000039211 | zgc:77439 | -1,136408723 | 5,971990799 | 130,5522199 | 2,90973E-08 | 1,33231E-05 |
| ENSDARG00000090882 | si:rp71-36a1.5 | -1,136769557 | -0,03583728 | 16,65897402 | 0,001234886 | 0,01008615 |
| ENSDARG00000089149 | gpx2 | -1,137114349 | 0,918421785 | 30,13621147 | 9,50427E-05 | 0,001953239 |
| ENSDARG00000056018 | itga6l | -1,137633986 | 2,145186958 | 9,46049144 | 0,010196259 | 0,042559393 |
| ENSDARG00000079824 | ninj2 | -1,138753219 | 0,058573521 | 17,24167215 | 0,001079626 | 0,009250355 |
| ENSDARG00000061383 | serpinf2b | -1,139101777 | 5,726162389 | 87,06049916 | 3,32691E-07 | 6,24314E-05 |
| ENSDARG00000030626 | lrit2 | -1,139390759 | 2,842286408 | 63,30582081 | 2,01628E-06 | 0,00018131 |
| ENSDARG00000076321 | col28a2a | -1,141686851 | 3,793441004 | 53,61149391 | 5,29898E-06 | 0,000305579 |
| ENSDARG00000018351 | hpda | -1,141730853 | 5,29034461 | 47,54455406 | 1,46157E-05 | 0,000601195 |
| ENSDARG00000079829 | ccr2 | -1,141837826 | -1,943282906 | 8,31678317 | 0,012526245 | 0,048925787 |
| ENSDARG00000062598 | ltc4s | -1,142214263 | -0,431673153 | 9,112469617 | 0,009642414 | 0,040953282 |
| ENSDARG00000070038 | rbp2a | -1,142290627 | 6,712398539 | 28,146787 | 0,000219213 | 0,003284466 |
| ENSDARG00000045299 | vmo1b | -1,143083376 | 3,468133292 | 97,91854606 | 1,63696E-07 | 3,91591E-05 |
| ENSDARG00000097592 | si:dkey-40c23.1 | -1,143490021 | -0,647332958 | 18,63220169 | 0,00079197 | 0,007545307 |
| ENSDARG00000056719 | slc6a19b | -1,14379472 | 4,3843954 | 21,87465643 | 0,00063033 | 0,006506214 |
| ENSDARG00000094408 | MFAP4 (1 of many) | -1,144560349 | 1,351605721 | 33,78047947 | 5,48016E-05 | 0,001396124 |
| ENSDARG00000040921 | zmp:0000000606 | -1,145521742 | -0,289774293 | 13,82518861 | 0,002477983 | 0,016171875 |
| ENSDARG00000094041 | krt17 | -1,145639794 | 9,210724015 | 78,44102572 | 5,97818E-07 | 8,55403E-05 |
| ENSDARG00000093006 | rarres3 | -1,145725775 | -0,528657134 | 13,85358099 | 0,002459799 | 0,016096514 |

| | | | | | | |
|---|---|---|---|---|---|---|
| ENSDARG00000039502 | eef1a1a | -1,145771407 | 6,297983131 | 103,7086023 | 1,16396E-07 | 3,06294E-05 |
| ENSDARG00000019686 | fgl2b | -1,147311775 | 2,65150239 | 24,24030468 | 0,000296089 | 0,003966269 |
| ENSDARG00000101514 | ftr64 | -1,147655772 | -0,232589368 | 30,02581531 | 9,67126E-05 | 0,001978018 |
| ENSDARG00000035458 | atp2a1l | -1,148848558 | 10,67252051 | 74,97628292 | 7,74821E-07 | 9,80041E-05 |
| ENSDARG00000099152 | si:dkey-190j3.2 | -1,148986528 | 0,495412369 | 20,26543501 | 0,000560296 | 0,006047819 |
| ENSDARG00000094067 | si:dkey-121n8.7 | -1,148988851 | 1,504503022 | 48,80698749 | 8,33395E-06 | 0,000409436 |
| ENSDARG00000108006 | CU693481.1 | -1,149077955 | 2,616552923 | 34,19838588 | 5,26075E-05 | 0,001373313 |
| ENSDARG00000010936 | abcb4 | -1,150163001 | 5,952632668 | 12,65485969 | 0,005267574 | 0,026704128 |
| ENSDARG00000011083 | si:dkey-42l23.2 | -1,150629778 | -1,246155346 | 16,25777936 | 0,001356808 | 0,010726095 |
| ENSDARG00000075626 | zgc:172090 | -1,151317723 | -1,449601609 | 8,849452806 | 0,010500617 | 0,043370219 |
| ENSDARG00000043818 | si:ch211-119c20.2 | -1,152399268 | 3,709084508 | 34,68868705 | 6,18653E-05 | 0,001504482 |
| ENSDARG00000104365 | BX323064.2 | -1,152879279 | -0,415798506 | 18,50911486 | 0,000813514 | 0,007670758 |
| ENSDARG00000093423 | swsap1 | -1,153074544 | 0,051422407 | 27,60458849 | 0,00014342 | 0,002531583 |
| ENSDARG00000086569 | zgc:172051 | -1,153814406 | 4,511512326 | 60,01330466 | 3,03738E-06 | 0,000231011 |
| ENSDARG00000068726 | masp1 | -1,154214464 | 2,579481151 | 91,43646079 | 2,45116E-07 | 5,29404E-05 |
| ENSDARG00000100214 | si:dkey-247i3.6 | -1,154725243 | -1,012674194 | 9,246667114 | 0,00923609 | 0,039799953 |
| ENSDARG00000077479 | msh5 | -1,154876187 | -1,930671689 | 13,7405618 | 0,00253311 | 0,016387055 |
| ENSDARG00000087532 | si:ch211-278p9.1 | -1,155299158 | 1,841404376 | 58,71297548 | 3,06142E-06 | 0,000231011 |
| ENSDARG00000041699 | piwil1 | -1,155706467 | 1,292160846 | 18,46098356 | 0,000845663 | 0,007861206 |
| ENSDARG00000079347 | zgc:194659 | -1,156776518 | 4,10524205 | 56,48818412 | 4,08882E-06 | 0,000265936 |
| ENSDARG00000095139 | CR388373.2 | -1,158294 | -0,432111468 | 29,25035477 | 0,000109443 | 0,002156266 |
| ENSDARG00000000380 | pde6a | -1,162231614 | 4,813106699 | 71,70009065 | 1,1026E-06 | 0,00012435 |
| ENSDARG00000098538 | CABZ01112049.1 | -1,16374937 | 1,874360285 | 8,836788806 | 0,012249436 | 0,048127439 |
| ENSDARG00000029890 | si:dkeyp-86f7.4 | -1,164130445 | 0,412039236 | 8,984184714 | 0,010521883 | 0,043418889 |
| ENSDARG00000099690 | vaspa | -1,164513374 | 0,728916535 | 33,98170394 | 5,32295E-05 | 0,001378547 |
| ENSDARG00000093068 | c3b.1 | -1,165457522 | 5,00597349 | 135,1070568 | 2,36082E-08 | 1,14997E-05 |
| ENSDARG00000097266 | AL954322.3 | -1,166246488 | 0,154040024 | 23,09424141 | 0,000320302 | 0,00417596 |
| ENSDARG00000054534 | ACOT12 | -1,166318685 | 2,563216669 | 18,81084413 | 0,000928464 | 0,008388424 |
| ENSDARG00000090969 | cbln18 | -1,167166743 | 2,353872639 | 33,26103409 | 5,95339E-05 | 0,001471889 |
| ENSDARG00000105015 | CU459147.1 | -1,167410536 | -1,739450889 | 11,91934222 | 0,004146636 | 0,022816566 |
| ENSDARG00000055359 | sp100.1 | -1,168825978 | 0,467636301 | 19,95626968 | 0,000597397 | 0,006292298 |
| ENSDARG00000093316 | adgrf8 | -1,16900983 | 3,196946481 | 29,91596696 | 0,000120482 | 0,00229096 |
| ENSDARG00000054588 | cox6a2 | -1,169420432 | 2,421822039 | 26,79329747 | 0,00017809 | 0,00287339 |
| ENSDARG00000039579 | cfd | -1,171701277 | 6,077322365 | 52,5561965 | 8,51275E-06 | 0,000414662 |
| ENSDARG00000102679 | FP074874.1 | -1,173007049 | 3,196373667 | 30,77019409 | 0,000104648 | 0,002083309 |
| ENSDARG00000091140 | pik3r6b | -1,174750705 | -1,683238093 | 8,524934206 | 0,011684823 | 0,046694422 |
| ENSDARG00000071724 | ankha | -1,176425493 | 2,585161705 | 32,03383608 | 7,51955E-05 | 0,001693248 |
| ENSDARG00000068923 | umodl1 | -1,17810486 | 0,258227989 | 20,35699889 | 0,000549823 | 0,005967087 |
| ENSDARG00000007086 | aqp10a | -1,17828202 | 2,159211815 | 42,83871998 | 1,65397E-05 | 0,000656258 |
| ENSDARG00000092303 | si:dkey-11o15.8 | -1,178472748 | -1,059199647 | 8,309877931 | 0,012555345 | 0,048993022 |
| ENSDARG00000099558 | NPC1L1 | -1,178720503 | 3,960755676 | 27,80783414 | 0,000203337 | 0,003143273 |
| ENSDARG00000056915 | si:ch211-237l4.6 | -1,179212676 | 2,786237932 | 46,94818164 | 1,02402E-05 | 0,000466083 |
| ENSDARG00000104398 | si:ch73-343g19.4 | -1,181107233 | -1,189287276 | 10,06840729 | 0,007140756 | 0,033093211 |
| ENSDARG00000055683 | pimr191 | -1,181627586 | -0,979934303 | 27,03318622 | 0,000157955 | 0,002691759 |
| ENSDARG00000098063 | alp3 | -1,181764452 | 2,001667904 | 9,685464401 | 0,009509584 | 0,040617987 |
| ENSDARG00000059020 | thap1 | -1,182295404 | 2,864984758 | 68,99585948 | 1,24331E-06 | 0,000135836 |
| ENSDARG00000054160 | zgc:113625 | -1,183291328 | 0,977951112 | 9,295438465 | 0,009986726 | 0,041951579 |
| ENSDARG00000039150 | lgmn | -1,183391686 | 8,205936544 | 62,49765633 | 2,38037E-06 | 0,000198168 |
| ENSDARG00000087907 | ndufa4l2b | -1,184157222 | -1,24452541 | 18,73523957 | 0,000774441 | 0,007449598 |
| ENSDARG00000069710 | vwa10.2 | -1,184945208 | -0,047777996 | 25,17087365 | 0,00021857 | 0,003279122 |
| ENSDARG00000105079 | BX649411.2 | -1,185343326 | 1,010286357 | 34,54322279 | 4,91098E-05 | 0,001319359 |
| ENSDARG00000068181 | dpep1 | -1,185416082 | 3,539494684 | 13,90127884 | 0,003481315 | 0,020157111 |
| ENSDARG00000040277 | fbxo32 | -1,185482911 | 2,896671818 | 20,08260169 | 0,00075009 | 0,00732034 |
| ENSDARG00000091249 | CR381686.1 | -1,187054058 | -0,118835155 | 26,64198668 | 0,000168885 | 0,002797724 |
| ENSDARG00000028485 | cabp5b | -1,18739625 | 3,145420619 | 27,36405205 | 0,0001902 | 0,003000985 |
| ENSDARG00000087195 | rorc | -1,190288788 | -0,385495074 | 18,09086779 | 0,000891941 | 0,008138736 |
| ENSDARG00000074547 | si:ch211-240l19.8 | -1,190946121 | -1,597382017 | 9,219675038 | 0,009316193 | 0,040053507 |
| ENSDARG00000076139 | si:dkey-71p21.13 | -1,191325954 | -0,531424035 | 29,71269951 | 0,000101636 | 0,002044684 |
| ENSDARG00000007465 | tdrd1 | -1,191650807 | 0,952660512 | 24,37818787 | 0,000252238 | 0,003575685 |
| ENSDARG00000075829 | kiss1 | -1,194651987 | -1,231485214 | 15,24259377 | 0,001732723 | 0,012702198 |
| ENSDARG00000105714 | CABZ01083442.1 | -1,197359945 | 3,308414402 | 9,106470479 | 0,012761809 | 0,049612643 |
| ENSDARG00000095184 | si:ch73-52p7.1 | -1,198361131 | 0,541771235 | 23,31545896 | 0,000307194 | 0,00405588 |
| ENSDARG00000089852 | si:dkey-51d8.1 | -1,198366627 | -1,392882601 | 15,56818951 | 0,001600387 | 0,012036552 |
| ENSDARG00000052012 | rtn4rl2a | -1,199450431 | 3,207923879 | 9,200534921 | 0,012314293 | 0,048315924 |
| ENSDARG00000094854 | ms4a17a.9 | -1,199875259 | 4,080181284 | 22,55475031 | 0,000545482 | 0,005929851 |
| ENSDARG00000028912 | si:dkey-10h3.2 | -1,200332026 | -2,468933259 | 9,335896526 | 0,008976925 | 0,039016196 |
| ENSDARG00000074476 | odam | -1,200338807 | 3,15924874 | 10,8762368 | 0,007547669 | 0,034403194 |
| ENSDARG00000069944 | p2ry13 | -1,200407906 | -0,539445136 | 18,27408902 | 0,000856557 | 0,007936068 |
| ENSDARG00000091916 | ugt5b4 | -1,201644937 | 4,017181633 | 50,86072469 | 7,8471E-06 | 0,000393475 |
| ENSDARG00000074851 | s1pr4 | -1,202901397 | -1,728130051 | 8,732195663 | 0,010911636 | 0,044537526 |
| ENSDARG00000006811 | sult1st6 | -1,203180985 | 4,606977078 | 28,52257928 | 0,000205853 | 0,003167713 |

| | | | | | | |
|---|---|---|---|---|---|---|
| ENSDARG00000105104 | si:dkey-223p19.1 | -1,205889252 | 0,636097862 | 22,56169787 | 0,000354588 | 0,004469428 |
| ENSDARG00000059026 | zgc:123217 | -1,206904544 | 1,521449472 | 20,36561655 | 0,000571882 | 0,006123791 |
| ENSDARG00000074684 | mlsl | -1,207225386 | 2,56048337 | 55,73695309 | 4,0723E-06 | 0,000265616 |
| ENSDARG00000030616 | nfe2l1a | -1,209185311 | 4,380918912 | 57,75682182 | 3,996E-06 | 0,000265616 |
| ENSDARG00000038891 | AL954146.1 | -1,209331041 | 1,615170922 | 19,16810314 | 0,00075375 | 0,007343126 |
| ENSDARG00000077643 | lypd6b | -1,209972107 | 2,135012845 | 80,12203814 | 5,29011E-07 | 8,14392E-05 |
| ENSDARG00000026726 | anxa1a | -1,210284851 | 7,134523564 | 76,87728348 | 7,45786E-07 | 9,64635E-05 |
| ENSDARG00000002259 | ca15c | -1,21059912 | 2,471218033 | 19,30354528 | 0,000843392 | 0,007849028 |
| ENSDARG00000110019 | CABZ01079267.1 | -1,21130354 | 1,451006615 | 16,34202235 | 0,001447469 | 0,011180278 |
| ENSDARG00000096959 | si:dkey-29h14.10 | -1,21133965 | -0,741168363 | 20,88736991 | 0,00049342 | 0,005559231 |
| ENSDARG00000095633 | si:ch211-133l5.7 | -1,211519486 | 4,396302385 | 15,62414156 | 0,002537784 | 0,016412436 |
| ENSDARG00000105568 | si:ch211-194p6.10 | -1,212556648 | 2,186220518 | 104,5760842 | 1,10761E-07 | 2,94856E-05 |
| ENSDARG00000036968 | si:ch1073-416d2.3 | -1,213318077 | 1,816641022 | 51,23678843 | 6,42609E-06 | 0,000347799 |
| ENSDARG00000060609 | cd109 | -1,216244383 | 3,189550727 | 59,13136424 | 2,94389E-06 | 0,000228404 |
| ENSDARG00000117560 | AL954145.2 | -1,216644963 | -1,776910038 | 9,030245082 | 0,009901634 | 0,041670588 |
| ENSDARG00000063059 | abcg8 | -1,217679058 | 1,904391821 | 22,78979486 | 0,000365029 | 0,004542419 |
| ENSDARG00000074613 | si:ch211-240l19.6 | -1,217797713 | 3,701159671 | 17,41496185 | 0,001568019 | 0,011855426 |
| ENSDARG00000102602 | si:ch211-220m17.4 | -1,222124727 | -1,029728534 | 9,410028434 | 0,00876802 | 0,038352129 |
| ENSDARG00000104153 | prkd4 | -1,22292208 | -1,304138496 | 14,79039598 | 0,001938052 | 0,013694371 |
| ENSDARG00000063626 | ddx21 | -1,224886698 | 7,707660007 | 55,0399648 | 5,72645E-06 | 0,000326123 |
| ENSDARG00000037790 | pvalb8 | -1,224989913 | 6,977209173 | 61,84308024 | 3,06708E-06 | 0,000231011 |
| ENSDARG00000044973 | krt98 | -1,226316776 | 0,555036769 | 25,17933788 | 0,00021824 | 0,003277079 |
| ENSDARG00000104329 | si:ch73-281k2.5 | -1,226340052 | 3,934298548 | 54,96672541 | 4,99392E-06 | 0,000297736 |
| ENSDARG00000035018 | thy1 | -1,226923101 | 4,365762237 | 27,13522341 | 0,00025354 | 0,003583058 |
| ENSDARG00000070000 | txnipb | -1,228051359 | 5,423009172 | 105,0726281 | 1,07677E-07 | 2,90019E-05 |
| ENSDARG00000077875 | slc2a5 | -1,228299084 | -0,673817942 | 20,5135482 | 0,000532436 | 0,005857561 |
| ENSDARG00000069542 | si:dkey-8e10.2 | -1,22891201 | -1,241520687 | 13,01794985 | 0,003066291 | 0,018610728 |
| ENSDARG00000042562 | allc | -1,229136026 | 2,355296482 | 53,73471801 | 4,96944E-06 | 0,000297051 |
| ENSDARG00000044875 | crygm2e | -1,233051563 | 0,307198402 | 17,09067826 | 0,00111758 | 0,009430845 |
| ENSDARG00000039351 | ccl19b | -1,234051571 | 3,252114785 | 15,35284813 | 0,002382214 | 0,015744341 |
| ENSDARG00000088915 | si:ch211-241n15.3 | -1,235677421 | -0,422120549 | 28,76330035 | 0,000118424 | 0,002260216 |
| ENSDARG00000017565 | itk | -1,236818694 | -1,127252759 | 14,61314867 | 0,002026106 | 0,014064791 |
| ENSDARG00000078878 | si:ch73-244f7.3 | -1,240884255 | 0,376652697 | 18,54500217 | 0,000807164 | 0,007632884 |
| ENSDARG00000087516 | CU693494.1 | -1,243962356 | -1,050579866 | 28,34939086 | 0,000126728 | 0,002376176 |
| ENSDARG00000062906 | kcnv2b | -1,245655286 | 1,97014849 | 14,74260119 | 0,002375023 | 0,015719509 |
| ENSDARG00000114879 | CABZ01072950.1 | -1,246181232 | -2,124567123 | 9,917864875 | 0,007479503 | 0,034158336 |
| ENSDARG00000024631 | nlrc5 | -1,246582683 | -1,430223035 | 12,27626293 | 0,003753391 | 0,021248794 |
| ENSDARG00000103760 | cfhl2 | -1,247249113 | 3,507136071 | 54,30586771 | 5,0852E-06 | 0,000300828 |
| ENSDARG00000045442 | cpb1 | -1,248140876 | 8,244747738 | 14,71248264 | 0,003158858 | 0,018940828 |
| ENSDARG00000104631 | CABZ01061592.2 | -1,248498033 | 0,278041103 | 12,248002 | 0,003882736 | 0,02175878 |
| ENSDARG00000087767 | CABZ01074298.1 | -1,248544498 | -0,692280356 | 20,60337921 | 0,000522745 | 0,005795508 |
| ENSDARG00000079946 | sqlea | -1,24900193 | 4,006106838 | 19,5052364 | 0,001052926 | 0,009077286 |
| ENSDARG00000109466 | si:dkeyp-77c4.5 | -1,249930533 | -1,384487159 | 16,29018302 | 0,001346461 | 0,010670082 |
| ENSDARG00000079980 | si:ch211-202h22.9 | -1,250052796 | 0,661066196 | 12,38535279 | 0,003858277 | 0,021687061 |
| ENSDARG00000035652 | sat1a.1 | -1,250068922 | 5,8788299 | 39,77245527 | 4,41676E-05 | 0,001252779 |
| ENSDARG00000100449 | BX322631.1 | -1,250430942 | 3,128451557 | 48,75156561 | 8,88618E-06 | 0,000426499 |
| ENSDARG00000113804 | si:dkey-205i10.1 | -1,256073965 | -0,343296388 | 24,02587902 | 0,000269095 | 0,003717429 |
| ENSDARG00000098736 | si:dkey-201i2.4 | -1,256809493 | -0,939895772 | 12,41299842 | 0,003614301 | 0,020707604 |
| ENSDARG00000093150 | fndc7rs3 | -1,257078843 | 1,522642568 | 37,48419751 | 3,2703E-05 | 0,001024748 |
| ENSDARG00000092916 | mlnl | -1,257481183 | 1,187873865 | 18,84675626 | 0,000787621 | 0,007521056 |
| ENSDARG00000105411 | si:ch211-113d11.5 | -1,26137557 | 6,213026428 | 43,56006954 | 2,72753E-05 | 0,000912925 |
| ENSDARG00000086752 | LO018205.1 | -1,261839527 | -0,682747323 | 21,42607243 | 0,000442848 | 0,005178019 |
| ENSDARG00000097639 | si:ch73-265h17.4 | -1,263484016 | -0,653417023 | 15,48774484 | 0,001631964 | 0,012221846 |
| ENSDARG00000060051 | slc47a2.1 | -1,26454699 | 2,486662353 | 33,36382501 | 6,2953E-05 | 0,001514613 |
| ENSDARG00000053684 | aldob | -1,2649697 | 10,13271301 | 98,98248932 | 1,5355E-07 | 3,73976E-05 |
| ENSDARG00000097611 | CU896691.2 | -1,265182869 | -1,975290374 | 8,524349165 | 0,011687094 | 0,046695347 |
| ENSDARG00000074666 | zgc:153760 | -1,265216801 | -0,001567787 | 24,30705691 | 0,000255541 | 0,003593581 |
| ENSDARG00000079544 | si:ch1073-464p5.5 | -1,26697634 | 6,01932057 | 78,55117526 | 7,57252E-07 | 9,68522E-05 |
| ENSDARG00000041030 | or111-1 | -1,26704051 | -0,95295928 | 8,285380118 | 0,012659219 | 0,049297527 |
| ENSDARG00000103659 | bco1l | -1,267950099 | 3,204014266 | 43,13343255 | 1,81715E-05 | 0,00069106 |
| ENSDARG00000056909 | LO018309.1 | -1,268362005 | 0,707224877 | 52,88357466 | 5,41831E-06 | 0,000311676 |
| ENSDARG00000103119 | si:dkeyp-33c10.7 | -1,268384455 | -0,330713328 | 15,38784014 | 0,001672186 | 0,01240941 |
| ENSDARG00000075745 | ovgp1 | -1,270666545 | -1,431444082 | 20,5929698 | 0,000523857 | 0,005796613 |
| ENSDARG00000013240 | zgc:172271 | -1,270786291 | 2,437964397 | 62,96571197 | 2,07777E-06 | 0,000182255 |
| ENSDARG00000055539 | epdl2 | -1,27151179 | 5,868384849 | 41,67446297 | 3,53898E-05 | 0,001073369 |
| ENSDARG00000107280 | FO744833.1 | -1,27232055 | -1,264377184 | 16,54388111 | 0,001268519 | 0,010276531 |
| ENSDARG00000100451 | olfcs2 | -1,272354789 | -1,539145221 | 12,62214808 | 0,003412924 | 0,019896991 |
| ENSDARG00000090428 | ctrb1 | -1,272387271 | 9,717153279 | 10,70886487 | 0,008369073 | 0,037024455 |
| ENSDARG00000052332 | ftr98 | -1,27379388 | -1,009180457 | 9,949677808 | 0,007406391 | 0,033939535 |
| ENSDARG00000099633 | sec14l7 | -1,278482761 | 4,307468959 | 100,6497154 | 1,3907E-07 | 3,46072E-05 |
| ENSDARG00000005335 | slc22a4 | -1,278738505 | 1,053637414 | 26,20725714 | 0,000182068 | 0,002908771 |

| | | | | | | |
|---|---|---|---|---|---|---|
| ENSDARG00000102316 | si:dkey-237m9.2 | -1,278892821 | -1,170766281 | 14,82392571 | 0,001921896 | 0,013626478 |
| ENSDARG00000010042 | dnm1a | -1,279541344 | 5,565736644 | 43,0376874 | 2,99783E-05 | 0,000973507 |
| ENSDARG00000073742 | prss59.2 | -1,280103888 | 9,539454046 | 24,20597694 | 0,000405529 | 0,004904481 |
| ENSDARG00000071429 | tdo2a | -1,28042042 | 5,040925846 | 41,04064291 | 3,74909E-05 | 0,001120518 |
| ENSDARG00000052027 | tspan1 | -1,281702787 | 0,203950742 | 27,70174257 | 0,000141105 | 0,002508249 |
| ENSDARG00000090930 | si:ch211-120g10.1 | -1,281818869 | 1,180151155 | 10,2884412 | 0,007613439 | 0,034648347 |
| ENSDARG00000004964 | cyp4t8 | -1,282015074 | 4,295454578 | 57,37226275 | 4,49916E-06 | 0,000280664 |
| ENSDARG00000098348 | znf326 | -1,282492603 | 1,738390355 | 109,271637 | 8,52017E-08 | 2,70498E-05 |
| ENSDARG00000105636 | BX323797.3 | -1,282775134 | 1,134156185 | 49,72863029 | 7,54229E-06 | 0,00038716 |
| ENSDARG00000034705 | pvalb7 | -1,283702469 | 6,914449649 | 50,03116087 | 1,23393E-05 | 0,00052902 |
| ENSDARG00000078455 | best4 | -1,285125879 | 0,558989782 | 28,51191334 | 0,00012339 | 0,002332689 |
| ENSDARG00000093545 | BX294165.1 | -1,286108319 | -1,46400216 | 11,67300344 | 0,004445931 | 0,023898802 |
| ENSDARG00000101769 | cysltr3 | -1,287226019 | -0,807059315 | 16,87225963 | 0,001175242 | 0,009773331 |
| ENSDARG00000006522 | otop2 | -1,287479246 | 1,09419131 | 13,96239855 | 0,002644368 | 0,016858862 |
| ENSDARG00000091800 | zgc:174260 | -1,288311329 | 3,085124446 | 124,2608951 | 3,92767E-08 | 1,52407E-05 |
| ENSDARG00000054898 | ms4a17a.16 | -1,288515634 | 0,936967585 | 19,42371949 | 0,000687566 | 0,006916141 |
| ENSDARG00000103324 | si:ch211-127b6.2 | -1,288766901 | 0,219242768 | 34,52819839 | 4,92152E-05 | 0,001319359 |
| ENSDARG00000102888 | gpr39 | -1,289659796 | 5,002677481 | 30,6538977 | 0,000171577 | 0,002817844 |
| ENSDARG00000075355 | ftr29 | -1,290360091 | -0,895851913 | 13,34720155 | 0,002808706 | 0,017559401 |
| ENSDARG00000088051 | AL935044.1 | -1,291161256 | -0,270055434 | 8,791670511 | 0,010988778 | 0,044716864 |
| ENSDARG00000043518 | si:dkey-239i20.2 | -1,291385733 | 1,958237072 | 14,17427753 | 0,00276302 | 0,017363869 |
| ENSDARG00000018587 | zgc:152658 | -1,291857681 | -1,856721154 | 8,371932305 | 0,012296634 | 0,048260569 |
| ENSDARG00000114880 | CU467856.3 | -1,292413302 | -1,202165142 | 9,35842288 | 0,00891284 | 0,038829792 |
| ENSDARG00000067992 | si:dkey-31e10.1 | -1,292706127 | -0,903943462 | 9,262707353 | 0,009191609 | 0,039651911 |
| ENSDARG00000100113 | AL935044.2 | -1,295864497 | -1,141846873 | 19,45880523 | 0,000663207 | 0,006751202 |
| ENSDARG00000042189 | tspan33b | -1,296579268 | 0,555271598 | 21,60297154 | 0,000427564 | 0,005056118 |
| ENSDARG00000044534 | CR631122.1 | -1,296610246 | -1,482925315 | 11,39340757 | 0,004816418 | 0,025163639 |
| ENSDARG00000092813 | fndc7rs4 | -1,300210202 | 3,905512176 | 36,5477427 | 5,65522E-05 | 0,001418599 |
| ENSDARG00000043131 | BX664625.1 | -1,303398142 | -2,121833815 | 10,49322267 | 0,006276762 | 0,030278166 |
| ENSDARG00000021339 | cpa5 | -1,303594816 | 9,715723837 | 11,03229057 | 0,007718045 | 0,03498949 |
| ENSDARG00000094284 | si:dkey-58f10.10 | -1,307591638 | -1,670943317 | 11,68807878 | 0,004426912 | 0,023858219 |
| ENSDARG00000104260 | si:ch211-117c19.1 | -1,307889529 | 1,125561248 | 39,9104984 | 2,38041E-05 | 0,000838644 |
| ENSDARG00000104008 | si:ch211-232d10.3 | -1,310021067 | 0,002401873 | 23,45923443 | 0,000299006 | 0,003982226 |
| ENSDARG00000075676 | fndc7b | -1,310031967 | 3,163375673 | 30,26643366 | 0,000127593 | 0,002382318 |
| ENSDARG00000089602 | si:dkey-217f16.1 | -1,31102605 | -1,369342388 | 19,19901249 | 0,000700879 | 0,007000838 |
| ENSDARG00000044935 | hpdb | -1,311879777 | 6,613410271 | 78,96874766 | 7,50704E-07 | 9,6554E-05 |
| ENSDARG00000067806 | zgc:158432 | -1,313945705 | 0,830436262 | 13,43214505 | 0,002999386 | 0,018337079 |
| ENSDARG00000051912 | hpxb | -1,320031721 | 2,649359106 | 19,00804836 | 0,00097853 | 0,008662983 |
| ENSDARG00000078474 | myo15ab | -1,321139108 | 0,360199908 | 23,95447885 | 0,000272667 | 0,003751923 |
| ENSDARG00000091499 | zmp:0000001020 | -1,321636507 | -1,771547729 | 12,77691149 | 0,003272249 | 0,019387773 |
| ENSDARG00000038742 | rbp1.2 | -1,322232424 | 2,801599434 | 55,46064496 | 4,18427E-06 | 0,00027126 |
| ENSDARG00000092604 | si:ch211-15j1.4 | -1,323358117 | 0,271761945 | 13,11810054 | 0,003078145 | 0,018633277 |
| ENSDARG00000060112 | sybu | -1,32490505 | 4,676433486 | 38,59158665 | 5,07264E-05 | 0,001342578 |
| ENSDARG00000095807 | hp | -1,325299732 | 4,450955538 | 83,14861312 | 4,71992E-07 | 7,83027E-05 |
| ENSDARG00000069029 | acss2l | -1,327734489 | 3,015003983 | 38,26896551 | 3,59016E-05 | 0,001081983 |
| ENSDARG00000097524 | BX005129.2 | -1,328194267 | -2,139978466 | 10,10839942 | 0,007053781 | 0,0327897 |
| ENSDARG00000021787 | abcb5 | -1,332674404 | 7,368717563 | 72,02700011 | 1,30224E-06 | 0,000137098 |
| ENSDARG00000038378 | sagb | -1,336729158 | 7,594259046 | 46,45327884 | 1,85488E-05 | 0,000699597 |
| ENSDARG00000098976 | CU457778.1 | -1,339130908 | -0,938435005 | 13,05013096 | 0,003039944 | 0,018485411 |
| ENSDARG00000053853 | slc13a2 | -1,341436198 | 5,848662925 | 28,81119483 | 0,000249775 | 0,003563376 |
| ENSDARG00000011463 | CABZ01044048.1 | -1,343230624 | 0,242139599 | 29,61498319 | 0,000103231 | 0,002069498 |
| ENSDARG00000101690 | si:ch73-158p21.2 | -1,344267203 | 1,63477599 | 42,22826894 | 1,78149E-05 | 0,000687735 |
| ENSDARG00000089383 | il1fma | -1,344704443 | -0,977217169 | 21,83213453 | 0,000408659 | 0,004921538 |
| ENSDARG00000099699 | BX649620.1 | -1,345152932 | -1,844978584 | 11,54581396 | 0,004610217 | 0,024494384 |
| ENSDARG00000095282 | BX950187.1 | -1,345387778 | -1,349701013 | 16,49786985 | 0,00128226 | 0,010343385 |
| ENSDARG00000102986 | csf1ra | -1,350451935 | 3,527989554 | 48,7944647 | 1,06134E-05 | 0,000474577 |
| ENSDARG00000075958 | si:dkey-266f7.9 | -1,35076782 | 2,765196659 | 26,07342063 | 0,000251081 | 0,003568127 |
| ENSDARG00000025914 | si:dkey-190g11.3 | -1,351763517 | -0,406225386 | 8,386524923 | 0,012616974 | 0,04918321 |
| ENSDARG00000090850 | serpina1l | -1,353204816 | 7,712241921 | 71,69844364 | 1,30329E-06 | 0,000137098 |
| ENSDARG00000077697 | si:dkey-105i14.1 | -1,357278984 | 1,70235444 | 44,95489389 | 1,28656E-05 | 0,000544002 |
| ENSDARG00000055395 | foxq1b | -1,359980769 | 3,922139754 | 195,0921699 | 2,42917E-09 | 2,64826E-06 |
| ENSDARG00000042993 | prss1 | -1,360767242 | 9,990433547 | 12,99540897 | 0,004706779 | 0,024795221 |
| ENSDARG00000086374 | isg15 | -1,361464813 | -1,029551895 | 13,47061052 | 0,002718672 | 0,017179486 |
| ENSDARG00000019532 | fads2 | -1,362081852 | 4,358423396 | 28,67484621 | 0,000222962 | 0,003303001 |
| ENSDARG00000036215 | lcp2b | -1,362446353 | -1,661345225 | 10,04511755 | 0,007191979 | 0,033256548 |
| ENSDARG00000037603 | zgc:162144 | -1,366342318 | 0,537598277 | 18,12134514 | 0,000904746 | 0,008222805 |
| ENSDARG00000095693 | CT573231.1 | -1,367414972 | -2,034876752 | 13,38520861 | 0,002780616 | 0,017426612 |
| ENSDARG00000094975 | si:ch211-214p16.2 | -1,36815155 | 2,341165167 | 37,77366812 | 3,38678E-05 | 0,001046384 |
| ENSDARG00000017314 | cela1.6 | -1,368346654 | 10,07482428 | 11,20518325 | 0,007430918 | 0,034024686 |
| ENSDARG00000106745 | FO904901.1 | -1,372894098 | 3,735240275 | 12,79073082 | 0,005016245 | 0,025850083 |
| ENSDARG00000104897 | si:dkey-16p6.1 | -1,376992065 | -0,540104839 | 35,09623029 | 4,54074E-05 | 0,001266208 |

| | | | | | | |
|---|---|---|---|---|---|---|
| ENSDARG00000042978 | cyp2p6 | -1,38285824 | 4,355165834 | 86,43286775 | 3,80519E-07 | 6,91397E-05 |
| ENSDARG00000105757 | or133-5 | -1,383618889 | -1,698338772 | 10,55944693 | 0,006153291 | 0,029858722 |
| ENSDARG00000079075 | si:ch211-71m22.3 | -1,383930136 | -1,418264389 | 21,83973029 | 0,000408049 | 0,004919583 |
| ENSDARG00000042613 | crp3 | -1,384463007 | 4,154074429 | 22,16232066 | 0,000689511 | 0,006922116 |
| ENSDARG00000078757 | si:ch211-212c13.8 | -1,385731098 | 3,854758056 | 87,73430277 | 3,12246E-07 | 6,10988E-05 |
| ENSDARG00000004459 | unc119.2 | -1,388997146 | 1,982005979 | 76,5385712 | 6,88446E-07 | 9,21224E-05 |
| ENSDARG00000095457 | si:ch211-147d7.5 | -1,392558375 | -0,632947196 | 23,770056 | 0,00028215 | 0,003831283 |
| ENSDARG00000099963 | si:dkey-27n6.1 | -1,393098548 | -0,908027866 | 17,84343879 | 0,000942434 | 0,008457891 |
| ENSDARG00000090537 | ifit11 | -1,395219569 | -0,129668885 | 13,33232587 | 0,002875588 | 0,017836282 |
| ENSDARG00000117454 | CR792457.1 | -1,39555238 | -0,58644595 | 32,70431537 | 6,4174E-05 | 0,001536819 |
| ENSDARG00000095911 | si:ch211-149b19.4 | -1,395945857 | 1,107891146 | 11,34806408 | 0,005696672 | 0,028217137 |
| ENSDARG00000075015 | soul5 | -1,396183817 | 5,521438326 | 34,86228682 | 0,000105538 | 0,002097386 |
| ENSDARG00000100738 | osmr | -1,396398107 | -1,46215163 | 23,70364987 | 0,000285657 | 0,003861165 |
| ENSDARG00000089179 | si:dkey-16p6.1 | -1,400070007 | -0,334578685 | 20,90145149 | 0,000492016 | 0,005555483 |
| ENSDARG00000014208 | zgc:55413 | -1,402617274 | -1,385239983 | 13,54495571 | 0,002666044 | 0,016935742 |
| ENSDARG00000077638 | zgc:171965 | -1,403566181 | -1,381682217 | 28,04466979 | 0,000133271 | 0,002436991 |
| ENSDARG00000079336 | strc1 | -1,403953679 | 1,489462953 | 77,51371756 | 6,40157E-07 | 8,82877E-05 |
| ENSDARG00000044748 | or115-12 | -1,404285703 | -1,393350458 | 13,51496712 | 0,00268713 | 0,017031878 |
| ENSDARG00000076900 | prozb | -1,407759612 | 1,904465186 | 32,71673702 | 6,83596E-05 | 0,001592091 |
| ENSDARG00000096913 | BX005129.1 | -1,411851967 | -0,368444362 | 25,10866257 | 0,000221017 | 0,003302842 |
| ENSDARG00000098861 | si:dkey-71b5.6 | -1,413563868 | -1,185231404 | 18,6863711 | 0,000782698 | 0,007500666 |
| ENSDARG00000075866 | cdcp2 | -1,416315093 | -0,317702708 | 25,26245633 | 0,000215025 | 0,003251504 |
| ENSDARG00000062442 | msrb1b | -1,416707735 | 0,073708508 | 24,03751936 | 0,000268517 | 0,003712218 |
| ENSDARG00000102805 | cyp2aa12 | -1,417334048 | 3,094849186 | 35,03479711 | 6,50721E-05 | 0,001545131 |
| ENSDARG00000100311 | si:dkey-165e24.1 | -1,417609478 | 3,202762325 | 56,20279571 | 4,37263E-06 | 0,000275777 |
| ENSDARG00000093303 | ifitm1 | -1,419247789 | 4,044938527 | 23,55366328 | 0,000538252 | 0,005888699 |
| ENSDARG00000055014 | si:dkey-33m11.8 | -1,419443491 | 2,716954216 | 78,79512366 | 5,82508E-07 | 8,41245E-05 |
| ENSDARG00000087753 | CR388166.1 | -1,419589221 | 0,723634359 | 28,75577623 | 0,000118569 | 0,002260216 |
| ENSDARG00000071306 | gip | -1,419732053 | 0,472401349 | 12,45212699 | 0,003904412 | 0,021839141 |
| ENSDARG00000019492 | shbg | -1,421958074 | 4,383314538 | 43,15015852 | 2,85841E-05 | 0,000938888 |
| ENSDARG00000099684 | trim35-24 | -1,424781147 | -0,95217526 | 18,7090428 | 0,000778855 | 0,007478137 |
| ENSDARG00000054741 | prkg2 | -1,428549733 | 1,481980801 | 34,70375336 | 4,80004E-05 | 0,001299995 |
| ENSDARG00000043997 | cyp2x8 | -1,431200024 | 1,656851509 | 10,63938099 | 0,007465017 | 0,034132946 |
| ENSDARG00000105346 | zp2.2 | -1,431509264 | -1,970668806 | 10,01037795 | 0,007269179 | 0,033530326 |
| ENSDARG00000056330 | mhc2dbb | -1,43703231 | -0,930002805 | 22,08224258 | 0,000389117 | 0,004746107 |
| ENSDARG00000029388 | si:dkey-90m5.4 | -1,439396531 | 3,091382689 | 53,32910521 | 5,92222E-06 | 0,000333804 |
| ENSDARG00000101030 | si:ch73-158p21.3 | -1,439402082 | 0,81684472 | 20,9581279 | 0,000507454 | 0,005672677 |
| ENSDARG00000105422 | si:dkey-1k23.3 | -1,440105927 | 0,528760326 | 8,897355471 | 0,011636043 | 0,046564511 |
| ENSDARG00000055876 | msmo1 | -1,443737371 | 5,267731771 | 25,98873373 | 0,000430602 | 0,00507893 |
| ENSDARG00000100360 | si:ch211-125c23.3 | -1,443899815 | -1,685544551 | 8,426013996 | 0,012076211 | 0,047708849 |
| ENSDARG00000096681 | BX927300.1 | -1,444395231 | -1,215966759 | 30,19894663 | 9,41086E-05 | 0,001943031 |
| ENSDARG00000019207 | abo | -1,444443583 | 0,135326433 | 15,04922643 | 0,001871836 | 0,013398717 |
| ENSDARG00000111625 | wu:fc75a09 | -1,446610866 | -0,065557192 | 24,20694063 | 0,000260274 | 0,003635585 |
| ENSDARG00000019025 | podn | -1,449937643 | 0,078815012 | 35,97422329 | 4,01693E-05 | 0,001170019 |
| ENSDARG00000010764 | flj13639 | -1,450658359 | 2,717080448 | 57,17363905 | 3,60514E-06 | 0,00025087 |
| ENSDARG00000060860 | pstpip1a | -1,450922044 | -0,389534736 | 20,85835499 | 0,000496327 | 0,005583745 |
| ENSDARG00000074824 | si:dkey-49c17.4 | -1,453081064 | -2,174628938 | 8,658643152 | 0,011179006 | 0,045241677 |
| ENSDARG00000108939 | FQ378016.1 | -1,455949015 | 1,79271788 | 29,56039356 | 0,000115266 | 0,002226911 |
| ENSDARG00000105298 | RF00001 | -1,456277119 | -1,58176691 | 12,07762964 | 0,00396663 | 0,02208464 |
| ENSDARG00000062182 | slc22a7b.3 | -1,458031015 | 1,441335401 | 22,8112582 | 0,000382196 | 0,004681646 |
| ENSDARG00000055520 | ftr61 | -1,459743241 | -2,196015636 | 10,3151179 | 0,006623338 | 0,031433398 |
| ENSDARG00000062132 | cyp4v8 | -1,467425195 | 5,068153606 | 81,69346613 | 7,90349E-07 | 9,9419E-05 |
| ENSDARG00000098189 | si:dkey-16p6.1 | -1,471350624 | 0,308793383 | 25,52625108 | 0,000205178 | 0,003161062 |
| ENSDARG00000093124 | scpp8 | -1,472844353 | -0,784692938 | 21,3267285 | 0,000451707 | 0,005256709 |
| ENSDARG00000092858 | si:ch1073-126c3.2 | -1,474547304 | 2,375822848 | 84,01588907 | 4,01863E-07 | 7,1877E-05 |
| ENSDARG00000038405 | zgc:153921 | -1,478421265 | -2,008057195 | 14,44731902 | 0,002112698 | 0,01446507 |
| ENSDARG00000117228 | CT025588.2 | -1,478541427 | -2,343544905 | 10,94804197 | 0,00548304 | 0,027447511 |
| ENSDARG00000089296 | si:dkey-240n22.6 | -1,484653603 | 3,189259923 | 17,59440574 | 0,001604371 | 0,012054634 |
| ENSDARG00000105495 | si:dkey-31g16.2 | -1,484709186 | -1,822767691 | 11,84235803 | 0,004237591 | 0,023171968 |
| ENSDARG00000006372 | ugt5c2 | -1,489253456 | 0,19896348 | 34,87779615 | 4,68301E-05 | 0,001283985 |
| ENSDARG00000001572 | prf1.3 | -1,493845869 | -1,794070603 | 10,16744542 | 0,006927606 | 0,032374077 |
| ENSDARG00000101071 | si:dkeyp-80d11.14 | -1,494010858 | 0,019667178 | 28,58094823 | 0,000122003 | 0,002314107 |
| ENSDARG00000013393 | guca1b | -1,496322579 | 2,426783189 | 49,91532894 | 7,93175E-06 | 0,00039562 |
| ENSDARG00000069607 | zgc:162331 | -1,497777807 | 3,000048537 | 14,36969785 | 0,003323067 | 0,019558632 |
| ENSDARG00000103006 | si:ch1073-303d10.1 | -1,506321781 | 2,521037981 | 56,58895467 | 3,81354E-06 | 0,000260127 |
| ENSDARG00000110290 | BX005380.1 | -1,512130081 | -1,256351864 | 21,20562449 | 0,000462785 | 0,005342909 |
| ENSDARG00000042112 | dio1 | -1,514325874 | 4,042095513 | 30,88894272 | 0,000166183 | 0,002779104 |
| ENSDARG00000101885 | si:dkey-20i20.11 | -1,514828309 | -0,073649278 | 13,1769116 | 0,003113392 | 0,018767244 |
| ENSDARG00000102525 | lck | -1,515360867 | -0,01280948 | 34,13516942 | 5,20653E-05 | 0,001364945 |
| ENSDARG00000099099 | CABZ01015815.1 | -1,517172823 | 3,39964832 | 14,57496316 | 0,003326034 | 0,019564803 |
| ENSDARG00000093193 | chia.6 | -1,518297729 | 3,280703614 | 41,21358734 | 3,10525E-05 | 0,000998479 |

| | | | | | | |
|---|---|---|---|---|---|---|
| ENSDARG00000105561 | BX072554.2 | -1,520508822 | -1,46031757 | 8,547887455 | 0,011625519 | 0,046536891 |
| 324818/100331324/569298/436641 | si:ch211-170d8.5 | -1,521503708 | 2,593905012 | 88,01028302 | 3,06568E-07 | 6,05049E-05 |
| ENSDARG00000088631 | BX000363.1 | -1,521966616 | -1,25984525 | 9,25959388 | 0,009236357 | 0,039799953 |
| ENSDARG00000027620 | bco2l | -1,522512461 | 3,017006076 | 27,82635281 | 0,000214672 | 0,003248314 |
| ENSDARG00000101621 | CABZ01040054.1 | -1,526756443 | 3,148087418 | 11,29590724 | 0,007301048 | 0,033591276 |
| ENSDARG00000056462 | crp2 | -1,526819408 | 0,123771892 | 13,24549561 | 0,003104833 | 0,018735844 |
| ENSDARG00000040004 | si:ch211-244a23.1 | -1,527228962 | 1,408551797 | 31,93047786 | 7,56408E-05 | 0,001697765 |
| ENSDARG00000053845 | cbln14 | -1,529286213 | 1,486938963 | 51,76239112 | 6,08267E-06 | 0,000338824 |
| ENSDARG00000095362 | si:ch211-77g15.32 | -1,530360045 | 1,807681903 | 16,86691471 | 0,00153158 | 0,01165304 |
| ENSDARG00000041033 | or119-2 | -1,531811332 | -0,996886139 | 13,67288368 | 0,002578218 | 0,016561649 |
| ENSDARG00000111123 | BX537263.1 | -1,53576122 | -0,379888678 | 16,40346942 | 0,001312869 | 0,010516735 |
| ENSDARG00000041645 | sb:cb37 | -1,536574226 | 4,956874413 | 87,64102412 | 5,18761E-07 | 8,14392E-05 |
| ENSDARG00000092066 | CT583708.1 | -1,538941524 | -2,196425497 | 12,61202511 | 0,003422367 | 0,019914315 |
| ENSDARG00000107066 | CABZ01074888.1 | -1,542866293 | -0,481670273 | 29,5952104 | 0,000103557 | 0,002072662 |
| ENSDARG00000103498 | epd | -1,543175218 | 6,722838317 | 52,45627653 | 1,28789E-05 | 0,000544002 |
| ENSDARG00000008788 | camk1gb | -1,545052125 | 5,30666493 | 67,4870839 | 3,08194E-06 | 0,000231011 |
| ENSDARG00000079274 | prss59.1 | -1,546735595 | 8,978100653 | 14,68555145 | 0,003391449 | 0,019812664 |
| ENSDARG00000055036 | itih3a | -1,546743523 | 4,061392245 | 69,76094833 | 1,7332E-06 | 0,000165333 |
| ENSDARG00000098011 | BX000438.2 | -1,547258037 | 3,257062548 | 29,13407002 | 0,000188236 | 0,002978219 |
| ENSDARG00000052688 | paqr5b | -1,547377951 | 2,618315809 | 68,65296062 | 1,27882E-06 | 0,00013681 |
| ENSDARG00000099583 | CABZ01034082.1 | -1,548741889 | 0,565342735 | 29,51919609 | 0,000104822 | 0,002084974 |
| ENSDARG00000034577 | si:dkey-23o4.6 | -1,549239551 | 1,446927364 | 13,12193549 | 0,003755546 | 0,02125049 |
| ENSDARG00000037836 | igfals | -1,549759537 | 1,073883791 | 35,04742817 | 4,57209E-05 | 0,001271483 |
| ENSDARG00000117300 | BX936456.1 | -1,55198805 | -0,274686601 | 32,79037336 | 6,336E-05 | 0,001520508 |
| ENSDARG00000074191 | zgc:172253 | -1,553806304 | 2,922303707 | 71,00460487 | 1,07837E-06 | 0,000123927 |
| ENSDARG00000026895 | tor1l3 | -1,554304798 | -0,915657478 | 21,58270803 | 0,000429283 | 0,005071214 |
| ENSDARG00000062057 | zgc:174945 | -1,557849357 | -1,964894378 | 11,86712112 | 0,004208084 | 0,023086479 |
| ENSDARG00000103077 | si:rp71-81e14.2 | -1,564405894 | -2,209863624 | 8,332912166 | 0,01245858 | 0,048731716 |
| ENSDARG00000078859 | g0s2 | -1,564520157 | 1,495293714 | 54,07346777 | 4,80281E-06 | 0,000292862 |
| ENSDARG00000103218 | BX927130.3 | -1,565669409 | 0,916350034 | 54,28876734 | 4,70024E-06 | 0,000289267 |
| ENSDARG00000092044 | si:dkey-22f5.9 | -1,567805798 | 3,285914131 | 11,87955427 | 0,006379169 | 0,030649462 |
| ENSDARG00000070960 | si:ch211-288g17.4 | -1,568983478 | 3,396649114 | 81,6274374 | 5,24686E-07 | 8,14392E-05 |
| ENSDARG00000098386 | ctps1b | -1,571032993 | 4,8438267 | 78,99737387 | 1,06621E-06 | 0,000123282 |
| ENSDARG00000091320 | nlrc6 | -1,576687004 | 0,636810093 | 76,322186 | 6,99722E-07 | 9,23717E-05 |
| ENSDARG00000116660 | pigrl2.3 | -1,57875463 | -2,184431954 | 13,70779905 | 0,002554831 | 0,016496082 |
| ENSDARG00000100792 | zgc:154142 | -1,579442114 | 4,222509077 | 12,69366133 | 0,005571933 | 0,027773534 |
| ENSDARG00000055240 | xdh | -1,579505357 | 4,773368492 | 79,45261743 | 1,00853E-06 | 0,000118078 |
| ENSDARG00000091211 | adh8a | -1,579884843 | 0,456295107 | 79,42324614 | 5,56444E-07 | 8,3263E-05 |
| ENSDARG00000112125 | FP236331.1 | -1,580885499 | -1,550584324 | 20,15350424 | 0,000573413 | 0,006128717 |
| ENSDARG00000030530 | slc22a2 | -1,582179703 | 4,086688996 | 33,51151376 | 0,000117771 | 0,00225816 |
| ENSDARG00000012903 | slc34a2a | -1,586559317 | 4,21287708 | 15,14198509 | 0,003148405 | 0,018898686 |
| ENSDARG00000094782 | si:ch211-196h16.4 | -1,588174007 | -1,035060786 | 25,2819534 | 0,000214279 | 0,003247978 |
| ENSDARG00000088911 | cbln17 | -1,588368572 | 0,428662036 | 33,37991189 | 5,8093E-05 | 0,001447204 |
| ENSDARG00000076448 | serpinf2a | -1,589767978 | 4,468612814 | 82,8090973 | 6,88968E-07 | 9,21224E-05 |
| ENSDARG00000012446 | zgc:113057 | -1,595175522 | -1,971648805 | 12,66935882 | 0,003369282 | 0,019727966 |
| ENSDARG00000062998 | pglyrp2 | -1,597949018 | 1,436807034 | 102,7566367 | 1,22962E-07 | 3,16303E-05 |
| ENSDARG00000076269 | zgc:172131 | -1,598970376 | 0,043020962 | 54,81471671 | 4,46011E-06 | 0,000279381 |
| ENSDARG00000033382 | grifin | -1,599694516 | 3,514305874 | 83,03683253 | 4,90761E-07 | 7,91232E-05 |
| ENSDARG00000094910 | si:dkey-22i16.7 | -1,607739372 | 3,196588134 | 25,60429259 | 0,000357904 | 0,004485725 |
| ENSDARG00000037619 | BX569798.1 | -1,607740692 | -1,20709956 | 27,23089353 | 0,000152742 | 0,002627248 |
| ENSDARG00000109654 | cmklrl2 | -1,611747986 | -1,15607122 | 17,02470641 | 0,001134645 | 0,009539686 |
| ENSDARG00000053460 | si:dkey-283b1.7 | -1,614746864 | 0,464200878 | 13,21596308 | 0,003343608 | 0,019635812 |
| ENSDARG00000092725 | si:dkey-188i13.8 | -1,615765109 | -0,555554591 | 15,83978552 | 0,001498872 | 0,011446198 |
| ENSDARG00000076844 | plin6 | -1,61908357 | 1,981458099 | 53,4986635 | 5,11136E-06 | 0,000301596 |
| ENSDARG00000041108 | ctsh | -1,623046826 | 4,78107274 | 137,0665319 | 2,47316E-08 | 1,1796E-05 |
| ENSDARG00000099377 | si:cabz01030277.1 | -1,624127741 | -0,917812585 | 22,12032384 | 0,000386237 | 0,004716005 |
| ENSDARG00000100406 | zgc:112265 | -1,628062053 | 6,946951464 | 134,4086485 | 2,90101E-08 | 1,33231E-05 |
| ENSDARG00000052122 | rag1 | -1,630178861 | 2,694167239 | 34,81835038 | 7,09483E-05 | 0,001627546 |
| ENSDARG00000042956 | cyp2ad6 | -1,630851523 | 4,321528069 | 25,61200534 | 0,000451656 | 0,005256709 |
| ENSDARG00000001303 | psmb8a | -1,631038315 | -0,656544552 | 21,24232375 | 0,000459395 | 0,00531627 |
| ENSDARG00000109996 | CABZ01101996.1 | -1,631531016 | 2,965996561 | 45,02057207 | 1,8813E-05 | 0,000705105 |
| ENSDARG00000058053 | serping1 | -1,631913183 | 2,825672333 | 54,03827206 | 6,11674E-06 | 0,00033941 |
| ENSDARG00000100795 | timp4.3 | -1,632034597 | 2,853178656 | 12,63472196 | 0,005161025 | 0,02638011 |
| ENSDARG00000087870 | adgrf3a | -1,632301118 | -0,287287602 | 14,05528735 | 0,002425195 | 0,015929785 |
| ENSDARG00000024503 | c6ast3 | -1,635568889 | 2,591231826 | 13,57577419 | 0,004026769 | 0,022343398 |
| ENSDARG00000092091 | cyp2x12 | -1,638323278 | -0,187361646 | 24,34090094 | 0,000253963 | 0,003586816 |
| ENSDARG00000070941 | igfbp6a | -1,644737862 | -2,17289051 | 14,98873094 | 0,001844715 | 0,013276616 |
| ENSDARG00000096843 | zgc:193593 | -1,645344521 | 1,916349917 | 17,96712475 | 0,001286069 | 0,010363934 |
| ENSDARG00000043492 | irf1a | -1,645523808 | 0,454796654 | 49,67912848 | 7,58255E-06 | 0,000388247 |
| ENSDARG00000036628 | cd74b | -1,649739027 | 0,808670958 | 23,62930795 | 0,000319062 | 0,004166914 |
| ENSDARG00000101479 | BX908782.2 | -1,651891703 | 1,62102036 | 25,04166204 | 0,000285063 | 0,003861165 |

| | | | | | | |
|---|---|---|---|---|---|---|
| ENSDARG00000117231 | FO834828.1 | -1,653277685 | 2,904823175 | 48,78695776 | 1,16743E-05 | 0,00051006 |
| ENSDARG00000016391 | calcoco1b | -1,658592883 | 3,158840151 | 34,24460739 | 8,70599E-05 | 0,001842098 |
| ENSDARG00000071143 | si:dkey-208m12.2 | -1,661619508 | 0,853497395 | 20,96438599 | 0,000560689 | 0,0060492 |
| ENSDARG00000005616 | bfb | -1,663103488 | 3,105825527 | 122,3106428 | 4,32257E-08 | 1,62231E-05 |
| ENSDARG00000104773 | junbb | -1,666178151 | 4,569437471 | 13,18485947 | 0,00515773 | 0,026374986 |
| ENSDARG00000078728 | znf1068 | -1,668263175 | -1,713451243 | 20,52424058 | 0,000531272 | 0,005850528 |
| ENSDARG00000059826 | crtac1a | -1,670065805 | 3,990625726 | 55,75087947 | 7,961E-06 | 0,000396215 |
| ENSDARG00000075614 | apoc4 | -1,674329023 | -1,847239001 | 10,9301663 | 0,005511951 | 0,027567271 |
| ENSDARG00000042010 | pklr | -1,676609569 | 7,10438142 | 27,22711707 | 0,000380493 | 0,004673287 |
| ENSDARG00000099599 | pigrl2.3 | -1,677850217 | -2,169248741 | 20,1692199 | 0,00057155 | 0,0061231 |
| ENSDARG00000012435 | spic | -1,679044388 | -1,703904883 | 13,65258311 | 0,002591929 | 0,016645054 |
| ENSDARG00000059951 | pld6 | -1,679977687 | -2,154557033 | 16,78891418 | 0,001198144 | 0,009877677 |
| ENSDARG00000052099 | agxta | -1,682884557 | 5,168253869 | 133,0836524 | 3,71653E-08 | 1,49274E-05 |
| ENSDARG00000114577 | si:dkey-159n16.2 | -1,68393921 | 2,057201335 | 21,19073795 | 0,000673883 | 0,006817442 |
| ENSDARG00000116378 | ms4a17a.17 | -1,69109553 | 2,27920201 | 57,2617079 | 3,81954E-06 | 0,000260127 |
| ENSDARG00000101380 | cryabb | -1,69130114 | 0,223208794 | 62,61636725 | 2,14319E-06 | 0,000185316 |
| ENSDARG00000105061 | si:ch73-261i21.5 | -1,691503136 | 1,790983489 | 44,61855913 | 1,45617E-05 | 0,00060092 |
| ENSDARG00000057408 | CABZ01067232.1 | -1,694067511 | 4,129758668 | 55,08685634 | 8,77384E-06 | 0,000421992 |
| ENSDARG00000097470 | FP015862.2 | -1,696505726 | -1,552856163 | 10,471206 | 0,006339125 | 0,030521122 |
| ENSDARG00000040497 | BX004816.1 | -1,706350563 | -1,546624643 | 19,35091832 | 0,00067856 | 0,006849631 |
| ENSDARG00000077960 | si:ch211-186e20.7 | -1,707130076 | 2,34771619 | 66,38482262 | 1,57564E-06 | 0,000154157 |
| ENSDARG00000044862 | opn1lw1 | -1,713561965 | 2,490370765 | 65,63103026 | 1,79568E-06 | 0,000168055 |
| ENSDARG00000091254 | si:ch73-59p9.2 | -1,721673514 | -1,607343825 | 11,18304057 | 0,005118844 | 0,026241127 |
| ENSDARG00000099401 | ccl33.3 | -1,721728403 | 1,384089645 | 12,80314713 | 0,004306375 | 0,023418085 |
| ENSDARG00000101617 | BX530032.1 | -1,721772007 | -1,479655803 | 12,06370225 | 0,003982101 | 0,022143846 |
| ENSDARG00000103951 | tspan34 | -1,723517579 | 1,024915198 | 24,38078852 | 0,000297871 | 0,003978539 |
| ENSDARG00000037191 | ttr | -1,725432446 | 4,558910795 | 32,19228757 | 0,00018183 | 0,002908771 |
| ENSDARG00000094386 | adgrg4b | -1,729095124 | 0,434920409 | 34,15219685 | 5,1938E-05 | 0,001364945 |
| ENSDARG00000100504 | ppp1r27b | -1,731672887 | 0,821701078 | 69,83934231 | 1,16068E-06 | 0,000128994 |
| ENSDARG00000045834 | si:dkey-14d8.7 | -1,741131587 | 4,271826573 | 12,31805215 | 0,006297457 | 0,030339642 |
| ENSDARG00000054649 | rac1l | -1,741237526 | -0,806430434 | 28,97961969 | 0,000114334 | 0,00221265 |
| ENSDARG00000093621 | si:dkey-27b3.4 | -1,742264282 | -0,844707895 | 22,75903133 | 0,000341412 | 0,004364202 |
| ENSDARG00000100370 | CT955963.1 | -1,744993748 | -1,263838759 | 23,55079585 | 0,000293923 | 0,00394667 |
| ENSDARG00000092027 | si:dkey-20i20.9 | -1,745831627 | -0,205798197 | 19,15336665 | 0,000732973 | 0,007220605 |
| ENSDARG00000114099 | trim35-30 | -1,747976179 | -1,047022698 | 14,40007322 | 0,002140787 | 0,014591003 |
| ENSDARG00000103342 | wu:fa56d06 | -1,751527378 | 5,8823427 | 22,48538991 | 0,000874007 | 0,008039182 |
| ENSDARG00000095455 | si:dkey-20i20.10 | -1,752394852 | 0,452775281 | 17,26613535 | 0,001258085 | 0,010213687 |
| ENSDARG00000091961 | pth1b | -1,753025059 | 0,015188281 | 48,16709783 | 8,93994E-06 | 0,000427288 |
| ENSDARG00000117109 | CR626884.5 | -1,757547238 | 1,564672058 | 28,01444104 | 0,000173371 | 0,002840174 |
| ENSDARG00000062995 | si:dkey-14o18.2 | -1,763146625 | 2,292319242 | 49,2911689 | 9,87532E-06 | 0,000453987 |
| ENSDARG00000092532 | si:dkey-21e2.8 | -1,763158452 | 2,151873249 | 20,4288273 | 0,000833103 | 0,007773364 |
| ENSDARG00000011886 | pdca | -1,767420148 | 3,944816247 | 29,28143122 | 0,000255312 | 0,003593581 |
| ENSDARG00000037495 | rtn4rl2b | -1,772288337 | 2,790600361 | 57,73772817 | 4,55017E-06 | 0,000282308 |
| ENSDARG00000077911 | treh | -1,772572916 | -0,723969378 | 50,08910892 | 7,25646E-06 | 0,000381031 |
| ENSDARG00000053620 | ebi3 | -1,772691002 | 2,22002407 | 33,37146718 | 8,67912E-05 | 0,001839813 |
| ENSDARG00000091783 | si:dkeyp-71f10.5 | -1,774885689 | 0,863499188 | 56,16202386 | 3,90671E-06 | 0,000263836 |
| ENSDARG00000069744 | mos | -1,787679033 | -1,432392936 | 35,41550551 | 4,34165E-05 | 0,00124092 |
| ENSDARG00000102776 | cxcl19 | -1,787998454 | -0,463496829 | 17,50258837 | 0,001048787 | 0,009060729 |
| ENSDARG00000101757 | si:dkey-27n6.4 | -1,788963845 | 2,865816709 | 25,45437351 | 0,000380191 | 0,004672087 |
| ENSDARG00000001712 | si:ch211-247n2.1 | -1,789996896 | 1,176835238 | 89,40617942 | 2,79607E-07 | 5,61521E-05 |
| ENSDARG00000090416 | scpp1 | -1,791421473 | 2,676603243 | 106,6655952 | 9,84304E-08 | 2,80384E-05 |
| ENSDARG00000098911 | zgc:153293 | -1,792246167 | 1,61490124 | 72,00092997 | 9,76239E-07 | 0,00011633 |
| ENSDARG00000077872 | CR626907.1 | -1,792264623 | 4,943172117 | 236,7101505 | 7,18456E-10 | 1,26526E-06 |
| ENSDARG00000093750 | cd180 | -1,794560677 | -1,714004269 | 15,46569539 | 0,001640744 | 0,012255529 |
| ENSDARG00000076388 | zgc:194686 | -1,795011202 | 0,653907048 | 24,31975531 | 0,000284617 | 0,003858032 |
| ENSDARG00000038199 | cdab | -1,797954445 | 2,863253501 | 58,39266782 | 4,51296E-06 | 0,00028076 |
| ENSDARG00000038968 | ccr6b | -1,799136694 | -0,899445955 | 22,31987998 | 0,000371541 | 0,004592902 |
| ENSDARG00000089758 | si:dkey-11o15.7 | -1,79963159 | -2,162999628 | 11,63003167 | 0,004500666 | 0,024096876 |
| ENSDARG00000099185 | chia.2 | -1,800088535 | 8,889518395 | 10,93529702 | 0,008804426 | 0,038489313 |
| ENSDARG00000020548 | si:dkey-77f5.10 | -1,801542026 | -2,285059036 | 10,87931826 | 0,005595161 | 0,027858987 |
| ENSDARG00000096564 | CU915827.1 | -1,807884049 | 0,413057595 | 61,8773731 | 2,28955E-06 | 0,00019342 |
| ENSDARG00000091136 | zgc:174259 | -1,807930147 | 0,574764005 | 69,99785291 | 1,14587E-06 | 0,000128596 |
| ENSDARG00000045854 | fgf23 | -1,810355877 | -0,23308173 | 23,32774442 | 0,000306484 | 0,004051183 |
| ENSDARG00000102856 | si:ch73-59f11.3 | -1,811141203 | -0,148513878 | 30,8594223 | 8,48893E-05 | 0,001819571 |
| ENSDARG00000102894 | CABZ01057159.1 | -1,811369437 | -1,840534294 | 14,48325827 | 0,002093574 | 0,014371902 |
| ENSDARG00000094344 | si:dkey-222h21.8 | -1,811858263 | -1,989221367 | 10,42098274 | 0,006414737 | 0,030762249 |
| ENSDARG00000041848 | rh50 | -1,814030892 | 1,397863313 | 19,30101577 | 0,000931463 | 0,008398941 |
| ENSDARG00000026229 | prnpa | -1,815176092 | 4,245077032 | 162,318147 | 7,67388E-09 | 5,16723E-06 |
| ENSDARG00000094491 | si:ch211-202m22.1 | -1,817110974 | -1,32727324 | 20,58029663 | 0,000525216 | 0,005806028 |
| ENSDARG00000054641 | tent5ab | -1,822958367 | 4,966557389 | 212,3316555 | 1,42713E-09 | 1,81515E-06 |
| ENSDARG00000086620 | gckr | -1,826170271 | -2,073163096 | 10,5498071 | 0,006171088 | 0,029911663 |

| | | | | | | |
|---|---|---|---|---|---|---|
| ENSDARG00000010097 | f9a | -1,827417089 | -0,410988871 | 35,47807056 | 4,30382E-05 | 0,001233188 |
| ENSDARG00000096645 | si:ch211-131k2.2 | -1,832169128 | -0,418021839 | 23,70284956 | 0,0002857 | 0,003861165 |
| ENSDARG00000055550 | FO834888.1 | -1,837155474 | 0,010042923 | 50,71321576 | 6,79054E-06 | 0,000360702 |
| ENSDARG00000109749 | si:ch211-162i8.7 | -1,838800439 | 1,097060597 | 25,27605511 | 0,000268428 | 0,003712218 |
| ENSDARG00000105263 | ccl36.1 | -1,839908396 | -0,678593977 | 25,05402602 | 0,000223191 | 0,003303001 |
| ENSDARG00000105708 | or109-11 | -1,845376682 | 2,151252816 | 81,17317466 | 4,90623E-07 | 7,91232E-05 |
| ENSDARG00000097571 | si:dkey-114l24.2 | -1,847910312 | -1,267237158 | 29,36655818 | 0,000107417 | 0,002121887 |
| ENSDARG00000041540 | sult1st2 | -1,853234477 | 3,289379282 | 16,33261541 | 0,002418335 | 0,01590045 |
| ENSDARG00000075128 | or103-5 | -1,854538159 | -1,613615076 | 22,73576839 | 0,000342936 | 0,004376347 |
| ENSDARG00000039900 | si:ch73-168d20.1 | -1,868226226 | -1,199489529 | 21,62681046 | 0,000425551 | 0,005037519 |
| ENSDARG00000092252 | si:dkey-103k4.1 | -1,869361459 | -1,475301374 | 13,64095354 | 0,002599821 | 0,016681702 |
| ENSDARG00000052578 | c6ast4 | -1,869534101 | 7,424442783 | 9,725623725 | 0,012360055 | 0,048429079 |
| ENSDARG00000089388 | ftr35 | -1,875839292 | -1,484999038 | 24,37064333 | 0,000252586 | 0,003577783 |
| ENSDARG00000110748 | CABZ01056052.1 | -1,87720779 | -1,976429825 | 21,14038058 | 0,000468882 | 0,005396976 |
| ENSDARG00000095592 | si:dkey-11o15.5 | -1,880728278 | -1,168948658 | 17,79510047 | 0,000952679 | 0,008533112 |
| ENSDARG00000100251 | si:ch211-152c8.2 | -1,883793643 | -0,748206664 | 37,50380224 | 3,26172E-05 | 0,001024748 |
| ENSDARG00000077058 | cela1.5 | -1,885346766 | 3,510236326 | 12,80656475 | 0,005530934 | 0,027605233 |
| ENSDARG00000104518 | tcrg | -1,886357489 | -1,925019682 | 22,54068338 | 0,000356025 | 0,004475186 |
| ENSDARG00000098478 | FO704758.1 | -1,892591115 | 1,543143061 | 62,33287185 | 2,19803E-06 | 0,00018847 |
| ENSDARG00000100890 | si:zfos-1069f5.1 | -1,89385939 | -2,044848332 | 12,51307087 | 0,00351628 | 0,020308205 |
| ENSDARG00000089312 | si:ch73-106l15.4 | -1,89948106 | 1,266089469 | 24,38436474 | 0,00034231 | 0,004370798 |
| ENSDARG00000036096 | smad3a | -1,900052507 | 4,704049943 | 310,4925663 | 1,27526E-10 | 4,17081E-07 |
| ENSDARG00000007040 | tmigd1 | -1,901070729 | 1,155302002 | 24,29816871 | 0,000336026 | 0,004321901 |
| ENSDARG00000068387 | slc6a18 | -1,908773533 | 2,255979113 | 25,89812554 | 0,000322743 | 0,004198223 |
| ENSDARG00000029230 | pnp4b | -1,918420857 | 5,728298113 | 28,23532701 | 0,000375138 | 0,004624882 |
| ENSDARG00000039116 | nr5a5 | -1,921072848 | 2,742063957 | 31,95083829 | 0,000135384 | 0,002456626 |
| ENSDARG00000113877 | hoxc13b | -1,92824164 | 0,112807863 | 42,78119428 | 1,66553E-05 | 0,000657914 |
| ENSDARG00000023759 | zgc:73226 | -1,932583723 | 3,314652541 | 79,35858526 | 8,80417E-07 | 0,000108367 |
| ENSDARG00000040683 | MEP1B | -1,937790482 | 3,266070235 | 22,64099773 | 0,00073736 | 0,007245934 |
| ENSDARG00000116216 | znf1046 | -1,93809524 | -1,521514584 | 29,11573819 | 0,000111844 | 0,002182914 |
| ENSDARG00000038293 | zgc:103559 | -1,938562629 | 2,130056594 | 58,9610711 | 3,59403E-06 | 0,000250859 |
| ENSDARG00000103295 | cyp3a65 | -1,93910264 | 7,782228573 | 15,26760137 | 0,003371588 | 0,01973642 |
| ENSDARG00000017490 | cel.1 | -1,946168123 | 6,526538134 | 17,39727335 | 0,002272058 | 0,015200611 |
| ENSDARG00000038666 | igfbp1b | -1,947050185 | 2,577238104 | 48,84041532 | 1,34831E-05 | 0,00056535 |
| ENSDARG00000018361 | sult1st3 | -1,950510894 | 3,145532959 | 36,9603133 | 7,58204E-05 | 0,001698467 |
| ENSDARG00000056012 | eve1 | -1,957797935 | -0,633364865 | 38,96890764 | 2,68781E-05 | 0,000906256 |
| ENSDARG00000053036 | si:ch211-203b20.7 | -1,95960981 | -2,055328358 | 21,80952195 | 0,000410481 | 0,004927926 |
| ENSDARG00000022832 | bnip4 | -1,963521616 | 1,497817769 | 166,7276776 | 6,46992E-09 | 4,93741E-06 |
| ENSDARG00000111882 | si:ch211-194m7.4 | -1,96532768 | -1,356526641 | 18,58456366 | 0,00080023 | 0,007589253 |
| ENSDARG00000089310 | gc | -1,969468118 | 5,687631765 | 109,7993965 | 2,33138E-07 | 5,0833E-05 |
| ENSDARG00000086419 | si:ch211-282j17.13 | -1,969817926 | -0,002619459 | 26,64168749 | 0,000175793 | 0,002856361 |
| ENSDARG00000088263 | zgc:174356 | -1,972965282 | 1,776047839 | 28,88286011 | 0,000175936 | 0,002856645 |
| ENSDARG00000023176 | tdo2b | -1,974067953 | 2,881562393 | 13,383049 | 0,004614269 | 0,024501688 |
| ENSDARG00000038439 | fabp10a | -1,980207128 | 7,667040061 | 67,80309703 | 4,03548E-06 | 0,000265616 |
| ENSDARG00000117382 | BX323560.1 | -1,982128016 | -1,698831961 | 19,7491941 | 0,000623831 | 0,006465367 |
| ENSDARG00000102641 | CR925761.3 | -1,99511271 | 0,574941769 | 54,06782359 | 4,80553E-06 | 0,000292862 |
| ENSDARG00000041433 | si:dkey-7c18.24 | -2,000618408 | 3,616410908 | 98,11944046 | 2,68248E-07 | 5,43474E-05 |
| ENSDARG00000044276 | spp1 | -2,012318523 | 3,981575543 | 158,2679853 | 1,0323E-08 | 6,46386E-06 |
| ENSDARG00000104016 | si:dkey-151j17.4 | -2,01502874 | 0,984872912 | 67,20100227 | 1,44271E-06 | 0,000145321 |
| ENSDARG00000089050 | cuzd1.1 | -2,021098847 | 2,024304254 | 37,0149991 | 5,60145E-05 | 0,001413827 |
| ENSDARG00000057498 | habp2 | -2,02723411 | 4,131478941 | 47,88849744 | 2,75394E-05 | 0,000919076 |
| ENSDARG00000117001 | CABZ01086812.1 | -2,027289894 | 2,960367211 | 229,5313389 | 8,73029E-10 | 1,33247E-06 |
| ENSDARG00000100635 | chia.1 | -2,0290893 | 5,741875001 | 31,20047562 | 0,000257743 | 0,003617181 |
| ENSDARG00000089432 | si:ch211-212k18.15 | -2,038262392 | -1,9056418 | 17,0962923 | 0,001116142 | 0,009425655 |
| ENSDARG00000105052 | cfhl5 | -2,041478835 | 0,754637573 | 87,15688733 | 3,24521E-07 | 6,20313E-05 |
| ENSDARG00000094840 | si:dkey-21e2.13 | -2,044089785 | 2,890844716 | 21,46234627 | 0,000886665 | 0,008113229 |
| ENSDARG00000094077 | si:dkey-21e2.16 | -2,044521379 | 3,872583234 | 18,52554962 | 0,001737637 | 0,012732217 |
| ENSDARG00000039269 | arg2 | -2,045039309 | 5,903519279 | 19,64792312 | 0,001537568 | 0,011694714 |
| ENSDARG00000092052 | gstk4 | -2,047534561 | 0,087493871 | 67,54781507 | 1,40147E-06 | 0,00014388 |
| ENSDARG00000017193 | cnn1a | -2,056703275 | 1,292261307 | 54,61644533 | 4,89048E-06 | 0,000294485 |
| ENSDARG00000095553 | lenep | -2,061821311 | 1,012054204 | 34,09881929 | 6,61069E-05 | 0,001560259 |
| ENSDARG00000097889 | si:ch73-265h17.2 | -2,065881775 | -1,387337028 | 18,56348344 | 0,000803916 | 0,007617902 |
| ENSDARG00000095909 | BX539325.1 | -2,067977906 | -1,888975982 | 22,34380033 | 0,000369823 | 0,004584041 |
| ENSDARG00000055595 | clul1 | -2,071529033 | 4,302737673 | 33,47944723 | 0,000170643 | 0,002812603 |
| ENSDARG00000060023 | ncapg2 | -2,072300237 | 3,442949113 | 163,5194054 | 7,29806E-09 | 5,11017E-06 |
| ENSDARG00000070353 | hoxc13a | -2,072649789 | 1,507485142 | 74,46506571 | 8,05745E-07 | 0,000100254 |
| ENSDARG00000011869 | galnt8a.2 | -2,077668024 | -2,189014547 | 28,09191255 | 0,000132232 | 0,002431579 |
| ENSDARG00000092788 | si:dkey-21e2.15 | -2,087173753 | 4,046726039 | 22,79803796 | 0,00081409 | 0,007673021 |
| ENSDARG00000009153 | pla2g1b | -2,096332109 | 4,187550564 | 16,87809802 | 0,002467752 | 0,016128094 |
| ENSDARG00000071025 | dytn | -2,101480509 | 0,931323644 | 118,1608206 | 5,325E-08 | 1,90485E-05 |
| ENSDARG00000037533 | mep1b | -2,101891585 | 0,661582273 | 9,651270453 | 0,010611446 | 0,043678251 |

| | | | | | | |
|---|---|---|---|---|---|---|
| ENSDARG00000070966 | insl5a | -2,102211891 | 1,153712201 | 25,53023335 | 0,000291987 | 0,003925283 |
| ENSDARG00000043468 | sytl3 | -2,10247479 | -1,477301435 | 22,62007976 | 0,00035063 | 0,004442343 |
| ENSDARG00000044566 | fabp6 | -2,103365389 | 6,276449333 | 17,12406342 | 0,002514894 | 0,016333614 |
| ENSDARG00000042953 | cyp2n13 | -2,103433604 | 5,532685946 | 13,11257644 | 0,005570302 | 0,027771447 |
| ENSDARG00000088794 | si:ch73-180n10.1 | -2,119541539 | -0,972173333 | 11,72167547 | 0,004822817 | 0,025179835 |
| ENSDARG00000102945 | ccl39.2 | -2,123613207 | -1,709870261 | 15,24292111 | 0,001732584 | 0,012702198 |
| ENSDARG00000056015 | hoxb13a | -2,124124707 | 1,555344674 | 94,36366431 | 2,03646E-07 | 4,57084E-05 |
| ENSDARG00000007080 | rhcgl1 | -2,139313389 | 5,226778318 | 22,73344128 | 0,000933966 | 0,008408263 |
| ENSDARG00000030915 | cpa1 | -2,170931531 | 4,973689853 | 16,20311442 | 0,002999579 | 0,018337079 |
| ENSDARG00000102175 | hamp | -2,175214292 | 2,640085505 | 19,91332424 | 0,001192637 | 0,009857126 |
| ENSDARG00000040736 | crygm7 | -2,193948416 | -0,889460121 | 46,38446909 | 1,09143E-05 | 0,000485189 |
| ENSDARG00000094845 | zmp:0000001323 | -2,22439958 | -2,279215358 | 37,68302675 | 3,18443E-05 | 0,001009756 |
| ENSDARG00000093600 | omd | -2,227423607 | 0,809771476 | 126,2623762 | 3,5649E-08 | 1,48391E-05 |
| ENSDARG00000070964 | si:rp71-15k1.1 | -2,227844675 | -0,995016716 | 27,46945134 | 0,000146713 | 0,002573836 |
| ENSDARG00000041685 | si:dkey-105h12.2 | -2,230187324 | 4,427641276 | 69,49186947 | 4,32251E-06 | 0,000275431 |
| ENSDARG00000104899 | CR931813.2 | -2,232023239 | 1,745767195 | 12,10473002 | 0,006022838 | 0,029387492 |
| ENSDARG00000092569 | BX571681.1 | -2,240442087 | -2,511623445 | 24,89531399 | 0,00022965 | 0,003359493 |
| ENSDARG00000100942 | zgc:194655 | -2,245050587 | -2,125218282 | 49,87875631 | 7,42171E-06 | 0,000385289 |
| ENSDARG00000102219 | si:ch211-12p12.2 | -2,245632238 | -1,677564069 | 25,56319606 | 0,00020384 | 0,003146804 |
| ENSDARG00000096445 | si:ch211-214p16.3 | -2,260253853 | 4,433030308 | 10,8066128 | 0,0094574 | 0,040425266 |
| ENSDARG00000091650 | igflr1 | -2,275724823 | 0,741469456 | 22,92381625 | 0,000466546 | 0,005380906 |
| ENSDARG00000006848 | parp9 | -2,280895892 | 1,440172275 | 124,7893579 | 3,82792E-08 | 1,51097E-05 |
| ENSDARG00000074712 | si:ch211-186e20.2 | -2,281872858 | 0,227456542 | 36,52066093 | 4,0027E-05 | 0,001168849 |
| ENSDARG00000088641 | grn2 | -2,298500649 | 0,261661773 | 18,51169551 | 0,001053829 | 0,009077286 |
| ENSDARG00000031745 | si:busm1-266f07.2 | -2,307919347 | -0,480365228 | 22,24510212 | 0,000408071 | 0,004919583 |
| ENSDARG00000045153 | slc16a8 | -2,313766673 | 2,186566665 | 33,66360031 | 0,000111156 | 0,002173188 |
| ENSDARG00000105807 | CU672227.1 | -2,315831412 | -2,277184834 | 16,15202542 | 0,001391224 | 0,010889125 |
| ENSDARG00000104947 | si:ch211-194m7.8 | -2,320832091 | -1,976059952 | 28,10845248 | 0,00013187 | 0,002428829 |
| ENSDARG00000116986 | zgc:172315 | -2,321283795 | 0,659177162 | 84,04896244 | 4,00945E-07 | 7,1877E-05 |
| ENSDARG00000079589 | si:dkeyp-73d8.6 | -2,326866442 | -2,108518057 | 26,23231836 | 0,000181277 | 0,002904234 |
| ENSDARG00000042980 | cyp2p7 | -2,339089829 | 3,324402547 | 26,38765742 | 0,000450743 | 0,005251561 |
| ENSDARG00000117264 | CR385050.2 | -2,367761516 | 2,301071516 | 31,24400964 | 0,00017364 | 0,002840174 |
| ENSDARG00000053802 | cbln5 | -2,391682089 | 0,311890866 | 72,68750402 | 9,24885E-07 | 0,000111535 |
| ENSDARG00000068290 | cyp2x12 | -2,403196097 | 0,848558044 | 82,64830891 | 4,42028E-07 | 7,61169E-05 |
| ENSDARG00000101087 | si:dkeyp-80d11.1 | -2,405874779 | -0,316423377 | 80,14663407 | 5,28074E-07 | 8,14392E-05 |
| ENSDARG00000003523 | itln3 | -2,409234414 | 3,667994513 | 32,39769236 | 0,000205886 | 0,003167713 |
| ENSDARG00000091970 | ms4a17a.3 | -2,417803183 | -0,590258217 | 61,73394736 | 2,31927E-06 | 0,000194057 |
| ENSDARG00000094959 | si:ch73-379f5.5 | -2,425577968 | -1,030057551 | 28,90475418 | 0,000115731 | 0,002232132 |
| ENSDARG00000093780 | BX548078.1 | -2,439703288 | 2,432152765 | 50,60361123 | 1,41772E-05 | 0,000586933 |
| ENSDARG00000023151 | ucp1 | -2,441441572 | 8,161992935 | 41,19792251 | 7,89483E-05 | 0,001749702 |
| ENSDARG00000115234 | crygm2b | -2,443424729 | -0,39501175 | 47,82217176 | 9,28755E-06 | 0,000435796 |
| ENSDARG00000069630 | tat | -2,453237055 | 4,806170223 | 49,59401935 | 3,5918E-05 | 0,001081983 |
| ENSDARG00000092510 | CR405715.1 | -2,470202134 | -1,805855346 | 14,92844601 | 0,001920505 | 0,013620829 |
| ENSDARG00000030254 | cbln9 | -2,485362133 | 1,711097132 | 108,0460706 | 9,115E-08 | 2,7664E-05 |
| ENSDARG00000074457 | si:dkey-96f10.1 | -2,486908349 | 1,475760858 | 43,9515651 | 2,34453E-05 | 0,000832182 |
| ENSDARG00000034095 | si:dkey-59l11.10 | -2,498879799 | -1,146432563 | 39,92776841 | 2,37516E-05 | 0,000838644 |
| ENSDARG00000105371 | CR382283.1 | -2,524738621 | -1,930660497 | 16,91528356 | 0,001170302 | 0,009736902 |
| ENSDARG00000069293 | ahsg2 | -2,52526773 | 3,028065661 | 26,85147587 | 0,000421385 | 0,005011526 |
| ENSDARG00000079221 | si:ch211-162k9.6 | -2,532369115 | -2,183989093 | 33,1418169 | 6,01571E-05 | 0,00148409 |
| ENSDARG00000103187 | syce2 | -2,539174183 | -1,92830232 | 34,49897296 | 4,94208E-05 | 0,001323322 |
| ENSDARG00000099456 | BX293991.1 | -2,554564182 | -1,143378703 | 47,53970081 | 9,58383E-06 | 0,000444154 |
| ENSDARG00000117741 | LO018107.1 | -2,561309129 | -0,301421967 | 39,03310735 | 2,76264E-05 | 0,00092064 |
| ENSDARG00000026764 | ahsg1 | -2,574932167 | 3,596291503 | 119,5978261 | 1,25236E-07 | 3,18573E-05 |
| ENSDARG00000105408 | BX248082.1 | -2,575703477 | 1,503505174 | 83,2525366 | 4,82517E-07 | 7,89053E-05 |
| ENSDARG00000007534 | itln1 | -2,600612383 | -0,131043845 | 80,86474495 | 5,01545E-07 | 8,02278E-05 |
| ENSDARG00000032465 | slc1a8b | -2,610453302 | 2,840750656 | 14,28828921 | 0,00411275 | 0,022693976 |
| ENSDARG00000025311 | cuzd1.2 | -2,620527506 | 2,683032302 | 73,52270523 | 1,9777E-06 | 0,000179673 |
| ENSDARG00000097247 | BX322555.3 | -2,622409584 | -0,976996786 | 15,93759904 | 0,001708349 | 0,012585377 |
| ENSDARG00000094002 | ccl34b.4 | -2,635743691 | -1,474184135 | 36,23620636 | 3,87435E-05 | 0,001143033 |
| ENSDARG00000068951 | si:ch211-219a15.4 | -2,641312649 | 2,170565214 | 19,87257981 | 0,001243744 | 0,010136804 |
| ENSDARG00000101051 | ctsbb | -2,651881644 | 4,075136241 | 13,65103475 | 0,00496864 | 0,025695065 |
| ENSDARG00000030778 | BX323854.1 | -2,660240741 | -1,82671829 | 29,42907991 | 0,000106346 | 0,002107946 |
| ENSDARG00000052731 | ankrd31 | -2,661762354 | -1,515029531 | 17,19161913 | 0,001158199 | 0,009680838 |
| ENSDARG00000100716 | si:dkey-285b23.4 | -2,662989996 | -1,282031552 | 36,98289326 | 3,49889E-05 | 0,001067062 |
| ENSDARG00000094643 | ms4a17c.1 | -2,663780845 | -2,054951366 | 52,72077227 | 5,50939E-06 | 0,00031533 |
| ENSDARG00000094923 | CU469568.2 | -2,711760703 | -0,321625601 | 168,3860711 | 6,08452E-09 | 4,80341E-06 |
| ENSDARG00000076534 | si:ch211-14a17.10 | -2,712532499 | 2,320843065 | 54,66237408 | 1,05249E-05 | 0,000473393 |
| ENSDARG00000076257 | si:ch211-285c6.1 | -2,714654165 | -1,12892856 | 31,84060657 | 7,30463E-05 | 0,001664002 |
| ENSDARG00000097608 | CU929312.1 | -2,717503766 | -0,847397993 | 67,69340116 | 1,38457E-06 | 0,000143539 |
| ENSDARG00000100540 | CABZ01114105.1 | -2,723969304 | 1,658225253 | 173,4618199 | 5,05924E-09 | 4,37783E-06 |
| ENSDARG00000096739 | si:dkey-219e21.2 | -2,724399295 | 2,277929259 | 87,82794899 | 5,60864E-07 | 8,33794E-05 |

| | | | | | | |
|---|---|---|---|---|---|---|
| ENSDARG00000060622 | si:ch73-14h1.2 | -2,725996924 | 2,587694695 | 89,42020069 | 5,73056E-07 | 8,41245E-05 |
| ENSDARG00000112540 | cst14b.2 | -2,72811296 | 0,131574469 | 74,14070039 | 8,26103E-07 | 0,000102231 |
| ENSDARG00000078349 | zmp:0000000845 | -2,741190079 | 0,536060758 | 48,19814456 | 1,10853E-05 | 0,000490884 |
| ENSDARG00000086418 | si:ch211-236p5.3 | -2,749129653 | -0,362011344 | 23,44835174 | 0,000375743 | 0,004629419 |
| ENSDARG00000051914 | slc14a2 | -2,75948995 | 2,728409888 | 204,7384335 | 1,79449E-09 | 2,14484E-06 |
| ENSDARG00000101749 | zgc:92161 | -2,796487275 | 3,466146629 | 41,29665162 | 7,54094E-05 | 0,001695898 |
| ENSDARG00000103773 | CR457444.5 | -2,80144589 | -1,724922049 | 15,22799447 | 0,001871137 | 0,013398717 |
| ENSDARG00000090769 | si:dkey-51d8.6 | -2,806369849 | -0,978836569 | 47,22621266 | 9,92546E-06 | 0,000455378 |
| ENSDARG00000098995 | cyp2k6 | -2,84579063 | 3,040245466 | 34,98182991 | 0,000147559 | 0,002582734 |
| ENSDARG00000093198 | c3a.4 | -2,84642824 | 0,675023023 | 70,96594114 | 1,1606E-06 | 0,000128994 |
| ENSDARG00000003902 | ctsl.1 | -2,849345157 | 7,06443232 | 34,83619245 | 0,000202193 | 0,003129817 |
| ENSDARG00000097728 | CR388132.1 | -2,860304601 | 0,473503902 | 15,70631665 | 0,002344986 | 0,015556681 |
| ENSDARG00000097397 | ugt5a4 | -2,879274718 | -1,204088207 | 34,23524035 | 5,1322E-05 | 0,001356774 |
| ENSDARG00000070581 | ggact.1 | -2,886206677 | -0,115206563 | 139,695487 | 1,92484E-08 | 1,02482E-05 |
| ENSDARG00000053854 | uts2a | -2,892467482 | -1,267758514 | 19,07488912 | 0,000798155 | 0,007579094 |
| ENSDARG00000097533 | BX571880.1 | -2,948002873 | 2,729886104 | 123,0039544 | 8,28455E-08 | 2,70498E-05 |
| ENSDARG00000045089 | si:ch211-270n8.1 | -2,992976291 | 4,104976568 | 101,8594394 | 6,7627E-07 | 9,16126E-05 |
| ENSDARG00000100640 | BX321870.2 | -3,012859978 | -1,961050187 | 33,01564975 | 6,1285E-05 | 0,001499917 |
| ENSDARG00000103806 | BX004785.3 | -3,046322343 | 0,672153804 | 37,68375155 | 5,27955E-05 | 0,001373886 |
| ENSDARG00000079901 | si:ch73-15n24.1 | -3,097420254 | 1,580883526 | 112,1870632 | 8,52079E-08 | 2,70498E-05 |
| ENSDARG00000075932 | si:ch73-211l2.3 | -3,152428672 | -2,191579591 | 40,87024368 | 2,10796E-05 | 0,000768466 |
| ENSDARG00000021172 | cyp2ad2 | -3,204325069 | 5,76454967 | 41,02253542 | 0,000117049 | 0,002248083 |
| ENSDARG00000087597 | si:dkey-51d8.3 | -3,20736909 | 1,281521784 | 279,8433678 | 2,47876E-10 | 6,3054E-07 |
| ENSDARG00000053476 | lipca | -3,216735684 | 2,366415621 | 225,0727329 | 9,88241E-10 | 1,41405E-06 |
| ENSDARG00000070892 | si:ch211-262h13.5 | -3,227083791 | 1,921311011 | 95,39534954 | 3,56001E-07 | 6,57281E-05 |
| ENSDARG00000087143 | serpina7 | -3,262129011 | 2,124542951 | 78,34995384 | 1,46631E-06 | 0,000145955 |
| ENSDARG00000096586 | MFAP4 (1 of many) | -3,262897556 | -1,495005979 | 22,96609767 | 0,000353992 | 0,004469428 |
| ENSDARG00000102617 | timp4.2 | -3,281353051 | 0,425688998 | 40,10051353 | 3,70762E-05 | 0,001112481 |
| ENSDARG00000102704 | si:ch1073-376c22.1 | -3,352587589 | 0,561831928 | 32,82913203 | 0,000112743 | 0,002194845 |
| ENSDARG00000103833 | spink2.2 | -3,371752121 | -0,152251779 | 23,41661565 | 0,000454861 | 0,0052727 |
| ENSDARG00000095200 | si:ch211-197e7.1 | -3,383263955 | 0,546362919 | 30,07522823 | 0,000176698 | 0,002862934 |
| ENSDARG00000102722 | itih3b | -3,455566703 | 4,624967313 | 320,7507989 | 3,17436E-10 | 6,6067E-07 |
| ENSDARG00000117628 | BX005011.1 | -3,464976838 | -1,935501475 | 34,7028193 | 4,80068E-05 | 0,001299995 |
| ENSDARG00000007371 | slc26a3.1 | -3,484282471 | -1,249906806 | 44,97654944 | 1,28331E-05 | 0,000544002 |
| ENSDARG00000095649 | si:dkey-97a13.12 | -3,508329811 | 2,28708209 | 41,6771427 | 6,54905E-05 | 0,001550506 |
| ENSDARG00000025046 | AL935186.1 | -3,529666068 | 1,78660637 | 32,19781625 | 0,000191332 | 0,003010561 |
| ENSDARG00000114345 | si:dkey-28d5.5 | -3,569461683 | -1,851179547 | 63,25970009 | 2,02449E-06 | 0,00018131 |
| ENSDARG00000099811 | si:dkey-203a12.7 | -3,605047137 | -0,313403067 | 44,48097642 | 1,70268E-05 | 0,000665206 |
| ENSDARG00000016290 | clca1 | -3,63125762 | 2,076422702 | 35,85097244 | 0,000130258 | 0,002410312 |
| ENSDARG00000097553 | BX908750.1 | -3,745269476 | 1,247518042 | 109,4285035 | 1,22344E-07 | 3,16303E-05 |
| ENSDARG00000095005 | si:dkey-21e2.10 | -3,790097908 | -1,33082902 | 55,81188059 | 4,04254E-06 | 0,000265616 |
| ENSDARG00000117613 | CABZ01030033.1 | -3,868515369 | -0,203317303 | 75,31152499 | 7,71395E-07 | 9,80041E-05 |
| ENSDARG00000056498 | crp2 | -3,91664426 | 6,154505356 | 50,26483626 | 5,6573E-05 | 0,001418599 |
| ENSDARG00000117543 | CU682640.1 | -3,971946974 | -1,735409185 | 79,96709698 | 5,34957E-07 | 8,14392E-05 |
| ENSDARG00000113858 | AL928650.4 | -4,111194806 | -0,754600306 | 31,3422343 | 0,00011152 | 0,002178449 |
| ENSDARG00000096577 | adgre9 | -4,135877044 | -0,073414796 | 116,8486983 | 5,69586E-08 | 1,97577E-05 |
| ENSDARG00000096217 | si:dkey-51d8.9 | -4,150109371 | -1,156022243 | 70,90523897 | 1,06515E-06 | 0,000123282 |
| ENSDARG00000096107 | BX649516.4 | -4,345862385 | -1,164784338 | 91,21991015 | 2,48551E-07 | 5,31807E-05 |
| ENSDARG00000101290 | spink2.5 | -4,39190808 | -0,054913621 | 55,48319278 | 6,45188E-06 | 0,000348371 |
| ENSDARG00000079043 | si:dkeyp-75b4.10 | -4,408701528 | 0,295690318 | 81,7537203 | 6,47997E-07 | 8,88338E-05 |
| ENSDARG00000102271 | si:dkey-203a12.2 | -4,534627548 | -1,627621343 | 57,6559493 | 3,38322E-06 | 0,000243196 |
| ENSDARG00000100941 | clca1 | -4,658850504 | -0,450726772 | 35,76154315 | 6,64702E-05 | 0,001561251 |
| ENSDARG00000097453 | si:ch211-225b7.5 | -4,675767742 | -0,785348139 | 107,2595587 | 9,52189E-08 | 2,80384E-05 |
| ENSDARG00000100051 | spink2.4 | -4,712951419 | -1,101862363 | 38,6833291 | 3,48294E-05 | 0,001066022 |
| ENSDARG00000092358 | BX469930.1 | -4,738968356 | 3,560543682 | 181,2167706 | 3,18624E-08 | 1,4028E-05 |
| ENSDARG00000096618 | BX000534.1 | -4,945377184 | -1,736061071 | 64,75039092 | 1,77738E-06 | 0,000168055 |
| ENSDARG00000103538 | BX571954.2 | -5,026883202 | -1,627244951 | 57,18997988 | 3,53731E-06 | 0,000247969 |
| ENSDARG00000109460 | CR762420.1 | -5,30033769 | -1,408913511 | 41,82118352 | 2,22513E-05 | 0,000798467 |
| ENSDARG00000096587 | itgam | -5,331606247 | -1,25845987 | 97,58825853 | 1,67001E-07 | 3,91591E-05 |
| ENSDARG00000104004 | si:dkey-146c18.2 | -5,339642266 | -1,132346125 | 29,89509061 | 0,000148073 | 0,002586273 |
| ENSDARG00000111606 | smyhc2 | -5,372358057 | 0,438246244 | 172,8965792 | 5,16299E-09 | 4,37783E-06 |
| ENSDARG00000093173 | si:dkey-93m18.6 | -5,430101672 | -0,534923161 | 139,9862355 | 1,9005E-08 | 1,02482E-05 |
| ENSDARG00000040738 | zgc:153846 | -5,435937697 | -1,578981112 | 72,94376875 | 9,06519E-07 | 0,000110393 |
| ENSDARG00000012609 | hpxa | -5,663299756 | 4,312719191 | 136,6547347 | 4,11479E-07 | 7,24646E-05 |
| ENSDARG00000098581 | CR457444.2 | -5,729666742 | 0,717370828 | 43,18524288 | 4,88212E-05 | 0,001316505 |
| ENSDARG00000019294 | cbln8 | -5,772995958 | 1,079799362 | 165,7561717 | 1,04116E-08 | 6,46386E-06 |
| ENSDARG00000105265 | zgc:171534 | -5,932464021 | 4,040221495 | 21,05434185 | 0,001451701 | 0,011201623 |
| ENSDARG00000117107 | CABZ01058647.1 | -5,993960663 | -0,286017119 | 136,7619399 | 2,19164E-08 | 1,14035E-05 |
| ENSDARG00000088274 | si:ch211-181d7.1 | -6,242196554 | -1,399171969 | 181,8199948 | 3,77375E-09 | 3,59984E-06 |
| ENSDARG00000104642 | si:dkey-31i7.2 | -6,509604778 | 0,016943183 | 191,5288596 | 2,72642E-09 | 2,75989E-06 |
| ENSDARG00000036833 | upp2 | -6,798264591 | 1,416293103 | 331,8285197 | 8,32559E-11 | 3,24968E-07 |

| | | | | | | |
|---|---|---|---|---|---|---|
| ENSDARG00000096579 | si:dkey-9c18.3 | -7,04119486 | -0,897300836 | 105,634964 | 1,04303E-07 | 2,87701E-05 |
| ENSDARG00000113739 | CABZ01055489.1 | -7,652454561 | -2,178866217 | 91,09374023 | 4,06415E-06 | 0,000265616 |
| ENSDARG00000057844 | zgc:171470 | -8,823746827 | 0,158760258 | 262,3498277 | 3,74003E-10 | 7,13536E-07 |
| ENSDARG00000110614 | ugt2a7 | -8,986670093 | -1,008578928 | 116,8618182 | 1,38872E-06 | 0,000143539 |
| ENSDARG00000096599 | AL928650.3 | -9,04545834 | 0,298889999 | 274,3795078 | 2,81096E-10 | 6,43542E-07 |
| ENSDARG00000068621 | si:ch211-181d7.3 | -9,693677019 | 2,38292667 | 918,9864403 | 1,09566E-13 | 1,2542E-09 |
| ENSDARG00000091847 | si:ch211-181d7.1 | -11,47728446 | 2,756632696 | 1124,356203 | 2,90384E-14 | 6,64806E-10 |

**Supplementary Table 5:** Gene Ontology term enrichment for upregulated genes in 5 dpf qKO zebrafish versus WT controls (p.adjust < 0.05). Table includes: GeneRatio (ratio of input genes that are annotated in a term) and BgRatio (ratio of all genes that are annotated in this term), p-value (raw p-value), p.adjust (Bonferroni adjusted p-values), geneID (identity of the DEG included in the GO category), Enrichment (the log2 of the ratio of the gene set (%) in the DEG and the gene set (%) in the universe).

| ID | Description | Gene Ratio | BgRatio | pvalue | p.adjust | geneID | log2 Enrichment |
|---|---|---|---|---|---|---|---|
| *GO:0044550* | secondary metabolite biosynthetic process | 6/146 | 10/20304 | 2,56E-11 | 2,42E-08 | ENSDARG00000029204/ENSDARG00000061303/ENSDARG00000056151/ENSDARG00000052700/ENSDARG00000006008/ENSDARG00000039077 | 4,424140944 |
| *GO:0042440* | pigment metabolic process | 9/146 | 58/20304 | 3,16E-10 | 1,49E-07 | ENSDARG00000029204/ENSDARG00000061303/ENSDARG00000007480/ENSDARG00000056151/ENSDARG00000052700/ENSDARG00000006008/ENSDARG00000038643/ENSDARG00000039077/ENSDARG00000101021 | 3,071748135 |
| *GO:0019748* | secondary metabolic process | 6/146 | 16/20304 | 9,41E-10 | 2,73E-07 | ENSDARG00000029204/ENSDARG00000061303/ENSDARG00000056151/ENSDARG00000052700/ENSDARG00000006008/ENSDARG00000039077 | 3,954137315 |
| *GO:0046148* | pigment biosynthetic process | 8/146 | 46/20304 | 1,22E-09 | 2,73E-07 | ENSDARG00000029204/ENSDARG00000061303/ENSDARG00000007480/ENSDARG00000056151/ENSDARG00000006008/ENSDARG00000038643/ENSDARG00000039077/ENSDARG00000101021 | 3,185766713 |
| *GO:0032438* | melanosome organization | 6/146 | 17/20304 | 1,45E-09 | 2,73E-07 | ENSDARG00000029204/ENSDARG00000056151/ENSDARG00000034572/ENSDARG00000039077/ENSDARG00000033760/ENSDARG00000091298 | 3,893512693 |
| *GO:0048753* | pigment granule organization | 6/146 | 18/20304 | 2,16E-09 | 3,39E-07 | ENSDARG00000029204/ENSDARG00000056151/ENSDARG00000034572/ENSDARG00000039077/ENSDARG00000033760/ENSDARG00000091298 | 3,83635428 |
| *GO:0030318* | melanocyte differentiation | 7/146 | 49/20304 | 5,74E-08 | 7,13E-06 | ENSDARG00000029204/ENSDARG00000061303/ENSDARG00000003732/ENSDARG00000056151/ENSDARG00000024771/ENSDARG00000039077/ENSDARG00000091298 | 2,989056419 |
| *GO:0043473* | pigmentation | 11/146 | 176/20304 | 6,03E-08 | 7,13E-06 | ENSDARG00000029204/ENSDARG00000061303/ENSDARG00000003732/ENSDARG00000056151/ENSDARG00000006008/ENSDARG00000034572/ENSDARG00000033539/ENSDARG00000024771/ENSDARG00000039077/ENSDARG00000033760/ENSDARG00000091298 | 2,162377846 |
| *GO:0048066* | developmental pigmentation | 8/146 | 98/20304 | 5,37E-07 | 5,64E-05 | ENSDARG00000029204/ENSDARG00000061303/ENSDARG00000003732/ENSDARG00000056151/ENSDARG00000006008/ENSDARG00000024771/ENSDARG00000039077/ENSDARG00000091298 | 2,429440631 |
| *GO:0046189* | phenol-containing compound biosynthetic process | 5/146 | 25/20304 | 8,49E-07 | 8,02E-05 | ENSDARG00000029204/ENSDARG00000061303/ENSDARG00000056151/ENSDARG00000006008/ENSDARG00000039077 | 3,325528656 |

| | | | | | | | |
|---|---|---|---|---|---|---|---|
| GO:0050931 | pigment cell differentiation | 7/146 | 74/20304 | 1,04E-06 | 8,89E-05 | ENSDARG00000029204/ENSDARG00000061303/ENSDARG00000003732/ENSDARG00000056151/ENSDARG00000024771/ENSDARG00000039077/ENSDARG00000091298 | 2,576811624 |
| GO:0033059 | cellular pigmentation | 6/146 | 66/20304 | 7,94E-06 | 0,000626 | ENSDARG00000029204/ENSDARG00000056151/ENSDARG00000034572/ENSDARG00000039077/ENSDARG00000033760/ENSDARG00000091298 | 2,537071295 |
| GO:0042541 | hemoglobin biosynthetic process | 4/146 | 21/20304 | 1,4E-05 | 0,001015 | ENSDARG00000101322/ENSDARG00000052700/ENSDARG00000038643/ENSDARG00000077372 | 3,276738492 |
| GO:0018958 | phenol-containing compound metabolic process | 5/146 | 44/20304 | 1,56E-05 | 0,00105 | ENSDARG00000029204/ENSDARG00000061303/ENSDARG00000056151/ENSDARG00000006008/ENSDARG00000039077 | 2,760214847 |
| GO:0020027 | hemoglobin metabolic process | 4/146 | 22/20304 | 1,7E-05 | 0,001069 | ENSDARG00000101322/ENSDARG00000052700/ENSDARG00000038643/ENSDARG00000077372 | 3,230218476 |
| GO:0048872 | homeostasis of number of cells | 8/146 | 167/20304 | 2,85E-05 | 0,001683 | ENSDARG00000055163/ENSDARG00000012881/ENSDARG00000017400/ENSDARG00000101322/ENSDARG00000052700/ENSDARG00000012945/ENSDARG00000038643/ENSDARG00000075500 | 1,896414298 |
| GO:0030218 | erythrocyte differentiation | 7/146 | 130/20304 | 4,35E-05 | 0,002397 | ENSDARG00000055163/ENSDARG00000012881/ENSDARG00000017400/ENSDARG00000101322/ENSDARG00000052700/ENSDARG00000038643/ENSDARG00000075500 | 2,013342267 |
| GO:0034101 | erythrocyte homeostasis | 7/146 | 131/20304 | 4,57E-05 | 0,002397 | ENSDARG00000055163/ENSDARG00000012881/ENSDARG00000017400/ENSDARG00000101322/ENSDARG00000052700/ENSDARG00000038643/ENSDARG00000075500 | 2,005679394 |
| GO:0002262 | myeloid cell homeostasis | 7/146 | 143/20304 | 7,96E-05 | 0,00396 | ENSDARG00000055163/ENSDARG00000012881/ENSDARG00000017400/ENSDARG00000101322/ENSDARG00000052700/ENSDARG00000038643/ENSDARG00000075500 | 1,918032087 |
| GO:1901605 | alpha-amino acid metabolic process | 7/146 | 171/20304 | 0,000242 | 0,010957 | ENSDARG00000035602/ENSDARG00000016733/ENSDARG00000100818/ENSDARG00000020028/ENSDARG00000006008/ENSDARG00000007377/ENSDARG00000039077 | 1,739213161 |
| GO:1901615 | organic hydroxy compound metabolic process | 9/146 | 288/20304 | 0,000243 | 0,010957 | ENSDARG00000001975/ENSDARG00000029204/ENSDARG00000061303/ENSDARG00000077652/ENSDARG00000056151/ENSDARG00000006008/ENSDARG00000042014/ENSDARG00000039077/ENSDARG00000078618 | 1,469230665 |
| GO:0055076 | transition metal ion homeostasis | 5/146 | 80/20304 | 0,00028 | 0,012017 | ENSDARG00000101322/ENSDARG00000052700/ENSDARG00000075641/ENSDARG00000079525/ENSDARG00000077372 | 2,162377846 |
| GO:0048821 | erythrocyte development | 4/146 | 55/20304 | 0,000658 | 0,027044 | ENSDARG00000055163/ENSDARG00000052700/ENSDARG00000038643/ENSDARG00000075500 | 2,313927744 |
| GO:0055072 | iron ion homeostasis | 4/146 | 57/20304 | 0,000754 | 0,029686 | ENSDARG00000101322/ENSDARG00000052700/ENSDARG00000075641/ENSDARG00000077372 | 2,278209662 |
| GO:0055080 | cation homeostasis | 10/146 | 430/20304 | 0,001121 | 0,042377 | ENSDARG00000012881/ENSDARG00000101322/ENSDARG00000052700/ENSDARG00000075641/ENSDARG00000020028/ENSDARG00000089824/ENSDARG00000024771/ENSDARG00000097816/ENSDARG00000079525/ENSDARG00000077372 | 1,173766453 |
| GO:0098771 | inorganic ion homeostasis | 10/146 | 441/20304 | 0,001354 | 0,049204 | ENSDARG00000012881/ENSDARG00000101322/ENSDARG00000052700/ENSDARG00000075641/ENSDARG00000020028/ENSDARG00000089824/ENSDARG0000002477 | 1,148506786 |

| | | | | | |
|---|---|---|---|---|---|
| | | | | | 1/ENSDARG00000097816/ENSDARG00000079525/ENSDARG00000077372 |

**Supplementary Table 6:** Gene Ontology term enrichment for downregulated genes in 5 dpf qKO zebrafish versus WT controls (p.adjust < 0.05). Table includes: GeneRatio (ratio of input genes that are annotated in a term) and BgRatio (ratio of all genes that are annotated in this term), pvalue (raw p-value), p.adjust (Bonferroni adjusted p-values), geneID (identity of the DEG included in the GO category), Enrichment (the log2 of the ratio of the gene set (%) in the DEG and the gene set (%) in the universe).

| *ID* | *Description* | *Gene Ratio* | *BgRatio* | *pvalue* | *p.adjust* | *geneID* | *log2 Enrichment* |
|---|---|---|---|---|---|---|---|
| *GO:0010466* | negative regulation of peptidase activity | 24/444 | 103/20304 | 2,81E-18 | 4,44E-15 | ENSDARG00000077960/ENSDARG00000091800/ENSDARG00000087143/ENSDARG00000091136/ENSDARG00000069293/ENSDARG00000090850/ENSDARG00000021208/ENSDARG00000058053/ENSDARG00000103833/ENSDARG00000102271/ENSDARG00000100051/ENSDARG00000099811/ENSDARG00000101290/ENSDARG00000041645/ENSDARG00000041685/ENSDARG00000061383/ENSDARG00000015065/ENSDARG00000076448/ENSDARG00000102617/ENSDARG00000100795/ENSDARG00000055036/ENSDARG00000026764/ENSDARG00000053973/ENSDARG00000070892 | 2,366073 |
| *GO:0051346* | negative regulation of hydrolase activity | 26/444 | 131/20304 | 7,78E-18 | 6,16E-15 | ENSDARG00000026726/ENSDARG00000077960/ENSDARG00000091800/ENSDARG00000087143/ENSDARG00000091136/ENSDARG00000044365/ENSDARG00000069293/ENSDARG00000090850/ENSDARG00000021208/ENSDARG00000058053/ENSDARG00000103833/ENSDARG00000102271/ENSDARG00000100051/ENSDARG00000099811/ENSDARG00000101290/ENSDARG00000041645/ENSDARG00000041685/ENSDARG00000061383/ENSDARG00000015065/ENSDARG00000076448/ENSDARG00000102617/ENSDARG00000100795/ENSDARG00000055036/ENSDARG00000026764/ENSDARG00000053973/ENSDARG00000070892 | 2,205648 |
| *GO:0045861* | negative regulation of proteolysis | 24/444 | 111/20304 | 1,81E-17 | 9,53E-15 | ENSDARG00000077960/ENSDARG00000091800/ENSDARG00000087143/ENSDARG00000091136/ENSDARG00000069293/ENSDARG00000090850/ENSDARG00000021208/ENSDARG00000058053/ENSDARG00000103833/ENSDARG00000102271/ENSDARG00000100051/ENSDARG00000099811/ENSDARG00000101290/ENSDARG00000041645/ENSDARG00000041685/ENSDARG00000061383/ENSDARG00000015065/ENSDARG00000076448/ENSDARG00000102617/ENSDARG00000100795/ENSDARG00000055036/ENSDARG00000026764/ENSDARG00000053973/ENSDARG00000070892 | 2,291272 |
| *GO:0052547* | regulation of peptidase activity | 24/444 | 156/20304 | 5,84E-14 | 2,31E-11 | ENSDARG00000077960/ENSDARG00000091800/ENSDARG00000087143/ENSDARG00000091136/ENSDARG00000069293/ENSDARG00000090850/ENSDARG00000021208/ENSDARG00000058053/ENSDARG00000103833/ENSDARG00000102271/ENSDARG00000100051/ENSDARG00000099811/ENSDARG00000101290/ENSDARG00000041645/ENSDARG00000041685/ENSDARG00000061383/ENSDARG00000015065/ENSDARG | 1,950946 |

| | | | | | | | |
|---|---|---|---|---|---|---|---|
| | | | | | | 00000076448/ENSDARG00000102617/ENSDARG00000100795/ENSDARG00000055036/ENSDARG00000026764/ENSDARG00000053973/ENSDARG00000070892 | |
| *GO:0043086* | negative regulation of catalytic activity | 26/444 | 211/20304 | 1,08E-12 | 3,41E-10 | ENSDARG00000026726/ENSDARG00000077960/ENSDARG00000091800/ENSDARG00000087143/ENSDARG00000091136/ENSDARG00000044365/ENSDARG00000069293/ENSDARG00000090850/ENSDARG00000021208/ENSDARG00000058053/ENSDARG00000103833/ENSDARG00000102271/ENSDARG00000100051/ENSDARG00000099811/ENSDARG00000101290/ENSDARG00000041645/ENSDARG00000041685/ENSDARG00000061383/ENSDARG00000015065/ENSDARG00000076448/ENSDARG00000102617/ENSDARG00000100795/ENSDARG00000055036/ENSDARG00000026764/ENSDARG00000053973/ENSDARG00000070892 | 1,728987 |
| *GO:0010951* | negative regulation of endopeptidase activity | 14/444 | 61/20304 | 4,11E-11 | 1,09E-08 | ENSDARG00000077960/ENSDARG00000091800/ENSDARG00000087143/ENSDARG00000091136/ENSDARG00000069293/ENSDARG00000090850/ENSDARG00000021208/ENSDARG00000058053/ENSDARG00000061383/ENSDARG00000076448/ENSDARG00000100795/ENSDARG00000026764/ENSDARG00000053973/ENSDARG00000070892 | 2,350932 |
| *GO:0044092* | negative regulation of molecular function | 26/444 | 254/20304 | 7,47E-11 | 1,69E-08 | ENSDARG00000026726/ENSDARG00000077960/ENSDARG00000091800/ENSDARG00000087143/ENSDARG00000091136/ENSDARG00000044365/ENSDARG00000069293/ENSDARG00000090850/ENSDARG00000021208/ENSDARG00000058053/ENSDARG00000103833/ENSDARG00000102271/ENSDARG00000100051/ENSDARG00000099811/ENSDARG00000101290/ENSDARG00000041645/ENSDARG00000041685/ENSDARG00000061383/ENSDARG00000015065/ENSDARG00000076448/ENSDARG00000102617/ENSDARG00000100795/ENSDARG00000055036/ENSDARG00000026764/ENSDARG00000053973/ENSDARG00000070892 | 1,543511 |
| *GO:0030162* | regulation of proteolysis | 24/444 | 231/20304 | 2,99E-10 | 5,91E-08 | ENSDARG00000077960/ENSDARG00000091800/ENSDARG00000087143/ENSDARG00000091136/ENSDARG00000069293/ENSDARG00000090850/ENSDARG00000021208/ENSDARG00000058053/ENSDARG00000103833/ENSDARG00000102271/ENSDARG00000100051/ENSDARG00000099811/ENSDARG00000101290/ENSDARG00000041645/ENSDARG00000041685/ENSDARG00000061383/ENSDARG00000015065/ENSDARG00000076448/ENSDARG00000102617/ENSDARG00000100795/ENSDARG00000055036/ENSDARG00000026764/ENSDARG00000053973/ENSDARG00000070892 | 1,558385 |
| *GO:0050776* | regulation of immune response | 23/444 | 225/20304 | 9,71E-10 | 1,71E-07 | ENSDARG00000026726/ENSDARG00000093750/ENSDARG00000093198/ENSDARG00000096586/ENSDARG00000094408/ENSDARG00000006848/ENSDARG00000032010/ENSDARG00000058045/ENSDARG00000026663/ENSDARG00000058053/ENSDARG00000102525/ENSDARG00000076306/ENSDARG00000017565/ENSDARG00000039579/ENSDARG00000076139/ENSDARG00000024631/ENSDARG00000015065/ENSDARG00000044612/ENSDARG00000005616/ENSDARG00000036628/ENSDARG00000043249/ENSDARG00000001818/ENSDARG00000093068 | 1,542142 |
| *GO:0002684* | positive regulation of immune system process | 22/444 | 219/20304 | 3,13E-09 | 4,95E-07 | ENSDARG00000101030/ENSDARG00000101675/ENSDARG00000026726/ENSDARG00000093198/ENSDARG00000096586/ENSDARG00000094408/ENSDARG00000006848/ENSDARG00000058053/ENSDARG00000031745/ENSDARG00000075932/ENSDARG00000102525/ENSDARG00000102986/ENSDARG00000017565/ENSDARG00000039579/ENSDARG00000076139/ENSDARG00000 | 1,524719 |

| | | | | | | | |
|---|---|---|---|---|---|---|---|
| | | | | | | 056330/ENSDARG00000015065/ENSDARG00000044612/ENSDARG00000005616/ENSDARG00000036628/ENSDARG00000001818/ENSDARG00000093068 | |
| *GO:0009410* | response to xenobiotic stimulus | 16/444 | 117/20304 | 5,57E-09 | 7,69E-07 | ENSDARG00000039211/ENSDARG00000092091/ENSDARG00000026726/ENSDARG00000098995/ENSDARG00000103295/ENSDARG00000042953/ENSDARG00000042956/ENSDARG00000021172/ENSDARG00000042978/ENSDARG00000042980/ENSDARG00000041540/ENSDARG00000018361/ENSDARG00000091211/ENSDARG00000102805/ENSDARG00000045442/ENSDARG00000043997 | 1,833163 |
| *GO:0002252* | immune effector process | 18/444 | 151/20304 | 5,83E-09 | 7,69E-07 | ENSDARG00000101030/ENSDARG00000101675/ENSDARG00000026726/ENSDARG00000093198/ENSDARG00000096586/ENSDARG00000094408/ENSDARG00000058053/ENSDARG00000031745/ENSDARG00000075932/ENSDARG00000076306/ENSDARG00000039579/ENSDARG00000056330/ENSDARG00000015065/ENSDARG00000044612/ENSDARG00000005616/ENSDARG00000036628/ENSDARG00000001818/ENSDARG00000093068 | 1,695841 |
| *GO:0051336* | regulation of hydrolase activity | 31/444 | 429/20304 | 6,68E-09 | 8,14E-07 | ENSDARG00000079892/ENSDARG00000026726/ENSDARG00000077960/ENSDARG00000091800/ENSDARG00000087143/ENSDARG00000091136/ENSDARG00000044365/ENSDARG00000069293/ENSDARG00000090850/ENSDARG00000021208/ENSDARG00000058053/ENSDARG00000103833/ENSDARG00000102271/ENSDARG00000100051/ENSDARG00000099811/ENSDARG00000101290/ENSDARG00000041645/ENSDARG00000041685/ENSDARG00000061383/ENSDARG00000015065/ENSDARG00000076448/ENSDARG00000102617/ENSDARG00000100795/ENSDARG00000055036/ENSDARG00000039351/ENSDARG00000094002/ENSDARG00000026764/ENSDARG00000053973/ENSDARG00000070892/ENSDARG00000099401/ENSDARG00000102945 | 1,195279 |
| *GO:0002682* | regulation of immune system process | 29/444 | 385/20304 | 8,27E-09 | 9,35E-07 | ENSDARG00000101030/ENSDARG00000101675/ENSDARG00000026726/ENSDARG00000093750/ENSDARG00000093198/ENSDARG00000096586/ENSDARG00000094408/ENSDARG00000006848/ENSDARG00000032010/ENSDARG00000058045/ENSDARG00000026663/ENSDARG00000058053/ENSDARG00000031745/ENSDARG00000075932/ENSDARG00000102525/ENSDARG00000102986/ENSDARG00000076306/ENSDARG00000017565/ENSDARG00000039579/ENSDARG00000076139/ENSDARG00000024631/ENSDARG00000056330/ENSDARG00000015065/ENSDARG00000044612/ENSDARG00000005616/ENSDARG00000036628/ENSDARG00000043249/ENSDARG00000001818/ENSDARG00000093068 | 1,236801 |
| *GO:0071466* | cellular response to xenobiotic stimulus | 14/444 | 93/20304 | 1,43E-08 | 1,51E-06 | ENSDARG00000039211/ENSDARG00000092091/ENSDARG00000098995/ENSDARG00000042953/ENSDARG00000042956/ENSDARG00000021172/ENSDARG00000042978/ENSDARG00000042980/ENSDARG00000041540/ENSDARG00000018361/ENSDARG00000091211/ENSDARG00000102805/ENSDARG00000045442/ENSDARG00000043997 | 1,929206 |
| *GO:0006959* | humoral immune response | 13/444 | 88/20304 | 5,95E-08 | 5,88E-06 | ENSDARG00000102776/ENSDARG00000093198/ENSDARG00000096586/ENSDARG00000094408/ENSDARG00000058045/ENSDARG00000058053/ENSDARG00000039579/ENSDARG00000015065/ENSDARG00000044612/ENSDARG00000005616/ENSDARG00000092044/ENSDARG00000001818/ENSDARG00000093068 | 1,910361 |

| | | | | | | | |
|---|---|---|---|---|---|---|---|
| GO:0006805 | xenobiotic metabolic process | 11/444 | 60/20304 | 6,31E-08 | 5,88E-06 | ENSDARG00000092091/ENSDARG00000098995/ENSDARG00000042953/ENSDARG00000042956/ENSDARG00000021172/ENSDARG00000042978/ENSDARG00000042980/ENSDARG00000041540/ENSDARG00000018361/ENSDARG00000102805/ENSDARG00000043997 | 2,126299 |
| GO:0050778 | positive regulation of immune response | 16/444 | 146/20304 | 1,35E-07 | 1,18E-05 | ENSDARG00000026726/ENSDARG00000093198/ENSDARG00000096586/ENSDARG00000094408/ENSDARG00000006848/ENSDARG00000058053/ENSDARG00000102525/ENSDARG00000017565/ENSDARG00000039579/ENSDARG00000076139/ENSDARG00000015065/ENSDARG00000044612/ENSDARG00000005616/ENSDARG00000036628/ENSDARG00000001818/ENSDARG00000093068 | 1,611731 |
| GO:0052548 | regulation of endopeptidase activity | 14/444 | 113/20304 | 1,78E-07 | 1,48E-05 | ENSDARG00000077960/ENSDARG00000091800/ENSDARG00000087143/ENSDARG00000091136/ENSDARG00000069293/ENSDARG00000090850/ENSDARG00000021208/ENSDARG00000058053/ENSDARG00000061383/ENSDARG00000076448/ENSDARG00000100795/ENSDARG00000026764/ENSDARG00000053973/ENSDARG00000070892 | 1,734418 |
| GO:0032269 | negative regulation of cellular protein metabolic process | 24/444 | 329/20304 | 2,95E-07 | 2,33E-05 | ENSDARG00000077960/ENSDARG00000091800/ENSDARG00000087143/ENSDARG00000091136/ENSDARG00000069293/ENSDARG00000090850/ENSDARG00000021208/ENSDARG00000058053/ENSDARG00000103833/ENSDARG00000102271/ENSDARG00000100051/ENSDARG00000099811/ENSDARG00000101290/ENSDARG00000041645/ENSDARG00000041685/ENSDARG00000061383/ENSDARG00000015065/ENSDARG00000076448/ENSDARG00000102617/ENSDARG00000100795/ENSDARG00000055036/ENSDARG00000026764/ENSDARG00000053973/ENSDARG00000070892 | 1,204745 |
| GO:0042178 | xenobiotic catabolic process | 9/444 | 44/20304 | 3,79E-07 | 2,86E-05 | ENSDARG00000092091/ENSDARG00000098995/ENSDARG00000042953/ENSDARG00000042956/ENSDARG00000021172/ENSDARG00000042978/ENSDARG00000042980/ENSDARG00000102805/ENSDARG00000043997 | 2,235784 |
| GO:0051248 | negative regulation of protein metabolic process | 24/444 | 338/20304 | 4,81E-07 | 3,46E-05 | ENSDARG00000077960/ENSDARG00000091800/ENSDARG00000087143/ENSDARG00000091136/ENSDARG00000069293/ENSDARG00000090850/ENSDARG00000021208/ENSDARG00000058053/ENSDARG00000103833/ENSDARG00000102271/ENSDARG00000100051/ENSDARG00000099811/ENSDARG00000101290/ENSDARG00000041645/ENSDARG00000041685/ENSDARG00000061383/ENSDARG00000015065/ENSDARG00000076448/ENSDARG00000102617/ENSDARG00000100795/ENSDARG00000055036/ENSDARG00000026764/ENSDARG00000053973/ENSDARG00000070892 | 1,177757 |
| GO:0006956 | complement activation | 10/444 | 59/20304 | 5,45E-07 | 3,75E-05 | ENSDARG00000093198/ENSDARG00000096586/ENSDARG00000094408/ENSDARG00000058053/ENSDARG00000039579/ENSDARG00000015065/ENSDARG00000044612/ENSDARG00000005616/ENSDARG00000001818/ENSDARG00000093068 | 2,047796 |
| GO:0007601 | visual perception | 17/444 | 195/20304 | 1,46E-06 | 9,62E-05 | ENSDARG00000036140/ENSDARG00000004459/ENSDARG00000011886/ENSDARG00000102129/ENSDARG00000053502/ENSDARG00000040736/ENSDARG00000040738/ENSDARG00000115234/ENSDARG00000044875/ENSDARG00000076790/ENSDARG00000002193/ENSDARG00000000380/ENSDARG00000055534/ENSDARG00000075067/ENSDARG00000013393/ENSDARG00000070666/ENSDARG00000044862 | 1,382962 |

| | | | | | | | |
|---|---|---|---|---|---|---|---|
| GO:0034097 | response to cytokine | 18/444 | 222/20304 | 2,05E-06 | 0,00013 | ENSDARG00000102776/ENSDARG00000105263/ENSDARG00000100738/ENSDARG00000026726/ENSDARG00000052413/ENSDARG00000091993/ENSDARG00000053620/ENSDARG00000006848/ENSDARG00000095475/ENSDARG00000087706/ENSDARG00000102986/ENSDARG00000024631/ENSDARG00000036628/ENSDARG00000100795/ENSDARG00000039351/ENSDARG00000094002/ENSDARG00000099401/ENSDARG00000102945 | 1,310443 |
| GO:0050953 | sensory perception of light stimulus | 17/444 | 201/20304 | 2,21E-06 | 0,000135 | ENSDARG00000036140/ENSDARG00000004459/ENSDARG00000011886/ENSDARG00000102129/ENSDARG00000053502/ENSDARG00000040736/ENSDARG00000040738/ENSDARG00000115234/ENSDARG00000044875/ENSDARG00000076790/ENSDARG00000002193/ENSDARG00000000380/ENSDARG00000055534/ENSDARG00000075067/ENSDARG00000013393/ENSDARG00000070666/ENSDARG00000044862 | 1,352657 |
| GO:0002253 | activation of immune response | 13/444 | 124/20304 | 3,33E-06 | 0,000195 | ENSDARG00000093198/ENSDARG00000096586/ENSDARG00000094408/ENSDARG00000058053/ENSDARG00000102525/ENSDARG00000017565/ENSDARG00000039579/ENSDARG00000076139/ENSDARG00000015065/ENSDARG00000044612/ENSDARG00000005616/ENSDARG00000001818/ENSDARG00000093068 | 1,567416 |
| GO:0019221 | cytokine-mediated signaling pathway | 16/444 | 188/20304 | 4,05E-06 | 0,000229 | ENSDARG00000102776/ENSDARG00000105263/ENSDARG00000100738/ENSDARG00000052413/ENSDARG00000091993/ENSDARG00000053620/ENSDARG00000006848/ENSDARG00000095475/ENSDARG00000087706/ENSDARG00000102986/ENSDARG00000024631/ENSDARG00000036628/ENSDARG00000039351/ENSDARG00000094002/ENSDARG00000099401/ENSDARG00000102945 | 1,358895 |
| GO:0042110 | T cell activation | 13/444 | 127/20304 | 4,35E-06 | 0,000238 | ENSDARG00000101030/ENSDARG00000101675/ENSDARG00000026726/ENSDARG00000091993/ENSDARG00000087706/ENSDARG00000031745/ENSDARG00000075932/ENSDARG00000102525/ENSDARG00000076306/ENSDARG00000017565/ENSDARG00000056330/ENSDARG00000036628/ENSDARG00000052122 | 1,543511 |
| GO:0002250 | adaptive immune response | 11/444 | 93/20304 | 5,88E-06 | 0,00031 | ENSDARG00000101030/ENSDARG00000101675/ENSDARG00000026726/ENSDARG00000103077/ENSDARG00000031745/ENSDARG00000075932/ENSDARG00000076306/ENSDARG00000017565/ENSDARG00000056330/ENSDARG00000044612/ENSDARG00000052122 | 1,688044 |
| GO:0071345 | cellular response to cytokine stimulus | 16/444 | 196/20304 | 6,92E-06 | 0,000353 | ENSDARG00000102776/ENSDARG00000105263/ENSDARG00000100738/ENSDARG00000052413/ENSDARG00000091993/ENSDARG00000053620/ENSDARG00000006848/ENSDARG00000095475/ENSDARG00000087706/ENSDARG00000102986/ENSDARG00000024631/ENSDARG00000036628/ENSDARG00000039351/ENSDARG00000094002/ENSDARG00000099401/ENSDARG00000102945 | 1,317223 |
| GO:0098542 | defense response to other organism | 23/444 | 374/20304 | 9,29E-06 | 0,000459 | ENSDARG00000093750/ENSDARG00000062998/ENSDARG00000053802/ENSDARG00000096586/ENSDARG00000094408/ENSDARG00000006848/ENSDARG00000032010/ENSDARG00000095475/ENSDARG00000058045/ENSDARG00000026663/ENSDARG00000102175/ENSDARG00000015752/ENSDARG00000102525/ENSDARG00000102986/ENSDARG00000039579/ENSDARG00000024631/ENSDARG00000090537/ENSDARG00000092044/ENSDARG00000043249/ENSDARG00000039351/ENSDARG00000094002/ENSDARG00000099401/ENSDARG00000102945 | 1,033987 |

| | | | | | | | |
|---|---|---|---|---|---|---|---|
| *GO:0007600* | sensory perception | 25/444 | 441/20304 | 1,55E-05 | 0,000738 | ENSDARG00000036140/ENSDARG00000004459/ENSDARG00000011886/ENSDARG00000102129/ENSDARG00000078474/ENSDARG00000053502/ENSDARG00000040736/ENSDARG00000040738/ENSDARG00000115234/ENSDARG00000044875/ENSDARG00000076790/ENSDARG00000056458/ENSDARG00000002193/ENSDARG00000000380/ENSDARG00000055534/ENSDARG00000075067/ENSDARG00000041033/ENSDARG00000041030/ENSDARG00000075128/ENSDARG00000013393/ENSDARG00000105757/ENSDARG00000044748/ENSDARG00000070666/ENSDARG00000044862/ENSDARG00000105708 | 0,95258 |
| *GO:0043207* | response to external biotic stimulus | 27/444 | 499/20304 | 1,63E-05 | 0,000738 | ENSDARG00000102776/ENSDARG00000105263/ENSDARG00000093750/ENSDARG00000062998/ENSDARG00000053802/ENSDARG00000096586/ENSDARG00000094408/ENSDARG00000006848/ENSDARG00000032010/ENSDARG00000095475/ENSDARG00000058045/ENSDARG00000026663/ENSDARG00000102175/ENSDARG00000015752/ENSDARG00000102525/ENSDARG00000102986/ENSDARG00000039579/ENSDARG00000024631/ENSDARG00000090537/ENSDARG00000000656/ENSDARG00000092044/ENSDARG00000043249/ENSDARG00000039351/ENSDARG00000094002/ENSDARG00000056462/ENSDARG00000099401/ENSDARG00000102945 | 0,905979 |
| *GO:0051707* | response to other organism | 27/444 | 499/20304 | 1,63E-05 | 0,000738 | ENSDARG00000102776/ENSDARG00000105263/ENSDARG00000093750/ENSDARG00000062998/ENSDARG00000053802/ENSDARG00000096586/ENSDARG00000094408/ENSDARG00000006848/ENSDARG00000032010/ENSDARG00000095475/ENSDARG00000058045/ENSDARG00000026663/ENSDARG00000102175/ENSDARG00000015752/ENSDARG00000102525/ENSDARG00000102986/ENSDARG00000039579/ENSDARG00000024631/ENSDARG00000090537/ENSDARG00000000656/ENSDARG00000092044/ENSDARG00000043249/ENSDARG00000039351/ENSDARG00000094002/ENSDARG00000056462/ENSDARG00000099401/ENSDARG00000102945 | 0,905979 |
| *GO:0002396* | MHC protein complex assembly | 5/444 | 17/20304 | 2,43E-05 | 0,000964 | ENSDARG00000101030/ENSDARG00000101675/ENSDARG00000031745/ENSDARG00000075932/ENSDARG00000056330 | 2,598973 |
| *GO:0002399* | MHC class II protein complex assembly | 5/444 | 17/20304 | 2,43E-05 | 0,000964 | ENSDARG00000101030/ENSDARG00000101675/ENSDARG00000031745/ENSDARG00000075932/ENSDARG00000056330 | 2,598973 |
| *GO:0002501* | peptide antigen assembly with MHC protein complex | 5/444 | 17/20304 | 2,43E-05 | 0,000964 | ENSDARG00000101030/ENSDARG00000101675/ENSDARG00000031745/ENSDARG00000075932/ENSDARG00000056330 | 2,598973 |
| *GO:0002503* | peptide antigen assembly with MHC class II protein complex | 5/444 | 17/20304 | 2,43E-05 | 0,000964 | ENSDARG00000101030/ENSDARG00000101675/ENSDARG00000031745/ENSDARG00000075932/ENSDARG00000056330 | 2,598973 |
| *GO:0019886* | antigen processing and presentation of exogenous peptide antigen via MHC class II | 5/444 | 17/20304 | 2,43E-05 | 0,000964 | ENSDARG00000101030/ENSDARG00000101675/ENSDARG00000031745/ENSDARG00000075932/ENSDARG00000056330 | 2,598973 |
| *GO:0045087* | innate immune response | 17/444 | 244/20304 | 2,87E-05 | 0,00111 | ENSDARG00000096586/ENSDARG00000094408/ENSDARG00000006848/ENSDARG00000032010/ENSDARG00000095475/ENSDARG00000058045/ENSDARG00000026663/ENSDARG00000102175/ENSDARG00000015752/ENSDARG00000102525/ENSDARG00000102986/ENSDARG00000039579/ENSDARG00000024631/ENSDARG0000003935 | 1,158794 |

| | | | | | | | |
|---|---|---|---|---|---|---|---|
| | | | | | | 1/ENSDARG00000094002/ENSDARG00000099401/ENSDARG00000102945 | |
| GO:0009074 | aromatic amino acid family catabolic process | 6/444 | 29/20304 | 3,28E-05 | 0,001191 | ENSDARG00000018351/ENSDARG00000044935/ENSDARG00000071429/ENSDARG00000069630/ENSDARG00000023176/ENSDARG00000058005 | 2,247212 |
| GO:0002478 | antigen processing and presentation of exogenous peptide antigen | 5/444 | 18/20304 | 3,31E-05 | 0,001191 | ENSDARG00000101030/ENSDARG00000101675/ENSDARG00000031745/ENSDARG00000075932/ENSDARG00000056330 | 2,541815 |
| GO:0019884 | antigen processing and presentation of exogenous antigen | 5/444 | 18/20304 | 3,31E-05 | 0,001191 | ENSDARG00000101030/ENSDARG00000101675/ENSDARG00000031745/ENSDARG00000075932/ENSDARG00000056330 | 2,541815 |
| GO:0002495 | antigen processing and presentation of peptide antigen via MHC class II | 5/444 | 20/20304 | 5,78E-05 | 0,002033 | ENSDARG00000101030/ENSDARG00000101675/ENSDARG00000031745/ENSDARG00000075932/ENSDARG00000056330 | 2,436454 |
| GO:0002460 | adaptive immune response based on somatic recombination of immune receptors built from immunoglobulin superfamily domains | 8/444 | 64/20304 | 7,44E-05 | 0,002509 | ENSDARG00000101030/ENSDARG00000101675/ENSDARG00000026726/ENSDARG00000031745/ENSDARG00000075932/ENSDARG00000076306/ENSDARG00000056330/ENSDARG00000044612 | 1,743307 |
| GO:0002504 | antigen processing and presentation of peptide or polysaccharide antigen via MHC class II | 5/444 | 21/20304 | 7,45E-05 | 0,002509 | ENSDARG00000101030/ENSDARG00000101675/ENSDARG00000031745/ENSDARG00000075932/ENSDARG00000056330 | 2,387664 |
| GO:0006144 | purine nucleobase metabolic process | 5/444 | 22/20304 | 9,47E-05 | 0,003122 | ENSDARG00000038293/ENSDARG00000037191/ENSDARG00000042562/ENSDARG00000055240/ENSDARG00000007024 | 2,341144 |
| GO:0009112 | nucleobase metabolic process | 6/444 | 35/20304 | 0,0001 | 0,003237 | ENSDARG00000038293/ENSDARG00000098386/ENSDARG00000037191/ENSDARG00000042562/ENSDARG00000055240/ENSDARG00000007024 | 2,05916 |
| GO:0046649 | lymphocyte activation | 13/444 | 171/20304 | 0,000103 | 0,003274 | ENSDARG00000101030/ENSDARG00000101675/ENSDARG00000026726/ENSDARG00000091993/ENSDARG00000087706/ENSDARG00000031745/ENSDARG00000075932/ENSDARG00000102525/ENSDARG00000076306/ENSDARG00000017565/ENSDARG00000056330/ENSDARG00000036628/ENSDARG00000052122 | 1,246034 |
| GO:0019882 | antigen processing and presentation | 6/444 | 36/20304 | 0,000118 | 0,003664 | ENSDARG00000101030/ENSDARG00000101675/ENSDARG00000031745/ENSDARG00000075932/ENSDARG00000056330/ENSDARG00000036628 | 2,030989 |
| GO:0045321 | leukocyte activation | 14/444 | 203/20304 | 0,00016 | 0,00487 | ENSDARG00000101030/ENSDARG00000101675/ENSDARG00000026726/ENSDARG00000091993/ENSDARG00000079829/ENSDARG00000087706/ENSDARG00000031745/ENSDARG00000075932/ENSDARG00000102525/ENSDARG00000076306/ENSDARG00000017565/ENSDARG00000056330/ENSDARG00000036628/ENSDARG00000052122 | 1,1486 |
| GO:0006570 | tyrosine metabolic process | 4/444 | 14/20304 | 0,00019 | 0,005667 | ENSDARG00000018351/ENSDARG00000044935/ENSDARG00000069630/ENSDARG00000058005 | 2,569986 |
| GO:0050870 | positive regulation of T cell activation | 6/444 | 40/20304 | 0,000216 | 0,006218 | ENSDARG00000101030/ENSDARG00000101675/ENSDARG00000026726/ENSDARG0 | 1,925629 |

| | | | | | | | |
|---|---|---|---|---|---|---|---|
| | | | | | | 0000031745/ENSDARG00000075932/ENSDARG00000056330 | |
| GO:1903039 | positive regulation of leukocyte cell-cell adhesion | 6/444 | 40/20304 | 0,000216 | 0,006218 | ENSDARG00000101030/ENSDARG00000101675/ENSDARG00000026726/ENSDARG00000031745/ENSDARG00000075932/ENSDARG00000056330 | 1,925629 |
| GO:0002381 | immunoglobulin production involved in immunoglobulin-mediated immune response | 5/444 | 26/20304 | 0,00022 | 0,00622 | ENSDARG00000101030/ENSDARG00000101675/ENSDARG00000031745/ENSDARG00000075932/ENSDARG00000056330 | 2,17409 |
| GO:0009072 | aromatic amino acid family metabolic process | 6/444 | 41/20304 | 0,000248 | 0,0069 | ENSDARG00000018351/ENSDARG00000044935/ENSDARG00000071429/ENSDARG00000069630/ENSDARG00000023176/ENSDARG00000058005 | 1,900936 |
| GO:0045785 | positive regulation of cell adhesion | 7/444 | 58/20304 | 0,000262 | 0,00715 | ENSDARG00000101030/ENSDARG00000101675/ENSDARG00000026726/ENSDARG00000076800/ENSDARG00000031745/ENSDARG00000075932/ENSDARG00000056330 | 1,708216 |
| GO:1903037 | regulation of leukocyte cell-cell adhesion | 7/444 | 59/20304 | 0,000292 | 0,007827 | ENSDARG00000101030/ENSDARG00000101675/ENSDARG00000026726/ENSDARG00000031745/ENSDARG00000075932/ENSDARG00000076306/ENSDARG00000056330 | 1,691121 |
| GO:0022409 | positive regulation of cell-cell adhesion | 6/444 | 43/20304 | 0,000325 | 0,008567 | ENSDARG00000101030/ENSDARG00000101675/ENSDARG00000026726/ENSDARG00000031745/ENSDARG00000075932/ENSDARG00000056330 | 1,853308 |
| GO:0048002 | antigen processing and presentation of peptide antigen | 5/444 | 30/20304 | 0,000444 | 0,011511 | ENSDARG00000101030/ENSDARG00000101675/ENSDARG00000031745/ENSDARG00000075932/ENSDARG00000056330 | 2,030989 |
| GO:0060326 | cell chemotaxis | 11/444 | 150/20304 | 0,000473 | 0,012069 | ENSDARG00000102776/ENSDARG00000026726/ENSDARG00000079829/ENSDARG00000038968/ENSDARG00000102986/ENSDARG00000039351/ENSDARG00000099738/ENSDARG00000094002/ENSDARG00000074851/ENSDARG00000099401/ENSDARG00000102945 | 1,210009 |
| GO:0007159 | leukocyte cell-cell adhesion | 7/444 | 64/20304 | 0,000484 | 0,012155 | ENSDARG00000101030/ENSDARG00000101675/ENSDARG00000026726/ENSDARG00000031745/ENSDARG00000075932/ENSDARG00000076306/ENSDARG00000056330 | 1,609776 |
| GO:0002440 | production of molecular mediator of immune response | 6/444 | 48/20304 | 0,000596 | 0,014746 | ENSDARG00000101030/ENSDARG00000101675/ENSDARG00000031745/ENSDARG00000075932/ENSDARG00000056330/ENSDARG00000036628 | 1,743307 |
| GO:0070374 | positive regulation of ERK1 and ERK2 cascade | 8/444 | 88/20304 | 0,000685 | 0,016679 | ENSDARG00000091993/ENSDARG00000087706/ENSDARG00000036628/ENSDARG00000039351/ENSDARG00000094002/ENSDARG00000109371/ENSDARG00000099401/ENSDARG00000102945 | 1,424853 |
| GO:0001775 | cell activation | 14/444 | 236/20304 | 0,000735 | 0,017627 | ENSDARG00000101030/ENSDARG00000101675/ENSDARG00000026726/ENSDARG00000091993/ENSDARG00000079829/ENSDARG00000087706/ENSDARG00000031745/ENSDARG00000075932/ENSDARG00000102525/ENSDARG00000076306/ENSDARG00000017565/ENSDARG00000056330/ENSDARG00000036628/ENSDARG00000052122 | 0,997974 |
| GO:0001523 | retinoid metabolic process | 6/444 | 51/20304 | 0,000828 | 0,019564 | ENSDARG00000103659/ENSDARG00000027620/ENSDARG00000038742/ENSDARG00000019260/ENSDARG00000037191/ENSDARG00000044982 | 1,682682 |

| | | | | | | | |
|---|---|---|---|---|---|---|---|
| *GO:0016101* | diterpenoid metabolic process | 6/444 | 52/20304 | 0,000919 | 0,021394 | ENSDARG00000103659/ENSDARG00000027620/ENSDARG00000038742/ENSDARG00000019260/ENSDARG00000037191/ENSDARG00000044982 | 1,663264 |
| *GO:0016064* | immunoglobulin mediated immune response | 6/444 | 53/20304 | 0,001018 | 0,023011 | ENSDARG00000101030/ENSDARG00000101675/ENSDARG00000031745/ENSDARG00000075932/ENSDARG00000056330/ENSDARG00000044612 | 1,644216 |
| *GO:0019724* | B cell mediated immunity | 6/444 | 53/20304 | 0,001018 | 0,023011 | ENSDARG00000101030/ENSDARG00000101675/ENSDARG00000031745/ENSDARG00000075932/ENSDARG00000056330/ENSDARG00000044612 | 1,644216 |
| *GO:0050863* | regulation of T cell activation | 7/444 | 74/20304 | 0,001163 | 0,025922 | ENSDARG00000101030/ENSDARG00000101675/ENSDARG00000026726/ENSDARG00000031745/ENSDARG00000075932/ENSDARG00000076306/ENSDARG00000056330 | 1,464594 |
| *GO:0002377* | immunoglobulin production | 5/444 | 37/20304 | 0,001197 | 0,026177 | ENSDARG00000101030/ENSDARG00000101675/ENSDARG00000031745/ENSDARG00000075932/ENSDARG00000056330 | 1,821269 |
| *GO:0050777* | negative regulation of immune response | 4/444 | 22/20304 | 0,001207 | 0,026177 | ENSDARG00000026726/ENSDARG00000058053/ENSDARG00000076306/ENSDARG00000024631 | 2,118001 |
| *GO:0097530* | granulocyte migration | 9/444 | 120/20304 | 0,001298 | 0,027765 | ENSDARG00000102776/ENSDARG00000026726/ENSDARG00000102986/ENSDARG00000039351/ENSDARG00000068923/ENSDARG00000094002/ENSDARG00000074851/ENSDARG00000099401/ENSDARG00000102945 | 1,232481 |
| *GO:0006721* | terpenoid metabolic process | 6/444 | 56/20304 | 0,001362 | 0,02875 | ENSDARG00000103659/ENSDARG00000027620/ENSDARG00000038742/ENSDARG00000019260/ENSDARG00000037191/ENSDARG00000044982 | 1,589156 |
| *GO:0071621* | granulocyte chemotaxis | 8/444 | 98/20304 | 0,001388 | 0,028912 | ENSDARG00000102776/ENSDARG00000026726/ENSDARG00000102986/ENSDARG00000039351/ENSDARG00000094002/ENSDARG00000074851/ENSDARG00000099401/ENSDARG00000102945 | 1,317223 |
| *GO:0002449* | lymphocyte mediated immunity | 6/444 | 57/20304 | 0,001495 | 0,029754 | ENSDARG00000101030/ENSDARG00000101675/ENSDARG00000031745/ENSDARG00000075932/ENSDARG00000056330/ENSDARG00000044612 | 1,571457 |
| *GO:0006576* | cellular biogenic amine metabolic process | 6/444 | 57/20304 | 0,001495 | 0,029754 | ENSDARG00000035652/ENSDARG00000041540/ENSDARG00000018361/ENSDARG00000071429/ENSDARG00000023176/ENSDARG00000086458 | 1,571457 |
| *GO:0002698* | negative regulation of immune effector process | 3/444 | 11/20304 | 0,001504 | 0,029754 | ENSDARG00000026726/ENSDARG00000058053/ENSDARG00000076306 | 2,523466 |
| *GO:0015696* | ammonium transport | 3/444 | 11/20304 | 0,001504 | 0,029754 | ENSDARG00000007080/ENSDARG00000041848/ENSDARG00000036722 | 2,523466 |
| *GO:0051251* | positive regulation of lymphocyte activation | 6/444 | 58/20304 | 0,001637 | 0,031602 | ENSDARG00000101030/ENSDARG00000101675/ENSDARG00000026726/ENSDARG00000031745/ENSDARG00000075932/ENSDARG00000056330 | 1,554065 |
| *GO:0070098* | chemokine-mediated signaling pathway | 6/444 | 58/20304 | 0,001637 | 0,031602 | ENSDARG00000102776/ENSDARG00000105263/ENSDARG00000039351/ENSDARG00000094002/ENSDARG00000099401/ENSDARG00000102945 | 1,554065 |
| *GO:0070372* | regulation of ERK1 and ERK2 cascade | 8/444 | 102/20304 | 0,001794 | 0,034212 | ENSDARG00000091993/ENSDARG00000087706/ENSDARG00000036628/ENSDARG00000039351/ENSDARG00000094002/ENSDARG00000109371/ENSDARG00000099401/ENSDARG00000102945 | 1,277217 |
| *GO:0042574* | retinal metabolic process | 3/444 | 12/20304 | 0,001973 | 0,036578 | ENSDARG00000103659/ENSDARG00000027620/ENSDARG00000019260 | 2,436454 |

| | | | | | | | |
|---|---|---|---|---|---|---|---|
| *GO:0006026* | aminoglycan catabolic process | 4/444 | 25/20304 | 0,001982 | 0,036578 | ENSDARG00000062998/ENSDARG00000093193/ENSDARG00000100635/ENSDARG00000099185 | 1,990167 |
| *GO:0070371* | ERK1 and ERK2 cascade | 8/444 | 104/20304 | 0,002029 | 0,036578 | ENSDARG00000091993/ENSDARG00000087706/ENSDARG00000036628/ENSDARG00000039351/ENSDARG00000094002/ENSDARG00000109371/ENSDARG00000099401/ENSDARG00000102945 | 1,257799 |
| *GO:0030155* | regulation of cell adhesion | 10/444 | 154/20304 | 0,002117 | 0,036578 | ENSDARG00000079892/ENSDARG00000101030/ENSDARG00000101675/ENSDARG00000026726/ENSDARG00000076800/ENSDARG00000031745/ENSDARG00000075932/ENSDARG00000076306/ENSDARG00000056330/ENSDARG00000039145 | 1,088381 |
| *GO:0002696* | positive regulation of leukocyte activation | 6/444 | 61/20304 | 0,002126 | 0,036578 | ENSDARG00000101030/ENSDARG00000101675/ENSDARG00000026726/ENSDARG00000031745/ENSDARG00000075932/ENSDARG00000056330 | 1,503634 |
| *GO:0044106* | cellular amine metabolic process | 6/444 | 61/20304 | 0,002126 | 0,036578 | ENSDARG00000035652/ENSDARG00000041540/ENSDARG00000018361/ENSDARG00000071429/ENSDARG00000023176/ENSDARG00000086458 | 1,503634 |
| *GO:0050867* | positive regulation of cell activation | 6/444 | 61/20304 | 0,002126 | 0,036578 | ENSDARG00000101030/ENSDARG00000101675/ENSDARG00000026726/ENSDARG00000031745/ENSDARG00000075932/ENSDARG00000056330 | 1,503634 |
| *GO:1990868* | response to chemokine | 6/444 | 61/20304 | 0,002126 | 0,036578 | ENSDARG00000102776/ENSDARG00000105263/ENSDARG00000039351/ENSDARG00000094002/ENSDARG00000099401/ENSDARG00000102945 | 1,503634 |
| *GO:1990869* | cellular response to chemokine | 6/444 | 61/20304 | 0,002126 | 0,036578 | ENSDARG00000102776/ENSDARG00000105263/ENSDARG00000039351/ENSDARG00000094002/ENSDARG00000099401/ENSDARG00000102945 | 1,503634 |
| *GO:0006954* | inflammatory response | 13/444 | 237/20304 | 0,002258 | 0,038428 | ENSDARG00000102776/ENSDARG00000105263/ENSDARG00000026726/ENSDARG00000058045/ENSDARG00000026663/ENSDARG00000102986/ENSDARG00000076306/ENSDARG00000096645/ENSDARG00000089383/ENSDARG00000039351/ENSDARG00000094002/ENSDARG00000099401/ENSDARG00000102945 | 0,919638 |
| *GO:0034587* | piRNA metabolic process | 3/444 | 13/20304 | 0,002523 | 0,04204 | ENSDARG00000059951/ENSDARG00000041699/ENSDARG00000007465 | 2,356412 |
| *GO:1901072* | glucosamine-containing compound catabolic process | 3/444 | 13/20304 | 0,002523 | 0,04204 | ENSDARG00000093193/ENSDARG00000100635/ENSDARG00000099185 | 2,356412 |
| *GO:0051923* | sulfation | 4/444 | 27/20304 | 0,002657 | 0,043813 | ENSDARG00000041540/ENSDARG00000018361/ENSDARG00000006811/ENSDARG00000071495 | 1,913206 |
| *GO:0009308* | amine metabolic process | 6/444 | 64/20304 | 0,002717 | 0,044348 | ENSDARG00000035652/ENSDARG00000041540/ENSDARG00000018361/ENSDARG00000071429/ENSDARG00000023176/ENSDARG00000086458 | 1,455625 |
| *GO:1901606* | alpha-amino acid catabolic process | 7/444 | 86/20304 | 0,002777 | 0,044859 | ENSDARG00000018351/ENSDARG00000044935/ENSDARG00000071429/ENSDARG00000039269/ENSDARG00000069630/ENSDARG00000023176/ENSDARG00000058005 | 1,314311 |

420

**Supplementary Table 7. Comparison of most prominent symptoms in Loeys-Dietz syndrome type 3, *SMAD6*-related disease and 15q22.31q23 microdeletion syndrome.** 15q22.31q23 syndrome shows an attenuated cardiovascular phenotype compared to Loeys-Dietz syndrome type 3. Legend: [a] only mild aortic root dilatation (identified in the proband's mother); [b] downslanted palpebral fissures, hypertelorism, prominent nose with a wide nasal bridge, and large protruding ears; [c] hypertelorism; [d] only mild scoliosis, no clinical relevance.

| | 15q22.31q23 microdeletion syndrome | Loeys-Dietz syndrome type 3 | *SMAD6*-related disease |
|---|---|---|---|
| **Affected genes** | Including *SMAD3*, *SMAD6*, and *AAGAB* | *SMAD3* | *SMAD6* |
| **Clinical characteristics** | Present case | van de Laar et al. | Luyckx et al. |
| TAAD | +[a] | + | + |
| Arterial tortuosity | + | + | - |
| Bicuspid aortic valve | + | - | + |
| Mitral valve prolapse | - | + | - |
| Other congenital heart diseases | - | + | + |
| Craniofacial dysmorphism | +[b] | +[c] | +/- |
| Craniosynostosis | - | - | + |
| High arched plate | + | + | - |
| Thin translucent skin | + | + | - |
| Umbilical/inguinal hernia | - | + | - |
| Radio-ulnar dysplasia | - | - | + |
| Arachnodactyly | + | + | - |
| Pectus deformity | + | + | - |
| Scoliosis | +/-[d] | + | - |
| Hypermobile joints | + | + | - |
| Osteoarthritis | - | + | - |
| Hyperkeratosis | + | - | - |

### SUPPLEMENTARY MOVIES

**Supplementary movie 1: synchrotron imaging of qKO zebrafish.** 5x imaging of the ventral aorta and ventricle in qKO zebrafish at 8 mpf. Aorta damage can be observed between aortic arch 1 and 2. Left aortic arch 2, right aortic arch 3 and both aortic arch 4 branches appear to be missing. Cross section images are taken starting at the snout, moving caudally. Legend: “BA” bulbus arteriosus, “V” ventricle, “A” atrium. Black arrow indicates the ventral aorta.

**Supplementary movie 2: synchrotron imaging of WT control.** 5x imaging of the ventral aorta and ventricle in WT zebrafish at 8 mpf. Cross section images are taken starting at the snout, moving caudally. Legend: “BA” bulbus arteriosus, “V” ventricle, “A” atrium. Black arrow indicates the ventral aorta.

**Supplementary movie 3: synchrotron imaging of qKO zebrafish between aortic arch 1 and 2.** 20x imaging of the ventral aorta in qKO zebrafish at 8 mpf. Aorta damage can be observed between aortic arch 1 and 2. Asymmetrical branching is visible. Only the right aortic arch 2 is present. Cross section images are taken starting at the snout, moving caudally. Black arrow indicates the ventral aorta.

**Supplementary movie 4: synchrotron imaging of WT zebrafish between aortic arch 1 and 2.** 20x imaging of the ventral aorta in WT zebrafish at 8 mpf. Branching of aortic arch 2 is symmetrical. Left and right branch are present. No damaged region in de ventral aorta can be observed. Cross section images are taken starting at the snout, moving caudally. Black arrow indicates the ventral aorta.

**Supplementary movie 5: synchrotron imaging of qKO zebrafish between aortic arch 2 and 3.** 20x imaging of the ventral aorta in qKO zebrafish at 8 mpf. Aorta damage can be observed between aortic arch 2 and 3. Asymmetrical branching of aortic arch is visible. Diameter of right aortic arch 3 is smaller compared to the diameter of left aortic arch 3. Aorta damage is visible anterior of aortic arch 3. Cross section images are taken starting at the snout, moving caudally. Black arrow indicates the ventral aorta. Red arrow indicates aorta damage.

**Supplementary movie 6: synchrotron imaging of WT zebrafish between aortic arch 2 and 3.** 20x imaging of the ventral aorta in WT zebrafish at 8 mpf. Cross section images are taken starting at the snout, moving caudally. Black arrow indicates the ventral aorta.

**Supplementary movie 7: overview synchrotron image of WT zebrafish**. 5x magnification overview of the full body scan of an 8 mpf WT zebrafish.

**Supplementary movie 8: overview synchrotron image of qKO zebrafish**. 5x magnification overview of the full body scan of an 8 mpf qKO zebrafish.

**Supplementary movie 9: 3D reconstruction of ventral aorta and aortic arches in adult qKO**. Aortic arch two is connected twice at the same side of the aorta. Aortic arch

3 and 4 show a severe reduction in diameter. Region with aorta damage is indicated in red. Regions in which sections were lost are left blank.

**Supplementary movie 10: 3D reconstruction of ventral aorta and aortic arches in adult qKO**. Aortic arch 2 is only present at the right side. A reduction of diameter of the left aortic arches 3 and 4 can be observed. Region with damage is indicated in pink.

**Supplementary movie 11: 3D reconstruction of ventral aorta and aortic arches in adult qKO.** Right aortic arch 2 and 4 are reduced in diameter compared to their left counterpart. Region with aorta damage has been indicated in blue. Rupture is indicated in yellow.

**Supplementary movie 12: 3D reconstruction of ventral aorta and aortic arches in adult qKO.** Right aortic arch 2 is severely reduced in diameter compared to the left counterpart. Region with aorta damage is indicated in blue. Rupture is indicated in yellow.

**Supplementary movie 13: 3D reconstruction of ventral aorta and aortic arches in adult qKO.** Right aortic arch 4 is severely reduced in diameter. Right aortic arch 2 and 3 are missing. Region with damage is indicated in yellow.

**Supplementary movie 14: 3D reconstruction of ventral aorta and aortic arches in adult WT controls**. Full 3D reconstruction of zebrafish 1. The ventral aorta shows symmetrical branching and aortic arches similar in diameter.

**Supplementary movie 15: 3D reconstruction of ventral aorta and aortic arches in adult WT controls**. Full 3D reconstruction of zebrafish 2. The ventral aorta shows symmetrical branching and aortic arches similar in diameter.

**Supplementary movie 16: 3D reconstruction of ventral aorta and aortic arches in adult WT controls**. Full 3D reconstruction of zebrafish 3. The ventral aorta shows symmetrical branching and aortic arches similar in diameter.

**Supplementary movie 17: 3D reconstruction of ventral aorta and aortic arches in adult *smad3a$^{-/-}$;smad3b$^{-/-}$* DKO show no clear abnormalities**. Full 3D reconstruction of zebrafish 1. The ventral aorta shows symmetrical branching and aortic arches similar in diameter.

**Supplementary movie 18: 3D reconstruction of ventral aorta and aortic arches in adult *smad3a$^{-/-}$;smad3b$^{-/-}$* DKO show no clear abnormalities**. Full 3D reconstruction of zebrafish 2: The ventral aorta shows symmetrical branching and aortic arches similar in diameter.

**Supplementary movie 19: 3D reconstruction of ventral aorta and aortic arches in adult *smad3a$^{-/-}$;smad3b$^{-/-}$* DKO show no clear abnormalities**. Full 3D reconstruction of zebrafish 3. The ventral aorta shows symmetrical branching and aortic arches similar in diameter.
